# Genome-wide characterization of human minisatellite VNTRs: population-specific alleles and gene expression differences

**DOI:** 10.1101/2020.11.03.367367

**Authors:** Marzieh Eslami Rasekh, Yozen Hernandez, Samantha D. Drinan, Juan I. Fuxman Bass, Gary Benson

## Abstract

Variable Number Tandem Repeats (VNTRs) are tandem repeat (TR) loci that vary in copy number across a population. Using our program, VNTRseek, we analyzed human whole genome sequencing datasets from 2,770 individuals in order to detect minisatellite VNTRs, i.e., those with pattern sizes ≥7 bp. We detected 35,638 VNTR loci and classified 5,676 as commonly polymorphic (i.e., with non-reference alleles occurring in >5% of the population). Commonly polymorphic VNTR loci were found to be enriched in genomic regions with regulatory function, i.e., transcription start sites and enhancers. Investigation of the commonly polymorphic VNTRs in the context of population ancestry revealed that 1,096 loci contained population-specific alleles and that those could be used to classify individuals into super-populations with near-perfect accuracy. Search for quantitative trait loci (eQTLs), among the VNTRs proximal to genes, indicated that in 187 genes expression differences correlated with VNTR genotype. We validated our predictions in several ways, including experimentally, through the identification of predicted alleles in long reads, and by comparisons showing consistency between sequencing platforms. This study is the most comprehensive analysis of minisatellite VNTRs in the human population to date.

## INTRODUCTION

Over 50% of the human genome consists of repetitive DNA sequence (1, 2) and tandem repeats (TRs) comprise one such class. A TR consists of a pattern of nucleotides repeated two or more times in succession. TR loci are defined by their position on the genome, the sequence and length of their repeat unit, and copy number.

TRs are commonly divided into three classes based on pattern size: short tandem repeats (STRs), or microsatellites, with pattern lengths of six or fewer base pairs (bp), minisatellites with patterns ranging from seven to several hundred bp, and macrosatellites with patterns from hundreds to thousands of bp (3, 4). This study focuses on the TR minisatellite class, which comprises more than a million loci in the human genome.

Many TRs appear to be monoallelic with regard to copy number. However a significant fraction exhibit copy number variability in the population and these variant TR loci are called Variable Number Tandem Repeats (VNTRs). Changes in VNTR copy number have been proposed to arise by slipped strand mispairing (5, 6, 7), unequal crossover (8, 9), and gene conversion (8, 10).

VNTRs are highly mutable, with germline mutation rates estimated between 10^−3^ and 10^−7^ per cell division (11, 12,13,14,15). This mutation rate, which far exceeds that of SNPs, makes VNTRs useful for DNA fingerprinting (16, 17, 18). VNTRs have also been predicted to have high heterozygosity, ranging from 43% to 59% (19), and the copy numbers of several VNTR loci have been shown to be population-specific in humans (20, 21), suggesting that these VNTRs may be useful for population wide studies.

More than half of previously identified human VNTR loci (22) are located near or within genes (23), and so their potential effects on gene expression or protein products are substantial. Indeed VNTRs have been associated with changes in levels of gene expression (24), including tissue specific expression (25, 26).

Minisatellite VNTRs have been proposed to regulate genes in several ways. In promoters, they can carry binding sites for transcription factors such as NF-*κ*B and myc/HLH (27, 28), permitting copy number to affect transcription factor binding and therefore level of transcription (29, 30, 31, 32). VNTRs in introns have been shown to contain enhancer sequences (33, 34, 35) or to cause differential splicing (36, 37). VNTRs in exons can also affect transcription (38, 39) and the stability or translation rate of the resulting proteins (40).

Microsatellites have been extensively studied (41,42,43,44,45) and linked to various diseases and cancer (46, 47, 48, 49). For example, in Ewing sarcoma, the EWS-FLI fusion protein activation of enhancer regions at GGAA microsatellite repeats contributes to tumor development (50, 51, 52) and repression of these enhancers impairs tumor growth (51).

Furthermore, minisatellite VNTRs have been associated with a variety of diseases (53, 54, 55), including neurodegenerative disorders such as Alzheimer’s disease (36, 56, 57) and Huntington’s disease (28, 58), and other psychiatric conditions, such as PTSD (59), ADHD (60, 61), depression (62), and addiction (63). VNTRs have been shown to be risk factors in various cancers (64,65,66,67,68,69,70,71,72,73,74) and have been linked to cancer prognosis and outcome (75,76,77,78,79,80). Commercial cancer diagnosis kits using minisatellite VNTRs have been introduced (81) and it has been proposed that VNTRs associated with cancers be used for targeted sequencing in personalized therapies (82, 83, 84).

VNTRs have been proposed as drivers of phenotypic variation in evolution (85, 86, 87). For example, an examination of functional enrichment in genes containing TRs or VNTRs, in both humans and apes, found that genes with ape-specific TR variability were associated with the senses of taste and smell, while genes with human-specific TR variability or human-specific copy number were associated with neurogenesis and neural development. These results suggest that VNTR polymorphisms may help account for “human-specific cognitive traits” (24). Additionally, the Eichler group (87) has examined TR loci on human and ape genome assemblies from PacBio sequencing data and identified 1,584 human-specific VNTR loci with 52 as candidate regions associated with disease.

Despite their biological significance, until recently, relatively few human minisatellite VNTRs have been identified and studied in detail. The ever–increasing availability of accurate whole genome sequencing (WGS) data, however, provides extensive opportunity for high throughput, genome-wide VNTR genotyping. Further, the emergence of PCR-free WGS datasets is reducing locus selection bias and enabling better filtering of false positive VNTR variants.

Nonetheless, genotyping variability in repeat sites remains challenging (88, 89). Although a number of tools have been designed to detect microsatellite copy number variability such as lobSTR (90), popSTR (91), hipSTR (92), GangSTR (93), and ExpansionHunter (94), only two high-throughput tools are available for minisatellite genotyping. The adVNTR tool (95) trains a Hidden Markov Model (HMM) for each VNTR locus of interest and has been used to predict variability in 2,944 VNTRs intersecting coding regions.

VNTRseek (96), developed in our lab, uses the Tandem Repeats Finder (TRF) (97) to detect and characterize TRs inside reads and then maps read TRs to TRs in a reference set. Because it builds pattern profiles before mapping, VNTRseek is robust in the presence of SNPs and small indels. While VNTRseek has high precision, it has two major limitations. It only detects minisatellites which contain a minimum of 1.8 to 1.9 pattern copies (depending on pattern length), a limitation from the Tandem Repeats Finder (TRF) software, and the tandem array, plus short flanking sequences, must fit completely inside a read. This means that arrays longer than the read length cannot be detected. Given these limitations, the results we report underestimate the true polymorphic nature of minisatellite TRs.

In this paper, we present the most comprehensive catalog of minisatellite VNTRs in the human genome to date, pooling results for WGS datasets from 2,770 individuals, processed with VNTRseek on the GRCh38 human reference genome. We report a large collection of previously unknown VNTR loci, find that many VNTR loci and alleles are common in the population, show that VNTR loci are enriched in gene and regulatory sequences, provide evidence of gene expression differences correlated with VNTR genotype and evidence of population-specific VNTR alleles.

## MATERIALS AND METHODS

### Datasets

Datasets comprising 2,801 PCR-free, WGS samples from 2,770 individuals were used in this study (Table 1): 30 individuals from the 1000 Genomes Project Phase 3 (98), including the Utah (CEU) and Yoruban (YRI) trios (mother-father-child); 2,504 unrelated individuals mostly overlapping with the 1000 Genomes Project, recently sequenced at *>*30 *×* coverage by the New York Genome Center (NYGC); 253 individuals from the Simons Genome Diversity Project (SGDP) (43), seven individuals sequenced by the Genome in a Bottle (GIAB) Consortium (99), including the Chinese (HAN) and Ashkenazi Jewish (AJ) trios and NA12878 (with ID HG001); two “haploid” hydatidiform mole cell line genomes, CHM1 (100) and CHM13 (101); tumor/normal tissues from two unrelated individuals with breast cancer (breast invasive ductal carcinoma cell line/lymphoblastoid cell line) from the Illumina Basespace public WGS datasets (102); and the AJ child sequenced with PacBio Circular Consensus Sequencing (CCS) reads (103). Duplicates of 27 genomes were present in two datasets, 1000 Genomes and NYGC. One of these, NA12878, was also included in the GIAB dataset.

**Table 1.**
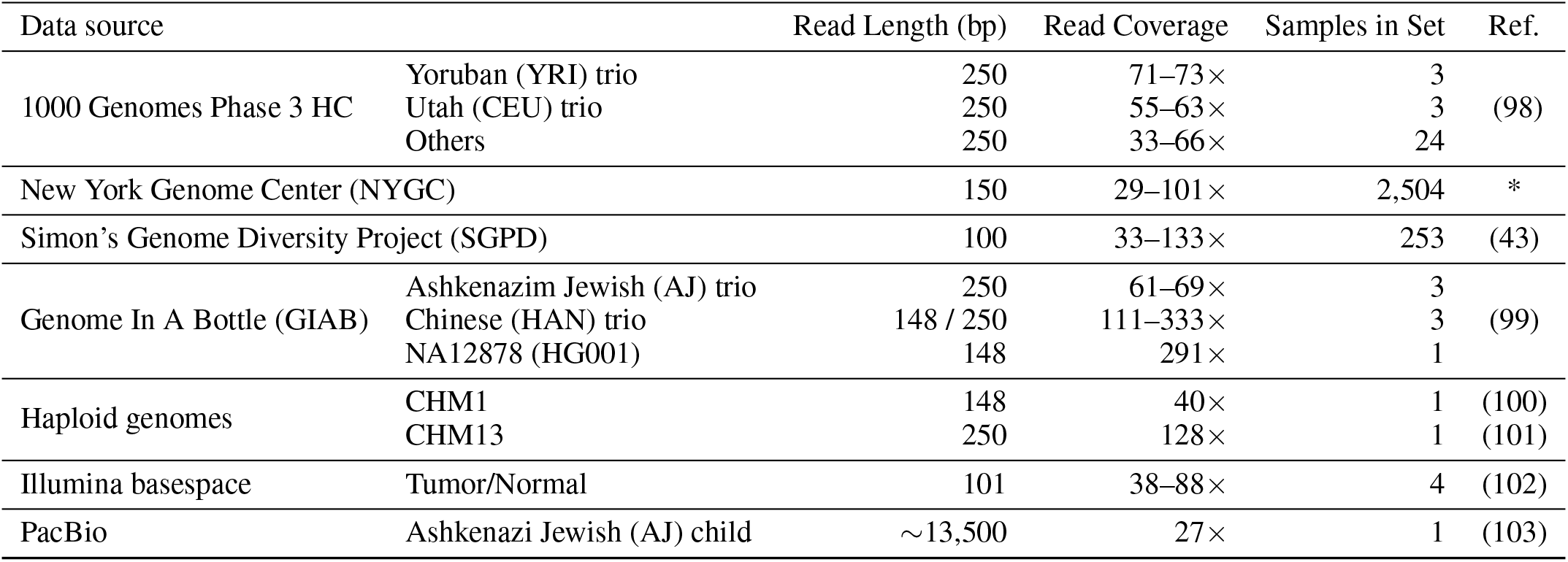
Data. 2,801 publicly available WGS samples, for 2,770 individuals, were used in this study. Read coverage was calculated as the product of the number of reads and the average read length, divided by the haploid genome size, as in the Lander/Waterman equation (104). All coverage values are approximate. The 1000 Genomes Phase 3 samples were released in 2015. The NYGC samples were released in 2020 by the New York Genome Center (NYGC). For the Simons Genome Diversity Project (SGDP), released in 2016, only datasets which were not present in the the 1000 Genomes datasets were used. The PacBio data were used only for comparison and validation purposes but not for our VNTR results. *These data were generated at the New York Genome Center with funds provided by NHGRI Grant 3UM1HG008901-03S1.

Overall, read coverage ranged from approximately 27x, in the PacBio sample to 333x, in the GIAB Chinese child. Besides the PacBio data, reads consisted of three lengths, 100/101 bp (257 samples), 148/150 bp (2,508 samples), and 250 bp (35 samples). All data were downloaded as raw fastq files, except for the PacBio data which were obtained as a BAM file with reads aligned to GRCh37. SRA links to the data are given in Table 4.

The majority of the analyses in this paper were performed on the 2,504 genomes from NYGC. The 253 genomes from SGDP provided insight into under-represented populations. The 27 genomes duplicated in the 1000 Genomes and NYGC datasets were used to measure consistency across sequencing platforms. The trios from the 1000 Genomes (CEU and YRI) and GIAB (AJ and Chinese HAN) datasets were used for analyzing Mendelian inheritance. The cancer datasets were used to find possible changes in VNTRs in tumor tissues. The PacBio data were used for validation purposes only.

### Curating the reference TRs

The 22 autosomes and sex chromosomes from the human reference genome GRCh38 were used to produce a reference set of TRs in TRDB (105) with the TRF software and four quality filtering steps as described in (96). In addition, centromere regions were excluded from the reference set. These filtering tools are available online in TRDB. Starting with 1,199,362 TRs found by TRF, we curated a filtered reference set with 228,486 TRs. Using VNTRseek, we classified the TRs into two subcategories, singletons and indistinguishables (Supplementary Section S2.1). A *singleton* TR appears to be unique in the genome based on a combination of its repeat pattern and flanking sequence. An *indistinguishable* TR belongs to a family of genomically dispersed TRs which share highly similar patterns and flanking sequence and may therefore produce misleading genotype calls. Indistinguishable TRs (total 37,200 or about 16% of the reference set) were flagged. Simulation testing revealed that some singletons produced false positive VNTRs. To minimize this issue, an additional filtering step was added to eliminate problematic singleton loci from the reference set (see Supplementary Section S2.2 and Supplementary Material `Reference TRs.txt` for the reference sets).

We assumed that genotyping was possible for a reference TR locus, given a particular read length, if the TR array length plus a minimum 10 bp flank on each side, would fit within the read. The number of reference TR alleles that could be genotyped using each of the read lengths in our data is summarized in Supplementary Table S1.

### Genotyping TRs and VNTRs

Each dataset was processed separately with VNTRseek using default parameters: minimum and maximum flanking sequence lengths of 10 bp and 50 bp, respectively, on each side of the array, and requiring at least two reads mapped with the same array copy number to make an allele call. Output from VNTRseek included two VCF files containing genotype calls, one reporting all detected TR and VNTR loci, and the other limited to VNTR loci only. (A locus was considered a VNTR if a non-reference allele was detected.) The VCF files contained two specialized FORMAT fields: SP, for number of reads *supporting* each allele, and CGL, to indicate genotype by the number of *copies gained or lost* with respect to the reference. For example, a genotype of 0 indicated detection of only the TR reference allele (zero copies gained or lost), while 0,+2 indicated a heterozygous locus with a reference allele and an allele with a gain of two copies.

To remove clear inconsistencies, for this study we filtered the VCF files to remove *per sample* VNTR loci with more alleles than the expected number of chromosomes. The filtering criteria for these loci, termed *multis* is detailed in Supplementary Section S2.3. After multi filtering, a TR locus was labeled as a VNTR if any remaining allele, different from the reference, was observed in any sample.

### Experimental validation

Accuracy of VNTRseek genotyping was experimentally tested for 13 predicted VNTR loci in the Ashkenazi Jewish (AJ) trio. The following DNA samples were obtained from the NIGMS Human Genetic Cell Repository at the Coriell Institute for Medical Research: NA24385, NA24149, and NA24143 (also identified as GIAB IDs HG002, HG003, and HG004). Selection criteria required that the PCR product was not contained in a repeat region, unique primers could be designed, the primer-defined allele length difference was between 10% and 20% of the longest allele, and the primer-defined GC content was between 40% and 60%. Given these criteria, we prioritized VNTRs in genes and regulatory regions which might be of interest to researchers. Primers were designed with Primer-BLAST (106) and used to amplify the VNTR loci from the genomic DNA of each individual using the following reagents: 0.2 *µ*L 5 U/*µ*L DreamTaq DNA Polymerase (ThermoFisher Scientific), 4.0 *µ*L 10X DreamTaq Buffer (ThermoFisher Scientific), 0.8 *µ*L 10mM dNTP mix (ThermoFisher Scientific), 3.2 *µ*L primer mix at a final concentration of 0.5 *µ*M, 1.6 *µ*L genomic DNA (40 ng), and 30.2 *µ*L nuclease-free water. PCR cycling conditions were as follows: 30 seconds at 95*°*C, 30 seconds at 56-60*°*C, 20 seconds at 72*°*C, for 30 cycles, with an initial denaturation of 3 min at 95*°*C and a final extension of 7 min at 72*°*C. The resulting amplicons were run on a 2% agarose gel at 100 V for 2 hours and visualized with UV light using Ethidium Bromide. A complete list of loci and primers is given in the Supplementary Material as `Experiment primers.txt`.

In addition, VNTRseek predictions in the NA12878 genome were compared to experimental validations in the paper describing the adVNTR software (95). We had three datasets for NA12878; HG001 (148 bp) from GIAB, NA12878 (150 bp) from NYGC, and NA12878 (250 bp) from 1000 Genomes. The adVNTR predictions used GRCh37 coordinates which were converted using the UCSC liftover tool (107) to coordinates to GRCh38.

### Validation using long reads

Aligned PacBio reads for the AJ child (GIAB ID HG002) were processed to validate VNTRseek predictions. The read sequences were extracted from the BAM file and mapped back to the GRCh38 genome using BWA MEM default settings (108). Using bedtools (109), the reads aligning to each TR reference locus were extracted. For each read, a local wraparound dynamic programming alignment was performed using the reference pattern and the same scoring parameters used to generate the reference set (match=+2, mismatch= − 5, and gap= −7). The number of copies of the pattern in the resulting alignment was then determined and compared with the VNTRseek predictions. If the difference between a PacBio copy number in at least one read and the VNTRseek copy number was within *±* 0.25 of a copy, we considered the VNTRseek allele to be validated.

### Measuring consistency of Mendelian inheritance

A locus on an autosomal chromosome is consistent with Mendelian inheritance if the genotype of a child can be explained as one allele from the mother and one from the father. Genotype consistency was evaluated for all mother-father-child trios, i.e., the AJ, CEU, HAN, and YRI trios. We evaluated loci defined by several increasingly stringent criteria: both parents heterozygous, all members of the trio heterozygous, all members of the trio heterozygous and with different genotypes. These criteria were selected to avoid false interpretations of consistency.

TR loci on the X and Y chromosome of male children were also selected for evaluation when both the son and the appropriate parent had a predicted genotype. In these cases, inheritance consistency means a son’s X chromosome allele is observed on one of the mother’s X chromosomes, and a son’s Y chromosome allele is observed on the father’s Y chromosome.

### Measuring allele consistency across platforms

VNTR calls were compared for each of 27 genomes that were represented twice, once in the 1000 Genomes dataset, sequenced in 2015 on an Illumina HiSeq2500 with 250 bp read length and once in the NYGC dataset, sequenced in 2019 on an Illumina NovaSeq 6000 with 150 bp read length. The two platforms have different error profiles.

Because read length and coverage differed among datasets, for each pairwise comparison, we only considered loci that were genotyped in both samples and classified as VNTR in at least one. We extracted the *non-reference* VNTR alleles (detected in at least one sample) and computed consistency as the ratio of the size of the intersection set of those alleles (found in both platforms) over the size of their union (found in either). For alleles detected in the 250 bp reads, we only counted those that could have been detected in the shorter 150 bp reads. Reference alleles were excluded to avoid inflating the ratio.

### Common and private VNTRs

To classify commonly polymorphic VNTRs (hereafter “common VNTRs”) and private VNTRs, we used results from the NYGC dataset (2,504 individuals) as the read length and coverage were comparable across all genomes. Additionally, these genomes contain no related individuals and represent a wide set of populations (26 populations from five continents). Loci were classified as common VNTRs if non-reference alleles were identified in at least 5% (126) of the individuals and classified as private VNTRs if non-reference alleles were identified in less than 1% (25).

### Annotation and enrichment

Annotation based on overlap with functional genomic regions was performed for the reference TR loci. Genomic annotations for GRCh38 were obtained from the UCSC Table Browser (110) in BED format. Known gene transcripts from GENCODE V32 (111) were used along with tracks for introns, coding exons, and 5^*′*^and 3^*′*^exons. Regulatory annotations included transcription factor binding site (TFBS) clusters (112, 113) and DNAse clusters (114) from ENCODE 3 (115), and CpG island tracks (116), comprising 25%, 15%, and 1% of the genome, respectively. Bedtools (109) was used to find overlaps between TR loci and the annotation features. Any size overlap was allowed.

LOLAweb (117) was used to determine VNTR enrichment for genomic regions in comparison to the background TR annotations, and common and private VNTR enrichment in comparison to all VNTR annotations. TRs on the sex chromosomes were excluded in the background set. To identify gene and pathway functions that could be affected by common VNTR copy number change, genes with exons or introns overlapping with common VNTRs were collected and their enrichment computed using GSEA (118) for Gene Ontology (GO) terms (119) for biological process and KEGG pathways (120) with FDR p-value *≤* 0.05.

### Association of VNTR alleles with gene expression

To detect expression differences among individuals with different VNTR genotypes, mRNA expression counts from lymphoblastoid cell lines of 660 individuals by the Geuvadis consortium (Accession: E-GEUV-1) were downloaded (121). A total of 445 individuals overlapped with the 2,504 NYGC genomes set. We paired VNTR loci with genes within 10 Kbp, and extracted the genotypes for each individual at those VNTRs. When no genotype was observed for an individual, we classified the genotype as *other*. We did this because we assumed that the alleles were outside the detection range, given that genotypes were observed in other individuals with similar coverage. VNTR loci were retained for analysis if at least two genotypes were detected for that VNTR across all individuals (at least three if *other* was one of the genotypes) and if each genotype was observed in at least 20 individuals. Genes were excluded from analysis if the median TPM (Transcripts Per Kilobase Million) expression value equaled zero.

To control for confounders we used covariates for sex and population structure and detected additional hidden covariates using Iteratively Adjusted Surrogate Variable Analysis (IA-SVA) (122) on the log_2_ normalized TPM values. For population structure we used the top five principal components determined from a principal components analysis of the informative SNP genotypes from the 445 individuals as reported by the 1000 Genomes project (http://ftp.1000genomes.ebi.ac.uk/vol1/ftp/release/20130502/supporting/hd_genotype_chip/). Using IA-SVA and observing that covariates sixteen and above were over 85% correlated with other covariates (Supplementary Figures S22–S24), we chose fifteen hidden factors to include in our model. Finally, we used a linear regression *expression ∼ sex*+*population_PCAs*+*hidden factors* with the log_2_ normalized TPM values to extract residuals to be used in the downstream association model.

For each gene-VNTR pair, we used a one-way ANOVA test as *residuals ∼ genotype* to detect if the mean of any genotype class was different from the others. The p-values of the ANOVA tests were adjusted using FDR. Any gene-VNTR pair with FDR*<*0.05 was reported. For significant eQTLs, we calculated, for reporting purposes, the maximum difference of the residual means over all pairs of genotype classes.

To associate eQTLs with histone marks or open chromatin, we downloaded narrow peaks data in GRCh38 in bed format from 14 experiments on histone marks and one on DNAse hypersensitive sites from the GM12878 (B-Lymphocyte) cell line from the ENCODE project (123) (source IDs are given in Supplementary Table S13). Any overlaps of peaks with the eQTL VNTRs were reported.

### Population-specific alleles

The 2,504 genomes in the NYGC dataset consisted of 26 populations of individuals with ancestry from five super-populations: African, American, East Asian, European, and South Asian. To investigate the predictive power of common VNTRs with regard to super-population membership, Principal Component Analysis (PCA) clustering was applied. For each sample, a vector of common loci *alleles* showing presence/absence (1/0) was produced. Uninformative alleles (that were not present in at least 5% of the samples) were removed and principal components (PCs) calculated over the resulting vector set. Using a 70% training to 30% testing split of the data, a decision tree based on the first 10 PCs was trained using 10-fold cross validation and then validated on the testing data. In order to find super-population markers among the common VNTRs, a one-sided Fisher’s exact test was used to calculate the odds ratio and p-value of each allele being in one super-population versus being collectively in all the others. We only considered alleles over-represented rather than both over-and under-represented because of an interest in identifying alleles that have a phenotypic effect. Odds ratio values were log_2_ transformed and p-values were adjusted for false discovery rate (FDR) (124). Any allele with FDR *<* 0.05 and log_2_(odds ratio) *>*1 was chosen as a significant marker for that population.

## RESULTS

In this section, we start with a summary and characterization of our VNTR predictions, followed by identification of commonly polymorphic VNTRs and an enrichment analysis of their association with genomic functional regions and genes sets. We next report on the effect of VNTRs on expression of nearby genes, and then identify population-specific VNTR alleles and show that they are predictive of ancestry. We conclude this section with evidence confirming the accuracy of our predictions using several validation methods.

### About one in five minisatellite TRs are variable in the human population

WGS datasets from 2,770 human genomes were analyzed with VNTRseek to detect VNTRs. Overall, 184,315 out of 191,286 singleton reference TR loci (*∼*96%) were genotyped across all samples (Table 2) while 5% of the loci had TR arrays too long to fit within the longest reads and could only be genotyped if they lost a sufficient number of copies.

**Table 2.**
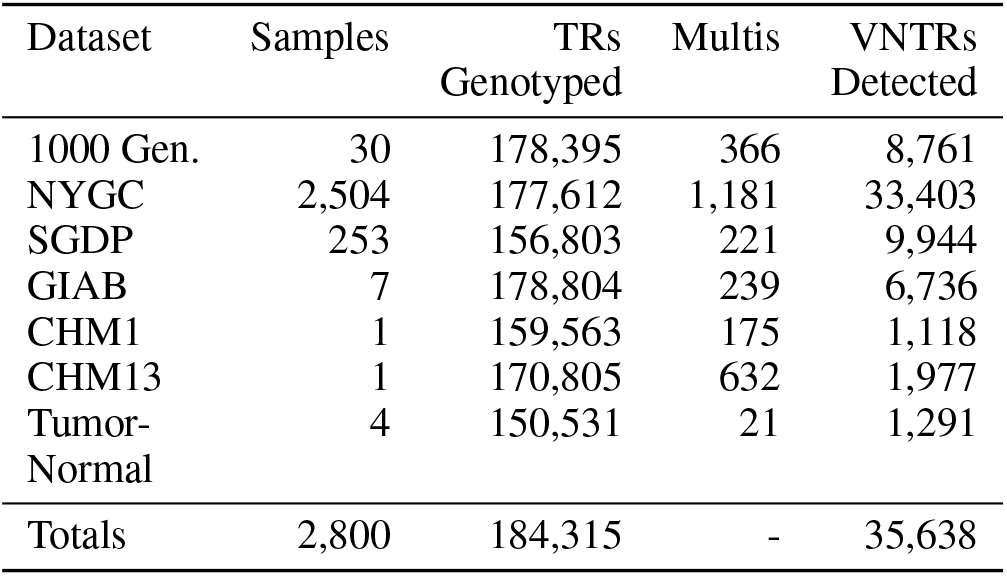
TRs and VNTRs detected, by dataset. *TRs Genotyped* is the number of distinct TR loci genotyped across all individuals within a dataset. (All other numbers are also per dataset.) *Multis* are TR loci genotyped in a single individual with more than the expected number of alleles. They could be artifacts or indicate copy number variation in a genomic segment. Multis were excluded from further analysis on a per sample basis. *VNTRs Detected* is the number of TR loci, excluding multis, with a detected allele different from the reference.

A total of 5,198,392 loci with non-reference alleles were detected, corresponding to 35,638 (*∼* 19%) distinct VNTR loci, indicating wide occurrence of these variable repeats. Their occurrence within genes was common, totaling 7,698 protein coding genes, and 3,512 exons. The resulting genotypes were output in VCF format files (see Data Availability Section) and summarized for each genome (Supplementary Material `Summary of results.txt`). A website is under development to view the VNTR alleles (http://orca.bu.edu/VNTRview/).

### The number of TRs genotyped and VNTRs detected depends on coverage and read length

To determine the effect of coverage and read length on genotyping, we measured two quantities: the percentage of reference singleton TRs that were genotyped, and the total number of singleton VNTRs that were detected in each genome. Only singleton loci were considered in all further analyses. Figure 1a shows that there was a strong positive correlation between coverage and the ability to genotype TRs. A strong correlation with read length was also apparent, however, the effect was larger, primarily due to the ability of longer reads to span, and thus allow VNTRseek to detect, longer TR arrays. These results suggest that our VNTR numbers are undercounts.

**Figure 1.**
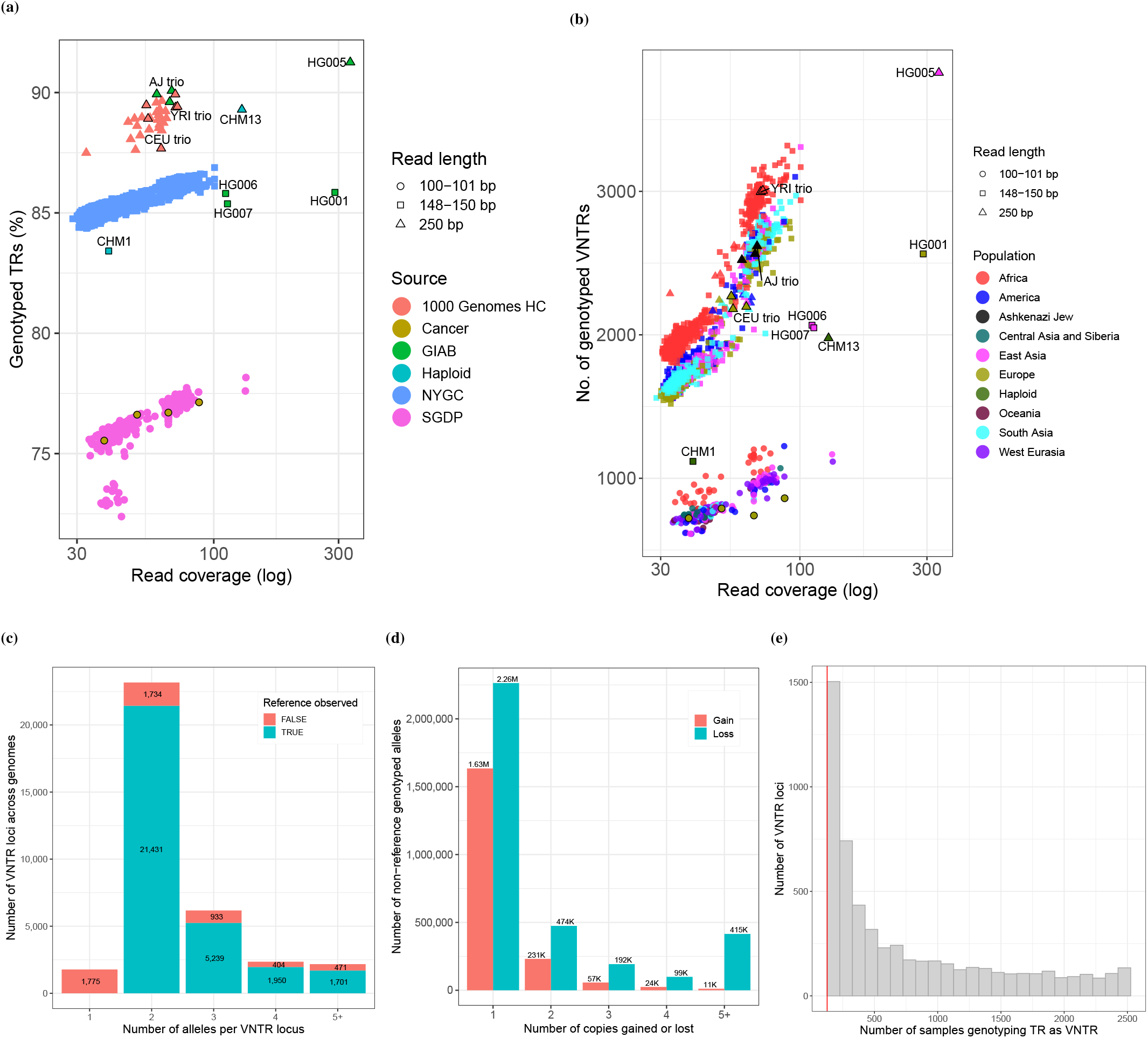
**(a) TR Genotyping sensitivity.** Graph shows the relationship between coverage, read length, and the percentage of TRs in the reference set that were genotyped. Each symbol represents a single sample and specific samples are labeled. Increasing read length had the largest effect on sensitivity because many reference TR alleles could not be detected at the shorter read lengths (see Supplementary Table S1). **(b) VNTRs detected**. Graph shows the relationship between coverage, read length, and the number of VNTR loci detected. Read length and coverage both had large effects. Coloring of symbols shows that population also had a strong effect, reflecting distance from the reference, which is primarily European. Note the reduced numbers for CHM1 (150 bp) and CHM13 (250 bp). Because they are “haploid” genomes, parental heterozygous loci with one reference allele would appear to be VNTRs, on average, only about half the time. **(c) Alleles detected per locus**. Each bar represents a specific number of alleles detected across all datasets. Coloring shows that proportion of loci where the reference allele was or was not observed. **(d) Copies gained or lost**. Each bar represents a specific number of copies gained or lost in non-reference VNTR alleles relative to the reference allele. Loss was always more frequently encountered. **(e) VNTR locus sample support**. Data shown are the *common* loci from the 2,504 sample NYGC dataset. Each bar represents the number of samples calling a locus as a VNTR. Bin size is 100. Bar height is number of loci with that sample support. Red line indicates the 5% cutoff for common loci (126 samples).

VNTR detection was similarly dependent on coverage and read length, as shown in Figure 1b. However, detection was also positively correlated with population, which seemed likely due to the evolutionary distance of populations from the reference genome, which is primarily European (125). For example, in the 250 bp trios with comparable coverage, the African Yoruban genomes (YRI) had the highest number of VNTRs detected, followed by the Ashkenazi Jewish genomes (AJ), and finally, the Utah genomes (CEU). Notably, within each trio, the VNTR counts were similar.

The “haploid” genomes CHM1 (150 bp) and CHM13 (250 bp) had greatly reduced VNTR counts relative to genomes with similar coverage and read length. This was because in these genomes, which consist of two copies of an underlying haploid genome, the single allele represented at any VNTR locus would frequently be a reference allele and so the locus would not be called as a VNTR.

### More than two alleles are common in VNTRs

Two alleles were detected in the majority of VNTR loci across all datasets (Figure 1c). However at 10,698 loci (29%), three or more alleles were detected. In a substantial number of loci (5,395), the reference allele was never seen, but in only 105 of these (2%) was the reference allele in the VNTRseek detectable range for the 150 bp and 100 bp reads, which made up the bulk of our data. Interestingly, in 1,166 loci, the reference allele, although detectable, was not the major allele (Supplementary Material `Major genotypes.txt`).

### Loss of VNTR copies relative to the reference is more common than gain

Overall, VNTRseek found approximately 1.8-fold more alleles with copy losses (3,444,128), with respect to the reference copy number, than gains (1,958,250). Loss of one copy (2,263,608) was the most common type of VNTR polymorphism (Figure 1d). Although there were allele lengths that VNTRseek could not detect, this bias persisted even when restricting the loci to only those where gain and loss could both be observed (Supplementary Figure S5). The overabundance of VNTR copy loss may actually be an underestimate. Since VNTRseek required a read to span a TR array for it to be detected, gain of one copy would have been possible in approximately 68%, 82%, and 92% of loci for samples with read lengths of 100 bp, 150 bp, and 250 bp, respectively. By contrast, the reference locus needed to have a minimum of 2.8 copies for a loss of one copy to be observed by TRF, and only 16% of the reference loci met this criterion. Higher observed copy loss could be explained by a bias in the reference genome towards including higher copy number repeats (126), or by an overall mutational preference for copy loss.

### VNTRs have high heterozygosity

High heterozygosity in human populations suggests higher genetic variability and may have beneficial effects on a range of traits associated with human health and disease (127). Since calculating heterozygosity for VNTRs is not straightforward (because of limitations on discovering alleles, especially within shorter reads), we used the percentage of detected, per-sample heterozygous VNTRs as an estimate for heterozygosity. At read length 250 bp, per-sample heterozygous VNTR loci comprised approximately 46-55% of the total, which is comparable to previous theoretical estimates of 43–59% (19). At shorter read lengths, the bottom of the range extended lower (*∼* 38–57% for 150 bp reads, *∼* 29–51% for 100 bp reads, Supplementary Figure S1), as expected, because longer alleles were undetectable if they did not fit within a single read.

Interestingly, despite the previous comment, within genomes that were comparable in read length and coverage, the fraction of heterozygous loci clustered within populations (Supplementary Figures S2–S4), with African genomes generally having more heterozygous calls and East Asians fewer. This result is consistent with previous findings of population differences in SNP heterozygosity among Yoruban and Ashkenazi Jewish individuals with respect to European individuals (128, 129), and suggests higher genomic diversity among African genomes, as has been previously noted (130).

### Loss of heterozygosity observed in tumor samples

A significant loss of heterozygosity (LOH) was observed in predicted VNTRs of one of the tumor tissues compared to its matching normal tissue (sample ID HC1187). The percentage of heterozygous VNTRs was roughly double in the normal tissue (*∼* 38% vs *∼* 19%) (Supplementary Tables S2 and S3). Extreme loss of heterozygosity in small variants has previously been reported in these samples by Illumina Basespace (131) with the number of heterozygous small variants in HC1187 being four times lower in the tumor tissue compared to the normal. Taken together, these results suggest that VNTR LOH could be linked to tumor progression.

Additionally, in both tumors a large number of loci exhibited loss of both alleles in comparison to the normal tissue (Supplementary Table S2). Given that the coverage for the tumor samples was significantly higher than for the normal tissue, it is unlikely that these observations were due to artifacts. Also, the tumor samples did not show a higher percentage of filtered multi VNTRs (too many alleles) than the normal samples (1.37% and 1.23% in normal tissue vs 1.72% and 1.71% in tumor tissue).

Knowledge of gene associations with somatic tumor mutations (VNTR alleles present in a tumor, but not normal tissue) could be useful as indicators of cancer prognosis and for therapy. In the HC2218 individual, somatic tumor mutations overlapped with lncRNAs (ACO73336.1, AC107959.2, AL355388.2), introns (C3orf67, COX17, DHRS3, DPP6, GAN, PCGF3, RGS12, SLC25A13, SLC6A19, TACR2, TEPP), and promoter regions (TRIM24, DUSP4). Of these, DHRS3, DUSP4, GAN, RGS12, and TRIM14 are known oncogenes or tumor suppressors (as indicated by CancerMine (132)). TRIM24 has been associated with prognosis in breast cancer (133, 134, 135) and over-expression of DUSP4 has been shown to improve the outcome of chemotherapy and overall survival (136, 137).

In the HC1187 individual, somatic tumor mutations overlapped with lncRNAs (LINC01708, AC1058290.1, AC104596.1), exons (THNSL2), introns (AJAP1, SMAD1, FLT4, PTPN3, ADAMTSL2, ANO2, SOX5, SGCG, WDR72, NQO1, CCDC200, ARHGAP45, AC005258.1, PEAK3) and promoters (HFM1, TBK1, GNS, LEMD3, FGFR3, VIPR2). Of these, AJAP1, FGFR3, GNS, NQO1, PTPN3, SMAD1, SOX5, and TBK1 are known oncogenes or tumor suppressors (132). TBK1 and FGFR3 have been used as treatment targets for HER2+ breast cancer (138, 139).

### Common vs. private VNTRs

Following methodology used with SNPs (140), we classified VNTR loci in the 2,504 healthy, unrelated individuals from the NYGC dataset (150 bp and coverage *>*30x) as commonly polymorphic, or “common VNTRs,” if non-reference alleles occurred in at least 5% of a population (126 individuals) and as private VNTRs if they occurred in less than 1% (25 individuals).

We classified 5,676 VNTRs as common (17% of the 33,403 VNTRs detected in this population) and 68% as private. In each sample, we detected, on average, 1,951 VNTR loci, and among those, 1,783 were common VNTRs (median 1,677) and 46 were private VNTRs (median 17). A total of 3,627 common VNTRs overlapped with 2,173 protein coding genes including 254 exons. Interestingly, increasing the threshold for common VNTRs did not reduce the number dramatically (Supplementary Figure S6), suggesting that these VNTRs have not occurred randomly, but rather have undergone natural selection. Widespread occurrence of common VNTRs indicates a fitness for use in Genome Wide Association Studies (GWAS). A list of common and private VNTRs can be found in Supplementary Material `Common VNTRs.txt` and `Private VNTRs.txt`.

### Common VNTR enrichment in functionally annotated regions

To determine possible functional effects of the common VNTRs, we classified the overlap of reference TRs with various functionally annotated genomic regions: upstream and downstream of genes, 3^*′*^UTRs, 5^*′*^UTRs, introns, exons, transcription factor binding site (TFBS) clusters, CpG islands, and DNAse clusters (Supplementary Material `Reference set.txt`).

Our reference TR set comprised only 0.52% of the genome, however, 49% of human genes contained at least one TR and 5% of all the TFBS clusters overlapped with TRs. Moreover, high proportions of our TR reference set and common VNTRs intersected with genes (63% and 64% respectively), TFBS clusters (38% and 51%), and DNAse clusters (21% and 28%) (Supplementary Table S5).

In comparison to TRs, VNTR loci were positively enriched in 1 Kbp upstream and downstream regions of genes, 5^*′*^and 3^*′*^UTRs, coding exons, TFBS clusters, DNAse clusters, and CpG islands (p-values*<*0.05) (Supplementary Tables S4 and S5). The common VNTRs, on the other hand, compared to all VNTRs, were enriched in 1 Kbp upstream regions of genes, TFBS, and CpG islands, suggesting regulatory function. Private VNTRs were less likely to occur in 1 Kbp upstream or downstream regions, inside TFBS clusters, open DNAse clusters, or CpG islands.

Focusing on the common VNTRs, we used the LOLAweb (141) online tool to perform enrichment analysis with various curated feature sets (Supplementary Figures S7–S12 in Supplementary Section S5.2), Among the results, DNAse enrichments by tissue type (142) pointed to brain, muscle, epithelial, fibroblast, bone, hematopoietic, cervix, skin, and endothelial (Figure S11) with brain showing up multiple times, consistent with findings in the literature (24). These results suggest that VNTR alleles may affect gene regulation in multiple tissues.

### VNTR genotypes are correlated with gene expression differences

To detect association between VNTR genotypes and expression of nearby genes, we paired VNTRs to any gene within 10 Kbp and after removing genes with low expression and controlling for confounders, applied a one-way ANOVA test to determine if there was a significant difference between the average gene expression levels for the VNTR genotypes. A total of 1,071 gene-VNTR pairs were tested and 193 pairs (187 genes, 188 VNTRs) exhibited significant expression differences at FDR*<*0.05 (Supplementary Figure S25). The top 10 genes by mean expression difference were: DPYSL4, KLF11, B4GALNT3, PIP5K1B, DNAJA4, THNSL2, CD151, MXRA7, HEBP1, and FARP1. Of these eQTL VNTRs, 47 also exhibited population-specific alleles (next section and Supplementary Table S15) and 81 overlapped with peaks for histone marks and DNAse hypersensitive sites (see Supplementary Table S13 and Supplementary Figure S31). There was significant enrichment of eQTL VNTRs were histone marks at p-value *<*5% (Supplementary Table S14).

Three of the top genes are shown in Figure 2. Gene MXRA7 is associated with a VNTR (id 182606303) in the 5’ UTR exon, DPYSL4 is associated with a VNTR (id 182316137) in the first intron, and CSTB is associated with an upstream VNTR (id 182814480). The VNTR region in MXRA7 is a target site for transcription factors METTL23 and JMJD6. METTL23 is known to function as a regulator in the transcriptional pathway for human cognition (143) and has been associated with mental retardation and intellectual disability (144). JMJD6 is associated with pancreatitis (145) and tumorgenesis (146, 147). Copy number expansions in the VNTR upstream of CSTB have been previously associated with progressive myoclonic epilepsy (EPM1) (148). For this VNTR, we observed the -1 and 0 alleles (2 and 3 copies, respectively), which are common in healthy individuals. However, 201 individuals had genotypes outside of our detection range which likely represented longer expansions and these individuals showed higher expression of this gene. More examples are given in Supplementary Figures S26–S30.

**Figure 2.**
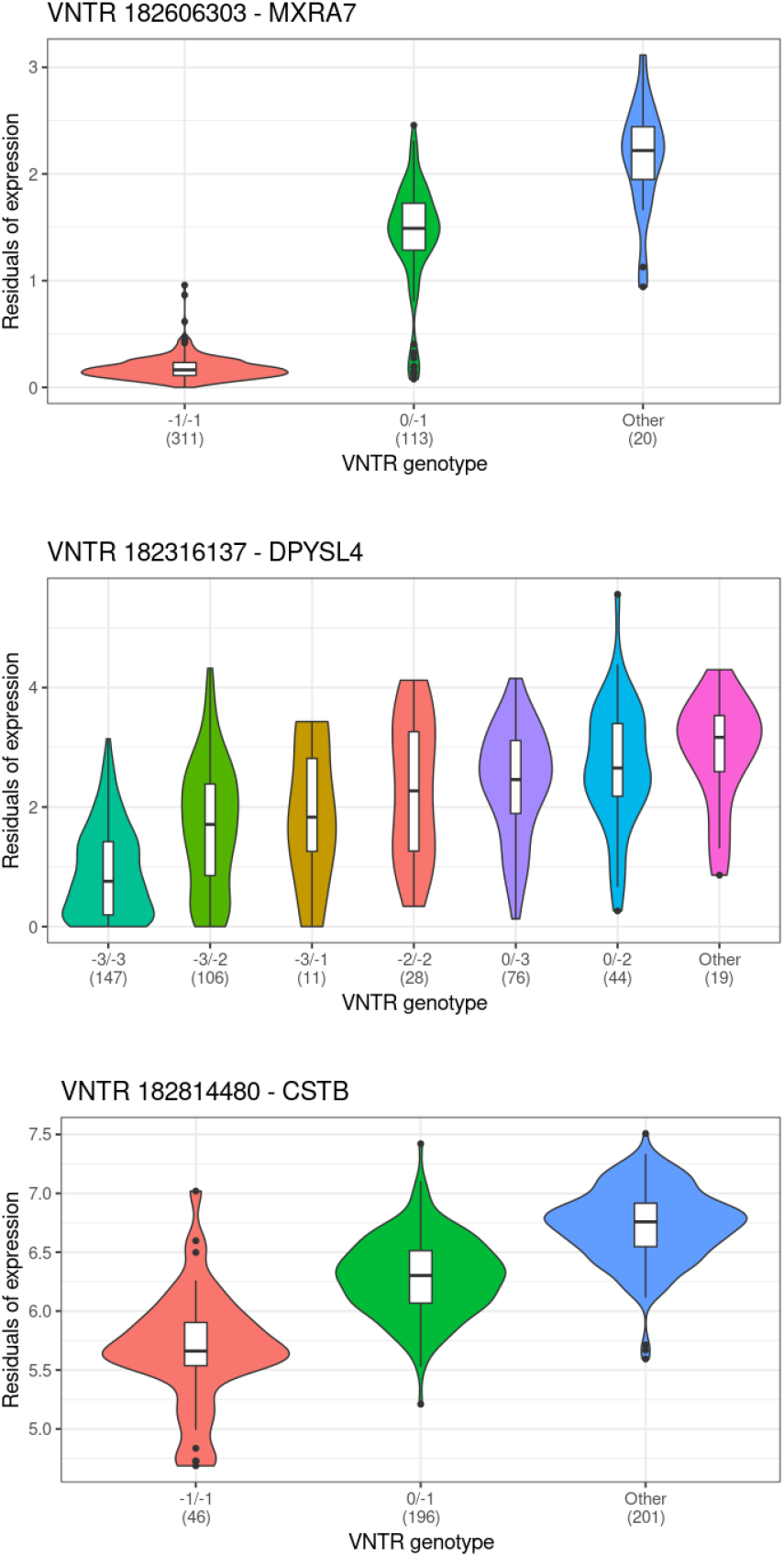
Gene expression differences and VNTR genotype. Shown are violin plots of gene expression values (log2 normalized TPM) for three genes which displayed significant differential expression when samples were partitioned by VNTR allele genotype. Additional examples are shown in Supplementary Figures S26–S30. Genotype is indicated in labels on the X-axis and numbers refer to copies gained or lost relative to the reference allele. “Other” indicates a partition with undetected alleles presumed outside the range of VNTRseek detection (see text). Number of samples in each partition is shown in parenthesis. In these examples, the effect size for at least one genotype class was significant. *Top*: VNTR 182606303 is upstream of MXRA7 and partially overlaps the 5’ UTR exon. *Middle*: VNTR 182316137 occurs inside the first intron of DPYSL4. *Bottom*: VNTR 182814480 occurs upstream of CSTB.

### One in five common VNTR loci have population-specific alleles

We further investigated whether VNTR *alleles* are population-specific and whether they can be used to predict ancestry. Understanding the occurrence of population-specific VNTR alleles will be useful when controlling for population effects in GWAS, and more generally in interpreting gene expression differences among people of different ancestry.

A total of 4,605 *alleles* from the common VNTR loci were classified as common if they were detected in at least 5% of the population (NYGC). We then constructed a matrix of presence/absence of each allele by sample and clustered the samples using Principal Component Analysis. We found that the first, second, fourth, and fifth principal components (PCs) separated the super populations as shown in Figure 3. Each PC captured a small fraction of the variation in the dataset, suggesting that there was substantial variation between individuals from the same population.

**Figure 3.**
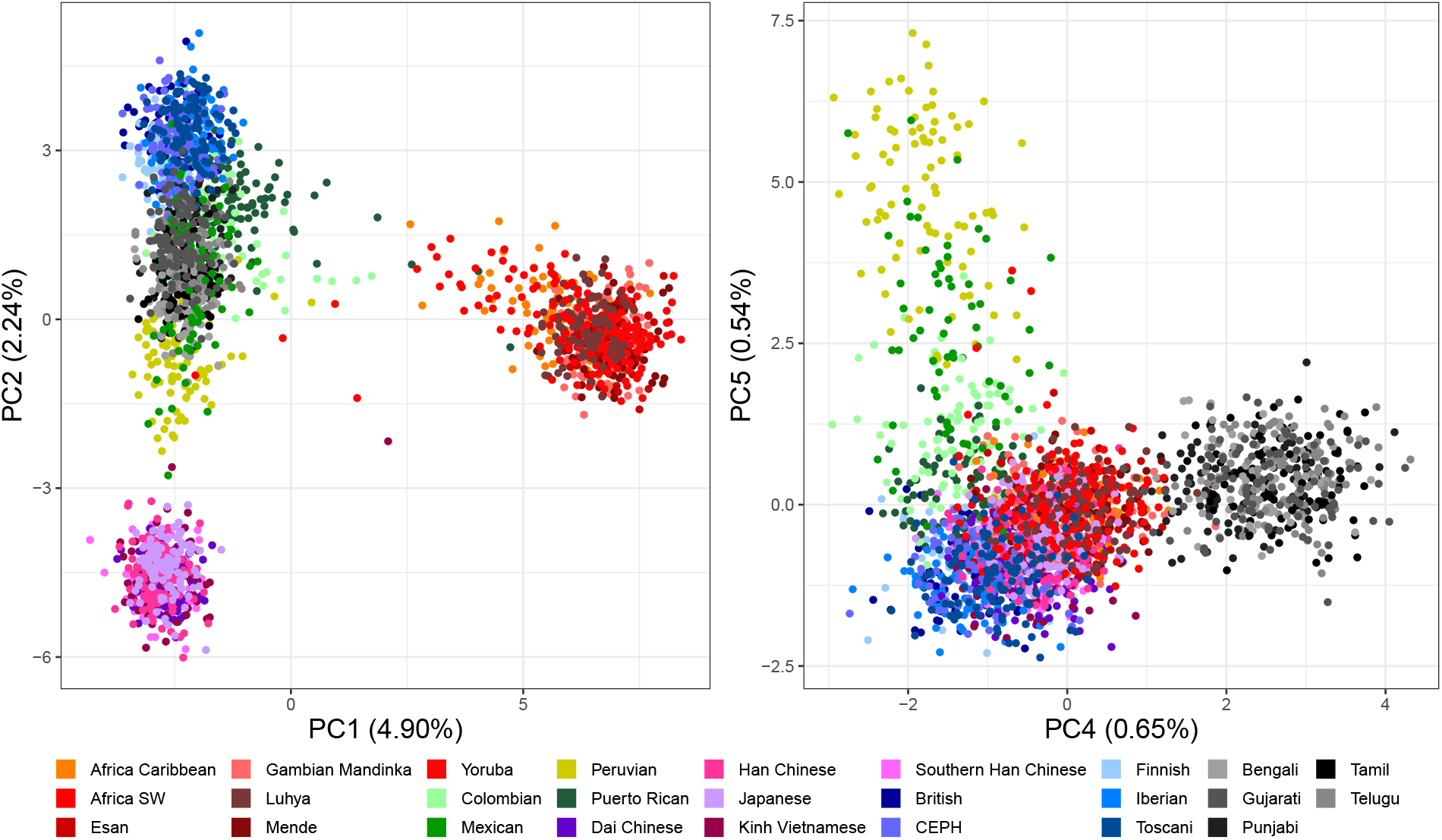
Principal Component Analysis (PCA) of common VNTR alleles in the NYGC population (150 bp). PCA was performed to reduce the dimensions of the data. *Left*: PC1 captured *∼* 5% of the variation and separated Africans from the other super-populations, suggesting that they had the greatest distance from the others. PC2 separated East Asian and European populations but left individuals from the Americas and South Asia mixed. *Right*: PC4 separated the South Asian population and PC5 separated the American populations. PC3 (not shown) captured batch effects due to differences in coverage. Some American sub-populations proved hardest to separate, likely due to ancestry mixing.

The first PC separated Africans, suggesting furthest evolutionary distance. The second PC separated East Asians. The third PC captured coverage bias. The fourth and fifth PCs separated South Asians and Americans, respectively. The American population had a sub-population of Puerto Ricans that clustered with the Iberian Spanish population, suggesting mixed ancestry (149). To show the power of these alleles to predict ancestry, we next trained a decision tree model (Supplementary Figure S19) using the top 10 PCs (11% of the total variation) and achieved a recall of *>*98% on every population when applied to the 30% test partition (Supplementary Table S10).

A one-sided Fisher’s Exact Test was applied to determine the population-specific VNTR alleles that were over-represented in one population versus all the others. A total of 3,850 VNTR alleles were identified as population-specific in one or more super-populations, corresponding to 1,096 VNTR loci (Supplementary Figures S20 and S21). The complete list of population-specific alleles can be found in Supplementary Material `Superpopulation VNTRs.txt`. These loci overlapped with 689 genes and 51 coding exons. Africans had the highest number of population-specific alleles (266), followed by East Asians (65), while Americans had the lowest (13), suggesting more mixed ancestry. We observed 63 loci that had a population-specific allele in each population. Figure 4 illustrates seven of the top population-specific loci in a “virtual gel” representation, mimicking the appearance of bands on an agarose gel for easier interpretation. Forty-eight genes that displayed expression differences correlated with VNTR genotype were associated with population-specific VNTR loci (Supplementary Table S15), including the VNTR 182316137 associated with the gene DPYSL4, discussed in the previous section, which exhibited seven different alleles, ? five of which were population-specific.

**Figure 4.**
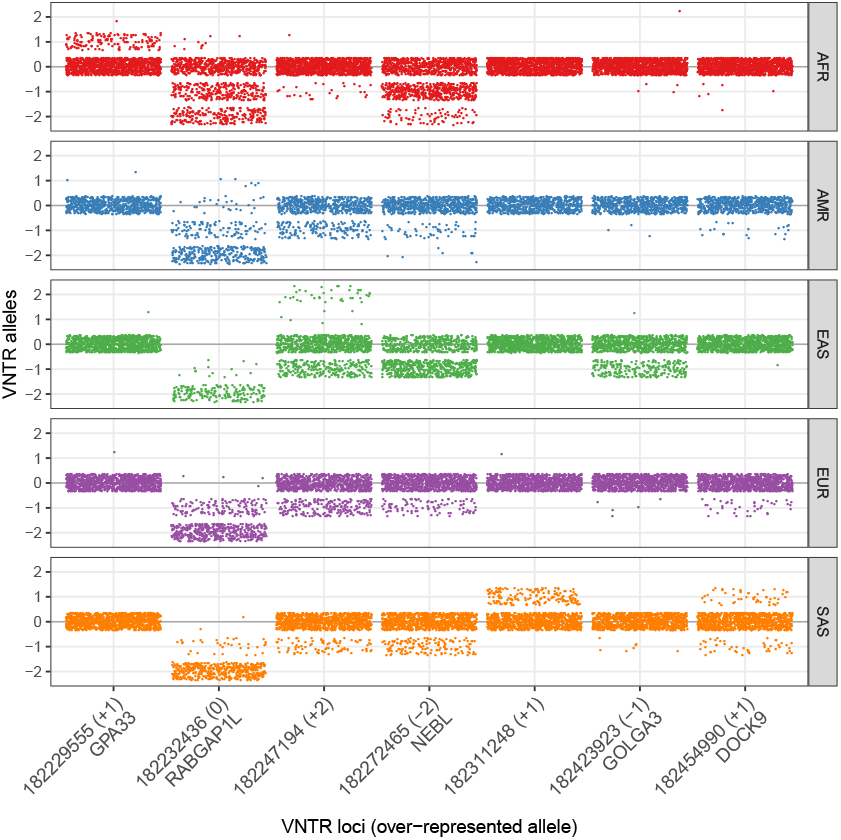
“Virtual gel” representation of seven population-specific VNTR alleles. Each dot represents an allele in one sample. Samples are separated vertically by super-population. Dots are jiggered in a rectangular area to reduce overlap. Population-specific alleles show up as bands over-represented in one population. Numbers and labels at bottom are VNTR locus ids with nearby genes indicated and the population-specific allele expressed as copy number change (+1, -2, *etc*.) from the reference. For example, in the leftmost column, the +1 allele was over-represented in the African population. Note that the allele bias towards pattern copy loss relative to the reference allele is apparent and that at one locus (second from left) the reference allele was the population-specific allele since almost no reference alleles were observed in the four other populations. The details of these seven loci are given in Supplementary Table S11.

Finally, to identify potential functional roles of the population-specific VNTR loci we performed Gene Set Enrichment Analysis (GSEA) for the associated genes against the Broad Institute MSigDB (150). Genes overlapping with the population-specific VNTRs were enriched for Endocytosis (hsa04144), Fatty acid metabolism (hsa01212), and Arrhythmogenic right ventricular cardiomyopathy (ARVC) (hsa05412) pathways (Supplementary Table S12). Among the GO biological processes affected by these genes were neurogenesis (GO:0022008; FDR=3.62*e*^−8^), neuron differentiation (GO:0030182; FDR=2.52*e*^−7^), and neuron development (GO:0048666; FDR=4.31*e*^−7^). These processes are potentially related to other findings that have linked VNTRs to neurodegenerative disorders and cognitive abilities (28, 36, 56, 57, 58, 59, 60, 61, 62, 151) (Supplementary material `Population specific Go BP.xlsx`). The GO term *behavior* (GO:0007610; FDR=2.22*e*^−4^) was also found, which could be related to the association of VNTR loci with aggressive behavior (152, 153, 154). Other notable GO terms were regulation of muscle contraction (GO:0006937) and neuromuscular processes related to balancing (GO:0050885) with FDR *<*1%. The genes were also highly enriched in midbrain neurotype cell gene signatures (FDR=5.49*e*^−25^), which might affect movement and emotions (155, 156, 157).

### Accuracy of VNTR predictions

To show the reliability of our results, we experimentally validated VNTR predictions at 13 loci in the three related AJ genomes, and also compared VNTRseek predictions to alleles experimentally validated in the literature. We additionally used accurate long reads on one genome (HG002) to find evidence of the predicted alleles. Separately, we showed the consistency of our predictions in two ways: first, we looked at inheritance consistency among four trios (mother, father, child), and second, we compared result for genomes sequenced on two different platforms.

#### Experimental validation

All but one of the 66 predicted VNTR alleles were confirmed at 13 loci in the three related AJ genomes (child HG002, father HG003, and mother HG004). In the remaining case, two predicted alleles were separated by only 15 nucleotides and could not be distinguished. At two loci, other bands were also observed. In one, all three family members contained an allele outside the detectable range of VNTRseek (longer than the reads). In the other, one allele that was detectable was missed in two family members. (See Table 3 for a summary of results; Supplementary Table S6 for details of the experiment and Supplementary Figures S13–S17 for gel images.)

**Table 3.** Experimental validation results: Thirteen VNTR loci were selected for experimental validation in the AJ trio. All but one of the 66 bands predicted by VNTRseek were validated. *†*For the remaining band, the results were questionable because the two predicted alleles for the father were only 15 nucleotides different in length, which was too close to distinguish in the image. &For all three individuals, the gel contained bands (**bold**) not predicted (or detectable) by VNTRseek. The extra band for the son corresponded to the -1 allele as found in the PacBio reads. The father’s extra band appeared to match with the -1 allele. The mother’s extra band appeared to be a +1 allele. *#*An extra band for the mother and son (**bold**) was not predicted by VNTRseek, although it seemed to match the +1 allele that was detectable. +These VNTRs overlapped regulatory sites that target the given genes.

| # | TR id | Pattern size | Ref. copies | Gene | VNTRseek predicted |  |  | Experimental validation |  |  |
| --- | --- | --- | --- | --- | --- | --- | --- | --- | --- | --- |
|  |  |  |  |  | C | F | M | C | F | M |
| 1* | 182316181 | 105 | 4 | STK32C | -2 | -2 | -2 | -2,-1 | -2,-1 | -2,+1 |
| 2 | 182316985 | 27 | 6 | LINC01168<sup>+</sup> | -3,0 | -1,0 | -3,-2 | yes | yes | yes |
| 3 | 182453735 | 30 | 2 | DNAJC3 | 0,+1 | 0,+1 | 0,+1 | yes | yes | yes |
| 4 | 182461997 | 38 | 7 | RASA3 | -5,-4 | -5,-4 | -5,-2 | yes | yes | yes |
| 5 | 182493720 | 70 | 3 | BEGAIN | 0 | 0 | 0,-1 | yes | yes | yes |
| 6 | 182515357 | 34 | 8 | MEGF11 | -5 | -5,-6 | -5 | yes | yes | yes |
| 7 | 182608886 | 27 | 6 | RPTOR | 0,-2 | 0,-3 | -2,+1 | yes | yes | yes |
| 8 | 182620950 | 48 | 3 | RNF138 | 0,-1 | 0,-1 | 0,-1 | yes | yes | yes |
| 9 | 182982510 | 34 | 4 | SLC12A7 | 0,-1 | 0 | 0,-1 | yes | yes | yes |
| 10<sup>‡</sup> | 183046759 | 38 | 4 | ARL10 | 0 | 0,-1 | -1 | 0,+1 | yes | -1,+1 |
| 11<sup>†</sup> | 183081195 | 15 | 4 | TENT5A | +2,-1 | +2,+1 | +2,-1 | yes | +2 | yes |
| 12 | 183117043 | 17 | 9 | MRM2<sup>+</sup>, LFNG<sup>+</sup> | -5,-3 | -5,-3 | 0,-3 | yes | yes | yes |
| 13 | 183169331 | 15 | 4 | IRF5 | -2 | -2,0 | -2 | yes | yes | yes |

**Table 4.** Dataset sources.

| Dataset (individuals) | URL or Accession numbers |
| --- | --- |
| 1000 Genomes phase 3 HC (30) | EBI: 1000 Genomes Project |
| NYGC (2,504) | Data collection 30X on GRCh38<br>ENA project: PRJEB31736 |
| SGDP (253) | IGSR: Data collection SGDP |
| GIAB (7) | NCBI SRA project: SRP047086 |
| CHM1 (1) | NCBI SRA: SRX652547 |
| CHM13 (1) | NCBI SRA: SRX1009644 |
| Tumor/Normal Pairs (4) | Illumina basespace project |
| PacBio (1) | NCBI SRA: SRX5327410 |

We also compared VNTRseek predictions in three datasets from the NA12878 genome with VNTRs validated in the adVNTR paper (95). Out of the original 17 VNTR loci experimentally validated in that paper, four were not included in our reference set and for one, the matching TR could not be determined. In total, 11 out of 16 detectable alleles were correctly predicted, four were not found in the NA12878 sample with sufficient read size (250 bp), and one was incorrectly predicted in the HG001 sample and not found in the other two (Supplementary Table S7).

#### Validation of predicted VNTRs using long reads

PacBio Circular Consensus Sequencing reads from the HG002 genome (103), with an average length of 13.5 Kbp and an estimated 99.8% sequence accuracy, were computationally tested to determine if they confirmed VNTRseek predicted alleles for the GIAB Illumina reads from the same genome. Overall, more than 97% of predicted alleles were confirmed, and at the predicted VNTR loci, more than 87% of alleles were confirmed (Supplementary Table S8).

#### VNTR predictions are consistent with Mendelian inheritance

We compared the predicted alleles in four trios (CEU and YRI trios from 1000 Genomes; Chinese HAN and AJ from GIAB), testing loci on autosomes and X and Y chromosomes (see Methods). In all cases, only a handful of loci were inconsistent (Supplementary Table S9).

#### VNTR non-reference allele predictions showed substantial consistency across platforms

In 2015, the 1000 Genomes Phase 3 sequenced 30 genomes using Illumina HiSeq2500 at read length 250 bp. In 2020, 27 of those 30 genomes were resequenced by NYGC using Illumina Novaseq 6000 at read length 150 bp. Comparing VNTR loci genotyped in both platforms and only non-reference alleles detectable at both read lengths, agreement ranged from 76%–91% (Supplementary Figure S18). When including reference alleles, the agreement increased to greater than 99%. Note, however, that read coverage was not the same for both datasets, causing variation in statistical power.

## DISCUSSION

The current study represents, to our knowledge, the largest analysis of human whole genome sequencing data to detect copy number variable tandem repeats (VNTRs) and greatly expands the growing information on this class of genetic variation. The TRs genotyped consisted of some 184,000 minisatellites occupying the mid-range of pattern sizes, from seven to 126 bp. Our results reveal that nearly 20% (35,828) are variable, exhibiting at least one non-reference allele, a number much larger than has been generally understood. Moreover, we have classified a large subset of these (5,676) as *common VNTRs*, with non-reference alleles occurring in *>*5% of the population. When considering the largest dataset in our study (2,504 individuals), we found that, on average, each genome exhibited non-reference alleles at 1,951 VNTR loci and among those, 1,783 were common VNTRs.

In addition to their widespread occurrence, further evidence of minisatellite VNTR importance can be seen in the enrichment of these loci in genes and gene regulatory regions (promoters, transcription factor binding sites, DNAse hypersensitive sites, and CpG islands). Our entire set of VNTRs overlapped with 7,698 protein coding genes and 3,512 exons. The common VNTRs occurred within or were proximal to over 2,173 protein coding genes, including overlapping with 254 exons. Biological function enrichment among these genes included neuron development and differentiation, and behavior. Examples include low-density lipoprotein receptor-related protein 6 (LRP6), a co-receptor for Wnt signaling, which is down-regulated in Alzheimer’s disease and which appears critical for maintaining synaptic integrity (158); Down syndrome cell adhesion molecule-like 1 (Dscaml1), which appears essential for development of GABAergic neurons in the entorhinal cortex (159); and ZMIZ1, a transcription factor co-activator, for which mutant allele over-expression leads to pyramidal neuron morphology abnormalities (160). These observations are consistent with the finding that VNTR expansions in humans compared to primates are associated with gain of cognitive abilities (24), and possible involvement of VNTRS with many neurodegenerative diseases and behavioral disorders (87).

The overabundance of VNTR proximity to genes suggests that variability at these loci could affect gene expression and indeed, we observed that the expression levels of 187 genes were significantly correlated with the presence of specific VNTR alleles in lymphoblastoid cell lines of 445 individuals. Others have also recently detected minisatellite VNTR eQTLs. In (25), expression levels in 46 tissues for 6,802 genes were tested and 161 eQTLs were found. Thirteen of those genes coincided with ones detected in our study: BPTF, CCDC57, CCDC66, EPDR1, EPS8L2, LSS, MVB12A, PTPRVP, S100A10, SNAPC1, TMEM52, TRAPPC2L, ZNF736. Similarly, in (26) 21 VNTRs were found associated with expression in 38 genes. However, their VNTR alleles were larger and did not overlap the ones we tested in this study.

eQTLs have also been found among the variant STR class. In (161), 2,060 genes were found to be affected by eQTL STRs. In another study (162), 28,375 variant STR loci were associated with differential expression of 12,494 genes in at least one tissue (FDR*<*10%) and 1,420 of the loci were predicted, by CAVIAR (163), to be causal in 17 tissues.

These findings are suggestive, but more study is required, both to determine if there is more evidence of *tissue specific* gene expression variation associated with VNTR genotypes (25, 26, 87) and if such correlational differences can be definitively tied to actions associated with specific VNTR alleles such as regulator binding affinity changes in regulatory regions. For more elaborate studies such as these, it will be essential that for each sample used to measure gene expression, the raw whole genome sequencing data be available, so that specialized software programs, such as VNTRseek can be used to determine VNTR genotype.

The frequency of VNTR occurrence and possible effects on gene expression suggest that minisatellite VNTR loci could be useful in genome-wide association studies (GWAS). However, it is well known that hidden differences can lead to misinterpretation of GWAS results, and care is particularly important when those differences are tied to human ancestry. Relevant to this, we have determined that 1,096 of the common VNTR loci contain alleles showing significant population specificity and that these loci intersect with 689 genes. Understanding such hidden variability will be essential for interpreting GWAS and future studies should investigate possible haplotype linkages between specific VNTR alleles and nearby SNP alleles.

Population-specific alleles also have the potential for use in tracing early human migration. We have shown through principal component analysis with common VNTR alleles that super-populations are easily separated. Further, we have constructed a decision tree based on common VNTR alleles that obtains nearly perfect classification of individuals at the super population level. It will be interesting to see whether, with more information, classification can be refined further to encompass specific sub-populations, whether a minimal minisatellite VNTR set can be established for high accuracy population classification, and whether VNTR alleles can be used to estimate mixed ancestry as is done now with SNP haplotyping.

Despite the high precision of VNTRseek, (see Supplementary Figure S32 and Supplementary Table S16– our curation of VNTR loci has certainly produced an undercount. This is true because VNTRseek requires that the tandem array fit within a read. Longer reads will help, but the abundance of high-coverage. low-error, long-read datasets is currently limited. Alternate methods exist (87, 95), but these have not reported an ability to handle *macrosatellite* VNTRs where the arrays and patterns are hundreds to thousands of base pairs long. For this range of the tandem repeat spectrum, new tools must be developed.

Another limitation comes from use of the Tandem Repeats Finder, which requires that the array contain at least 1.9 copies to be detected. At read length 150 bp, which included the majority of our samples, a gain of one copy compared to the reference genome could be detected in 82% of the TR loci while loss of one copy could be detected in only 16%. Despite this imbalance, one copy loss was observed nearly 40% more often than one copy gain, an important observation with regard to potential tandem repeat copy number bias in the reference genome.

Previous studies on VNTR prevalence in the human genome have been limited to a subset of minisatellites inside the transcriptome and a limited number of genomes. Here, we have shown a broader prevalence of VNTR loci and suggested their importance with regard to gene function, population studies, and GWAS. Future research can be expected to further enhance our understanding of this important class of genomic variation.

## Supporting information

Reference_TRs

Summary_of_results.txt

Common_VNTRs.txt

Private_VNTRs

Major_genotypes.txt

Superpopulation_VNTRs.txt

Population_specific_GO_BP.xlsx

RNA_VNTRs.txt

Experiment_primers.txt

## DATA AVAILABILITY

The reference TR set files, output VCF files, and the pre-processed data files along with the code to create figures and tables are published at: DOI 10.5281/zenodo.4065850 VNTRseek can be downloaded at: https://github.com/Benson-Genomics-Lab/VNTRseek.

## ACKNOWLEDGEMENT

This work was supported in part by NSF grants IIS-1017621, IIS-1423022 and DBI-1559829 and NIH grant R35 GM128625.

We thank Thomas Gilmore (Boston University) for helpful discussions and comments on the manuscript, Alberto Cruz-Martiń (Boston University) for discussion on gene association with neuron development, and Xiaoling Zhang (Boston University School of Medicine) for her help with the eQTL analysis.

## References

1. Treangen, T. J. and Salzberg, S. L. (2012) Repetitive DNA and next-generation sequencing: computational challenges and solutions. Nature Reviews Genetics, 13(1), 36.

2. de Koning, A. J., Gu, W., Castoe, T. A., Batzer, M. A., and Pollock, D. D. (2011) Repetitive elements may comprise over two-thirds of the human genome. PLoS Genetics, 7(12), e1002384.

3. Lim, K. G., Kwoh, C. K., Hsu, L. Y., and Wirawan, A. (2013) Review of tandem repeat search tools: a systematic approach to evaluating algorithmic performance. Briefings in Bioinformatics, 14(1), 67–81.

4. Richard, G.-F., Kerrest, A., and Dujon, B. (2008) Comparative genomics and molecular dynamics of DNA repeats in eukaryotes. Microbiology and Molecular Biology Reviews, 72(4), 686–727.

5. Taylor, J. S. and Breden, F. (2000) Slipped-strand mispairing at noncontiguous repeats in Poecilia reticulata: a model for minisatellite birth. Genetics, 155(3), 1313–1320.

6. Levinson, G. and Gutman, G. A. (1987) Slipped-strand mispairing: a major mechanism for DNA sequence evolution.. Molecular Biology and Evolution, 4(3), 203–221.

7. Madsen, C. S., Ghivizzani, S. C., and Hauswirth, W. W. (1993) In vivo and in vitro evidence for slipped mispairing in mammalian mitochondria. Proceedings of the National Academy of Sciences, 90(16), 7671–7675.

8. Jeffreys, A. J., Neil, D. L., and Neumann, R. (1998) Repeat instability at human minisatellites arising from meiotic recombination. The EMBO Journal, 17(14), 4147–4157.

9. Debrauwére, H., Buard, J., Tessier, J., Aubert, D., Vergnaud, G., and Nicolas, A. (1999) Meiotic instability of human minisatellite CEB1 in yeast requires DNA double-strand breaks. Nature Genetics, 23(3), 367.

10. Pâques, F., Richard, G.-F., and Haber, J. E. (2001) Expansions and contractions in 36-bp minisatellites by gene conversion in yeast. Genetics, 158(1), 155–166.

11. Bustamante, A. V., Sanso, A. M., Segura, D., Parma, A. E., and Lucchesi, P. M. A. (2013) Dynamic of mutational events in variable number tandem repeats of Escherichia coli O157: H7. BioMed Research International, 2013.

12. Vogler, A. J., Keys, C., Nemoto, Y., Colman, R. E., Jay, Z., and Keim, P. (2006) Effect of repeat copy number on variable-number tandem repeat mutations in Escherichia coli O157: H7. Journal of Bacteriology, 188(12), 4253–4263.

13. Fu, S., Octavia, S., Wang, Q., Tanaka, M. M., Tay, C. Y., Sintchenko, V., and Lan, R. (2016) Evolution of variable number tandem repeats and its relationship with genomic diversity in Salmonella Typhimurium. Frontiers in Microbiology, 7, 2002.

14. Verstrepen, K. J., Jansen, A., Lewitter, F., and Fink, G. R. (2005) Intragenic tandem repeats generate functional variability. Nature Genetics, 37(9), 986.

15. Legendre, M., Pochet, N., Pak, T., and Verstrepen, K. J. (2007) Sequence-based estimation of minisatellite and microsatellite repeat variability. Genome Research, 17(12), 1787–1796.

16. Panigrahi, I. (2018) Genetic Fingerprinting for Human Diseases: Applications and Implications. In DNA Fingerprinting: Advancements and Future Endeavors pp. 141–150 Springer.

17. Sinha, M., Rao, I. A., and Mitra, M. (2018) Molecular Basis of Identification Through DNA Fingerprinting in Humans. In DNA Fingerprinting: Advancements and Future Endeavors pp. 129–140 Springer.

18. Imam, J., Reyaz, R., Rana, A. K., and Yadav, V. K. (2018) DNA Fingerprinting: Discovery, Advancements, and Milestones. In DNA Fingerprinting: Advancements and Future Endeavors pp. 3–24 Springer.

19. Denoeud, F., Vergnaud, G., and Benson, G. (2003) Predicting human minisatellite polymorphism. Genome Research, 13(5), 856–867.

20. Deka, R., Chakraborty, R., and Ferrell, R. E. (1991) A population genetic study of six VNTR loci in three ethnically defined populations. Genomics, 11(1), 83–92.

21. Deka, R., DeCroo, S., Yu, L. M., and Ferrell, R. E. (1992) Variable number of tandem repeat (VNTR) polymorphism at locus D17S5 (YNZ22) in four ethnically defined human populations. Human Genetics, 90(1-2), 86–90.

22. Hancock, J. M. and Santibáñez-Koref, M. F. (1998) Trinucleotide Expansion Diseases in the Context of Micro-and Minisatellite Evolution Hammersmith Hospital, April 1–3, 1998. The EMBO Journal, 17(19), 5521–5524.

23. Duitama, J., Zablotskaya, A., Gemayel, R., Jansen, A., Belet, S., Vermeesch, J. R., Verstrepen, K. J., and Froyen, G. (2014) Large-scale analysis of tandem repeat variability in the human genome. Nucleic Acids Research, 42(9), 5728–5741.

24. Sonay, T. B., Carvalho, T., Robinson, M. D., Greminger, M. P., Krützen, M., Comas, D., Highnam, G., Mittelman, D., Sharp, A., Marques-Bonet, T., et al. (2015) Tandem repeat variation in human and great ape populations and its impact on gene expression divergence. Genome Research, 25(11), 1591–1599.

25. Bakhtiari, M., Park, J., Ding, Y.-C., Shleizer-Burko, S., Neuhausen, S. L., Halldórsson, B. V., Stefansson, K., Gymrek, M., and Bafna, V. (2020) Variable Number Tandem Repeats mediate the expression of proximal genes. bioRxiv, doi:10.1101/2020.05.25.114082,.

26. Lu, T.-Y. T., Chaisson, M. J., Consortium, H. G. S. V., et al. (2020) Profiling variable-number tandem repeat variation across populations using repeat-pangenome graphs. bioRxiv, doi: 10.1101/2020.08.13.249839,.

27. Trepicchio, W. L. and Krontiris, T. G. (1992) Members of the rel/NF-χB family of transcriptional regulatory proteins bind the HRAS1minisatellite DNA sequence. Nucleic Acids Research, 20(10), 2427– 2434.

28. Krontiris, T. G., Devlin, B., Karp, D. D., Robert, N. J., and Risch, N. (1993) An association between the risk of cancer and mutations in the HRAS1 minisatellite locus. New England Journal of Medicine, 329(8), 517–523.

29. Wang, S., Wang, M., Yin, S., Fu, G., Li, C., Chen, R., Li, A., Zhou, J., Zhang, Z., and Liu, Q. (2008) A novel variable number of tandem repeats (VNTR) polymorphism containing Sp1 binding elements in the promoter of XRCC5 is a risk factor for human bladder cancer. Mutation Research/Fundamental and Molecular Mechanisms of Mutagenesis, 638(1-2), 26–36.

30. Zukic, B., Radmilovic, M., Stojiljkovic, M., Tosic, N., Pourfarzad, F., Dokmanovic, L., Janic, D., Colovic, N., Philipsen, S., Patrinos, G. P., et al. (2010) Functional analysis of the role of the TPMT gene promoter VNTR polymorphism in TPMT gene transcription. Pharmacogenomics, 11(4), 547–557.

31. Vasiliou, S. A., Ali, F. R., Haddley, K., Cardoso, M. C., Bubb, V. J., and Quinn, J. P. (2012) The SLC6A4 VNTR genotype determines transcription factor binding and epigenetic variation of this gene in response to cocaine in vitro. Addiction Biology, 17(1), 156–170.

32. Vafiadis, P., Bennett, S. T., Todd, J. A., Nadeau, J., Grabs, R., Goodyer, C. G., Wickramasinghe, S., Colle, E., and Polychronakos, C. (1997) Insulin expression in human thymus is modulated by INS VNTR alleles at the IDDM2 locus. Nature Genetics, 15(3), 289–292.

33. Greenwood, T. A. and Kelsoe, J. R. (2003) Promoter and intronic variants affect the transcriptional regulation of the human dopamine transporter gene. Genomics, 82(5), 511–520.

34. Lovejoy, E., Scott, A., Fiskerstrand, C., Bubb, V., and Quinn, J. (2003) The serotonin transporter intronic VNTR enhancer correlated with a predisposition to affective disorders has distinct regulatory elements within the domain based on the primary DNA sequence of the repeat unit. European Journal of Neuroscience, 17(2), 417–420.

35. Klenova, E., Scott, A. C., Roberts, J., Shamsuddin, S., Lovejoy, E. A., Bergmann, S., Bubb, V. J., Royer, H.-D., and Quinn, J. P. (2004) YB-1 and CTCF differentially regulate the 5-HTT polymorphic intron 2 enhancer which predisposes to a variety of neurological disorders. Journal of Neuroscience, 24(26), 5966–5973.

36. De Roeck, A., Duchateau, L., Van Dongen, J., Cacace, R., Bjerke, M., Van den Bossche, T., Cras, P., Vandenberghe, R., De Deyn, P. P., Engelborghs, S., et al. (2018) An intronic VNTR affects splicing of ABCA7 and increases risk of Alzheimer’s disease. Acta Neuropathologica, 135(6), 827–837.

37. Pacheco, A., Berger, R., Freedman, R., and Law, A. J. (2019) A VNTR Regulates miR-137 expression through novel Alternative Splicing and contributes to Risk for Schizophrenia. Scientific Reports, 9(1), 1–12.

38. Schoots, O. and Van Tol, H. (2003) The human dopamine D4 receptor repeat sequences modulate expression. The Pharmacogenomics Journal, 3(6), 343–348.

39. Xiao, X., Jones, G., Sevilla, W. A., Stolz, D. B., Magee, K. E., Haughney, M., Mukherjee, A., Wang, Y., and Lowe, M. E. (2016) A carboxyl ester lipase (CEL) mutant causes chronic pancreatitis by forming intracellular aggregates that activate apoptosis. Journal of Biological Chemistry, 291(44), 23224–23236.

40. Ræder, H., Johansson, S., Holm, P. I., Haldorsen, I. S., Mas, E., Sbarra, V., Nermoen, I., Eide S. Å., Grevle, L., Bjørkhaug, L., et al. (2006) Mutations in the CEL VNTR cause a syndrome of diabetes and pancreatic exocrine dysfunction. Nature Genetics, 38(1), 54.

41. Willems, T., Gymrek, M., Highnam, G., Mittelman, D., Erlich, Y., Consortium, . G. P., et al. (2014) The landscape of human STR variation. Genome Research, 24(11), 1894–1904.

42. Willems, T., Zielinski, D., Yuan, J., Gordon, A., Gymrek, M., and Erlich, Y. (2017) Genome-wide profiling of heritable and de novo STR variations. Nature Methods, 14(6), 590–592.

43. Mallick, S., Li, H., Lipson, M., Mathieson, I., Gymrek, M., Racimo, F., Zhao, M., Chennagiri, N., Nordenfelt, S., Tandon, A., et al. (2016) The Simons genome diversity project: 300 genomes from 142 diverse populations. Nature, 538(7624), 201–206.

44. Gettings, K. B., Borsuk, L. A., Zook, J., and Vallone, P. M. (2019) Unleashing novel STRs via characterization of genome in a bottle reference samples. Forensic Science International: Genetics Supplement Series, 7(1), 218–220.

45. Krishnan, V., Utiramerur, S., Ng, Z., Datta, S., Snyder, M. P., and Ashley, E. A. (2021) Benchmarking workflows to assess performance and suitability of germline variant calling pipelines in clinical diagnostic assays. BMC Bioinformatics, 22(1), 1–17.

46. Brouwer, J. R., Willemsen, R., and Oostra, B. A. (2009) Microsatellite repeat instability and neurological disease. BioEssays, 31(1), 71–83.

47. Rohilla, K. J. and Gagnon, K. T. (2017) RNA biology of disease-associated microsatellite repeat expansions. Acta Neuropathologica Communications, 5(1), 1–22.

48. Hannan, A. J. (2018) Tandem repeats mediating genetic plasticity in health and disease. Nature Reviews Genetics, 19(5), 286.

49. Rodriguez, C. and Todd, P. (2019) New pathologic mechanisms in nucleotide repeat expansion disorders. Neurobiology of Disease, 130, 104515.

50. Beck, R., Monument, M. J., Watkins, W. S., Smith, R., Boucher, K. M., Schiffman, J. D., Jorde, L. B., Randall, R. L., and Lessnick, S. L. (2012) EWS/FLI-responsive GGAA microsatellites exhibit polymorphic differences between European and African populations. Cancer Genetics, 205(6), 304–312.

51. Boulay, G., Volorio, A., Iyer, S., Broye, L. C., Stamenkovic, I., Riggi, N., and Rivera, M. N. (2018) Epigenome editing of microsatellite repeats defines tumor-specific enhancer functions and dependencies. Genes & Development, 32(15-16), 1008–1019.

52. Nacev, B. A., Jones, K. B., Intlekofer, A. M., Jamie, S., Allis, C. D., Tap, W. D., Ladanyi, M., and Nielsen, T. O. (2020) The epigenomics of sarcoma. Nature Reviews Cancer, 20(10), 608–623.

53. Antwi-Boasiako, C., Dzudzor, B., Kudzi, W., Doku, A., Dale, C., Sey, F., Otu, K., Boatemaa, G., Ekem, I., Ahenkorah, J., et al. (2018) Association between eNOS Gene Polymorphism (T786C and VNTR) and Sickle Cell Disease Patients in Ghana. Diseases, 6(4), 90.

54. Ksiazek, K., Blaszczak, J., and Buraczynska, M. (2019) IL4 gene VNTR polymorphism in chronic periodontitis in end-stage renal disease patients. Oral Diseases, 25(1), 258–264.

55. Cong, L., Tu, G., and Liang, D. (2018) A systematic review of the relationship between the distributions of aggrecan gene VNTR polymorphism and degenerative disc disease/osteoarthritis. Bone & Joint Research, 7(4), 308–317.

56. Katsumata, Y., Fardo, D. W., Bachstetter, A. D., Artiushin, S. C., Wang, W.-X., Wei, A., Brzezinski, L. J., Nelson, B. G., Huang, Q., Abner, E. L., et al. (2019) Alzheimer Disease Pathology-Associated Polymorphism in a Complex Variable Number of Tandem Repeat Region Within the MUC6 Gene, Near the AP2A2 Gene. Journal of Neuropathology & Experimental Neurology,.

57. Chang, H.-I., Chang, Y.-T., Tsai, S.-J., Huang, C.-W., Hsu, S.-W., Liu, M.-E., Chang, W.-N., Lien, C.-Y., Huang, S.-H., Lee, C.-C., et al. (2019) MAOA-VNTR Genotype Effects on Ventral Striatum-Hippocampus Network in Alzheimer’s Disease: Analysis Using Structural Covariance Network and Correlation with Neurobehavior Performance. Molecular Neurobiology, 56(6), 4518–4529.

58. Scott, H., Nelson, P., Hopwood, J., and Morris, C. (1991) PCR of a VNTR linked to mucopolysaccharidosis type I and Huntington disease. Nucleic Acids Research, 19(22), 6348.

59. Hoxha, B., Goçi, A. U., Agani, F., Haxhibeqiri, S., Haxhibeqiri, V., Sabic, E. D., Kucukalic, S., Bravo, A. M., Kucukalic, A., Dzubur, A. K., et al. (2019) The Role of TaqI DRD2 (rs1800497) and DRD4 VNTR Polymorphisms in Posttraumatic Stress Disorder (PTSD). Psychiatria Danubina, 31(2), 263–268.

60. Šer ý, O., Paclt, I., Drtílková, I., Theiner, P., Kopečková, M., Zvolský, P., and Balcar, V. J. (2015) A 40-bp VNTR polymorphism in the 3’-untranslated region of DAT1/SLC6A3 is associated with ADHD but not with alcoholism. Behavioral and Brain Functions, 11(1), 21.

61. Grünblatt, E., Werling, A. M., Roth, A., Romanos, M., and Walitza, S. (2019) Association study and a systematic meta-analysis of the VNTR polymorphism in the 3’-UTR of dopamine transporter gene and attention-deficit hyperactivity disorder. Journal of Neural Transmission, 126(4), 517–529.

62. Van Assche, E., Moons, T., Van Leeuwen, K., Colpin, H., Verschueren, K., Van Den Noortgate, W., Goossens, L., and Claes, S. (2016) Depressive symptoms in adolescence: The role of perceived parental support, psychological control, and proactive control in interaction with 5-HTTLPR. European Psychiatry, 35, 55–63.

63. Stolf, A. R., Cupertino, R. B., Müller, D., Sanvicente-Vieira, B., Roman, T., Vitola, E. S., Grevet, E. H., von Diemen, L., Kessler, F. H., Grassi-Oliveira, R., et al. (2019) Effects of DRD2 splicing-regulatory polymorphism and DRD4 48 bp VNTR on crack cocaine addiction. Journal of Neural Transmission, 126(2), 193–199.

64. Ramírez-Patiño, R., Figuera, L. E., Puebla-Pérez, A. M., Delgado-Saucedo, J. I., Legazpí-Macias, M. M., Mariaud-Schmidt, R. P., Ramos-Silva, A., Gutiérrez-Hurtado, I. A., Gómez Flores-Ramos, L., Zúñiga-González, G. M., et al. (2013) Intron 4 VNTR (4a/b) polymorphism of the endothelial nitric oxide synthase gene is associated with breast cancer in Mexican women. Journal of Korean Medical Science, 28(11), 1587–1594.

65. Vairaktaris, E., Serefoglou, Z. C., Yapijakis, C., Vassiliou, S., Nkenke, E., Avgoustidis, D., Vylliotis, A., Stathopoulos, P., Neukam, F. W., and Patsouris, E. (2007) The Platelet Glycoprotein Ibα VNTR Polymorphism is Associated with Risk for Oral Cancer. AntiCancer Research, 27(6B), 4121–4125.

66. Sousa, H., Santos, A. M., Catarino, R., Pinto, D., Moutinho, J., Canedo, P., Machado, J. C., and Medeiros, R. (2012) IL-1RN VNTR polymorphism and genetic susceptibility to cervical cancer in Portugal. Molecular Biology Reports, 39(12), 10837–10842.

67. Safarinejad, M. R., Safarinejad, S., Shafiei, N., and Safarinejad, S. (2013) Effects of the T-786C, G894T, and Intron 4 VNTR (4a/b) polymorphisms of the endothelial nitric oxide synthase gene on the risk of prostate cancer. In Urologic Oncology: Seminars and Original Investigations Elsevier Vol. 31, pp. 1132–1140.

68. Ibrahimi, M., Moossavi, M., Mojarad, E. N., Musavi, M., Mohammadoo-khorasani, M., and Shahsavari, Z. (2019) Positive correlation between interleukin-1 receptor antagonist gene 86bp VNTR polymorphism and colorectal cancer susceptibility: a case-control study. Immunologic Research, 67(1), 151–156.

69. Cui, J., Luo, J., Kim, Y. C., Snyder, C., Becirovic, D., Downs, B., Lynch, H., and Wang, S. M. (2016) Differences of variable number tandem repeats in XRCC5 promoter are associated with increased or decreased risk of breast cancer in BRCA gene mutation carriers. Frontiers in Oncology, 6, 92.

70. Al-Eitan, L. N., Rababa’h, D. M., Alghamdi, M. A., and Khasawneh, R. H. (2019) The influence of an IL-4 variable number tandem repeat (VNTR) polymorphism on breast cancer susceptibility. Pharmacogenomics and Personalized Medicine, 12, 201.

71. Ahn, E.-K., Kim, W.-J., Kwon, J.-A., Choi, P.-J., Kim, W. J., Sunwoo, Y., Heo, J., and Leem, S.-H. (2009) Variants of MUC5B minisatellites and the susceptibility of bladder cancer. DNA and Cell Biology, 28(4), 169–176.

72. Kwon, J.-A., Lee, S.-Y., Ahn, E.-K., Seol, S.-Y., Kim, M. C., Kim, S. J., Kim, S. I., Chu, I.-S., and Leem, S.-H. (2010) Short rare MUC6 minisatellites-5 alleles influence susceptibility to gastric carcinoma by regulating gene. Human Mutation, 31(8), 942–949.

73. Weitzel, J. N., Ding, S., Larson, G. P., Nelson, R. A., Goodman, A., Grendys, E. C., Ball, H. G., and Krontiris, T. G. (2000) The HRAS1 minisatellite locus and risk of ovarian cancer. Cancer Research, 60(2), 259–261.

74. Wang, L., Soria, J.-C., Chang, Y.-S., Lee, H.-Y., Wei, Q., and Mao, L. (2003) Association of a functional tandem repeats in the downstream of human telomerase gene and lung cancer. Oncogene, 22(46), 7123–7129.

75. Calvo, R., Pifarré, A., Rosell, R., Sánchez, J. J., Monzoó, M., Ribera, J. M., and Feliu, E. (1998) H-RAS 1 minisatellite rare alleles: a genetic susceptibility and prognostic factor for non-Hodgkin’s lymphoma. JNCI: Journal of the National Cancer Institute, 90(14), 1095–1098.

76. Yoon, S.-L., Jung, S.-I., Kim, W.-J., Kim, S. I., Park, I.-h., and Leem, S.-H. (2011) Variants of BORIS minisatellites and relation to prognosis of prostate cancer. Genes & Genomics, 33(1), 49–56.

77. Batra, S. K., McLendon, R. E., Koo, J. S., Castelino-Prabhu, S., Fuchs, H. E., Krischer, J. P., Friedman, H. S., Bigner, D. D., and Bigner, S. H. (1995) Prognostic implications of chromosome 17p deletions in human medulloblastomas. Journal of Neuro-Oncology, 24(1), 39–45.

78. Andersson, U., Osterman, P., Sjöström, S., Johansen, C., Henriksson, R., Brännström, T., Broholm, H., Christensen, H. C., Ahlbom, A., Auvinen, A., et al. (2009) MNS16A minisatellite genotypes in relation to risk of glioma and meningioma and to glioblastoma outcome. International Journal of Cancer, 125(4), 968–972.

79. Lim, S. W., Kim, H. R., Kim, H. Y., Huh, J. W., Kim, Y. J., Shin, J. H., Suh, S. P., Ryang, D. W., Kim, H. R., and Shin, M. G. (2012) High-frequency minisatellite instability of the mitochondrial genome in colorectal cancer tissue associated with clinicopathological values. International Journal of Cancer, 131(6), 1332–1341.

80. Xia, X., Rui, R., Quan, S., Zhong, R., Zou, L., Lou, J., Lu, X., Ke, J., Zhang, T., Zhang, Y., et al. (2013) MNS16A tandem repeats minisatellite of human telomerase gene and cancer risk: a meta-analysis. PLoS One, 8(8), e73367.

81. Leem, S. H., Jeong, Y. H., Yoon, S. L., and Seol, S.-Y. Diagnosis kits and method for detecting cancer using polymorphic minisatellite. (July 19, 2011) US Patent 7, 981,613.

82. Singh, R. and Bandyopadhyay, D. (2007) MUC1: a target molecule for cancer therapy. Cancer Biology & Therapy, 6(4), 481–486.

83. Yoon, S.-L., Roh, Y.-G., Chu, I.-S., Heo, J., Kim, S. I., Chang, H., Kang, T.-H., Chung, J. W., Koh, S. S., Larionov, V., et al. (2016) A polymorphic minisatellite region of BORIS regulates gene expression and its rare variants correlate with lung cancer susceptibility. Experimental & Molecular Medicine, 48(7), e246–e246.

84. Rose, A. M. Therapeutics and diagnostics based on minisatellite repeat element 1 (msr1). (December 17, 2015) US Patent App. 14/761,952.

85. Fondon, J. W. and Garner, H. R. (2004) Molecular origins of rapid and continuous morphological evolution. Proceedings of the National Academy of Sciences, 101(52), 18058–18063.

86. Laidlaw, J., Gelfand, Y., Ng, K.-W., Garner, H. R., Ranganathan, R., Benson, G., and Fondon III, J. W. (2007) Elevated basal slippage mutation rates among the Canidae. Journal of Heredity, 98(5), 452–460.

87. Sulovari, A., Li, R., Audano, P. A., Porubsky, D., Vollger, M. R., Logsdon, G. A., Warren, W. C., Pollen, A. A., Chaisson, M. J., Eichler, E. E., et al. (2019) Human-specific tandem repeat expansion and differential gene expression during primate evolution. Proceedings of the National Academy of Sciences, 116(46), 23243–23253.

88. Gymrek, M. (2017) A genomic view of short tandem repeats. Current Opinion in Genetics & Development, 44, 9–16.

89. Tørresen, O. K., Star, B., Mier, P., Andrade-Navarro, M. A., Bateman, A., Jarnot, P., Gruca, A., Grynberg, M., Kajava, A. V., Promponas, V. J., et al. (2019) Tandem repeats lead to sequence assembly errors and impose multi-level challenges for genome and protein databases. Nucleic Acids Research, 47(21), 10994–11006.

90. Gymrek, M., Golan, D., Rosset, S., and Erlich, Y. (2012) lobSTR: A short tandem repeat profiler for personal genomes. Genome Research,.

91. Kristmundsdóttir, S., Sigurpálsdóttir, B. D., Kehr, B., and Halldórsson, B. V. (2017) popSTR: population-scale detection of STR variants. Bioinformatics, 33(24), 4041–4048.

92. Willems, T., Zielinski, D., Yuan, J., Gordon, A., Gymrek, M., and Erlich, Y. (2017) Genome-wide profiling of heritable and extlessi extgreaterde novo extless/i extgreater STR variations. Nature Methods, 14(6), 590– 592.

93. Mousavi, N., Shleizer-Burko, S., Yanicky, R., and Gymrek, M. (2019) Profiling the genome-wide landscape of tandem repeat expansions. Nucleic Acids Research, 47(15), e90–e90.

94. Dolzhenko, E., Deshpande, V., Schlesinger, F., Krusche, P., Petrovski, R., Chen, S., Emig-Agius, D., Gross, A., Narzisi, G., Bowman, B., et al. (2019) ExpansionHunter: a sequence-graph-based tool to analyze variation in short tandem repeat regions. Bioinformatics, 35(22), 4754– 4756.

95. Bakhtiari, M., Shleizer-Burko, S., Gymrek, M., Bansal, V., and Bafna, V. (2018) Targeted genotyping of variable number tandem repeats with adVNTR. Genome Research, 28(11), 1709–1719.

96. Gelfand, Y., Hernandez, Y., Loving, J., and Benson, G. (2014) VNTRseek-a computational tool to detect tandem repeat variants in high-throughput sequencing data. Nucleic Acids Research, 42(14), 8884–8894.

97. Benson, G. (1999) Tandem repeats finder: a program to analyze DNA sequences. Nucleic Acids Research, 27(2), 573–580.

98. 1000 Genomes Project Consortium and others (2015) A global reference for human genetic variation. Nature, 526(7571), 68.

99. Zook, J. M., Catoe, D., McDaniel, J., Vang, L., Spies, N., Sidow, A., Weng, Z., Liu, Y., Mason, C. E., Alexander, N., et al. (2016) Extensive sequencing of seven human genomes to characterize benchmark reference materials. Scientific Data, 3(1), 1–26.

100. Chaisson, M. J. P., Huddleston, J., Dennis, M. Y., Sudmant, P. H., Malig, M., Hormozdiari, F., Antonacci, F., Surti, U., Sandstrom, R., Boitano, M., Landolin, J. M., Stamatoyannopoulos, J. A., Hunkapiller, M. W., Korlach, J., and Eichler, E. E. (2014) Resolving the complexity of the human genome using single-molecule sequencing. Nature, 517(7536), 608–611.

101. Huddleston, J., Chaisson, M. J. P., Steinberg, K. M., Warren, W., Hoekzema, K., Gordon, D., Graves-Lindsay, T. A., Munson, K. M., Kronenberg, Z. N., Vives, L., Peluso, P., Boitano, M., Chin, C.-S., Korlach, J., Wilson, R. K., and Eichler, E. E. (2017) Discovery and genotyping of structural variation from long-read haploid genome sequence data. Genome Research, 27(5), 677–685.

102. Drmanac, R., Sparks, A. B., Callow, M. J., Halpern, A. L., Burns, N. L., Kermani, B. G., Carnevali, P., Nazarenko, I., Nilsen, G. B., Yeung, G., Dahl, F., Fernandez, A., Staker, B., Pant, K. P., Baccash, J., Borcherding, P., Brownley, A., Cedeno, R., Chen, L., Chernikoff, D., Cheung, A., Chirita, R., Curson, B., Ebert, J. C., Hacker, C. R., Hartlage, R., Hauser, B., Huang, S., Jiang, Y., Karpinchyk, V., Koenig, M., Kong, C., Landers, T., Le, C., Liu, J., McBride, C. E., Morenzoni, M., Morey, R. E., Mutch, K., Perazich, H., Perry, K., Peters, B. A., Peterson, J., Pethiyagoda, C. L., Pothuraju, K., Richter, C., Rosenbaum, A. M., Roy, S., Shafto, J., Sharanhovich, U., Shannon, K. W., Sheppy, C. G., Sun, M., Thakuria, J. V., Tran, A., Vu, D., Zaranek, A. W., Wu, X., Drmanac, S., Oliphant, A. R., Banyai, W. C., Martin, B., Ballinger, D. G., Church, G. M., and Reid, C. A. (2010) Human Genome Sequencing Using Unchained Base Reads on Self-Assembling DNA Nanoarrays. Science, 327(5961), 78–81.

103. Wenger, A. M., Peluso, P., Rowell, W. J., Chang, P.-C., Hall, R. J., Concepcion, G. T., Ebler, J., Fungtammasan, A., Kolesnikov, A., Olson, N. D., et al. (2019) Accurate circular consensus long-read sequencing improves variant detection and assembly of a human genome. Nature Biotechnology, 37(10), 1155–1162.

104. Lander, E. S. and Waterman, M. S. (1988) Genomic mapping by fingerprinting random clones: A mathematical analysis. Genomics, 2(3), 231–239.

105. Gelfand, Y., Rodriguez, A., and Benson, G. (2007) TRDB—the tandem repeats database. Nucleic Acids Research, 35(suppl 1), D80–D87.

106. Ye, J., Coulouris, G., Zaretskaya, I., Cutcutache, I., Rozen, S., and Madden, T. L. (2012) Primer-BLAST: a tool to design target-specific primers for polymerase chain reaction. BMC Bioinformatics, 13(1), 134.

107. Kent, W. J., Sugnet, C. W., Furey, T. S., Roskin, K. M., Pringle, T. H., Zahler, A. M., and Haussler, D. (2002) The human genome browser at UCSC. Genome Research, 12(6), 996–1006.

108. Li, H. (2013) Aligning sequence reads, clone sequences and assembly contigs with BWA-MEM. arXiv Preprint, doi:10.6084/M9.FIGSHARE.963153.V1,.

109. Quinlan, A. R. and Hall, I. M. (2010) BEDTools: a flexible suite of utilities for comparing genomic features. Bioinformatics, 26(6), 841– 842.

110. Karolchik, D., Hinrichs, A. S., Furey, T. S., Roskin, K. M., Sugnet, C. W., Haussler, D., and Kent, W. J. (2004) The UCSC Table Browser data retrieval tool. Nucleic Acids Research, 32(suppl 1), D493–D496.

111. Hsu, F., Kent, W. J., Clawson, H., Kuhn, R. M., Diekhans, M., and Haussler, D. (2006) The UCSC known genes. Bioinformatics, 22(9), 1036–1046.

112. ENCODE Project Consortium and others (2012) An integrated encyclopedia of DNA elements in the human genome. Nature, 489(7414), 57–74.

113. Davis, C. A., Hitz, B. C., Sloan, C. A., Chan, E. T., Davidson, J. M., Gabdank, I., Hilton, J. A., Jain, K., Baymuradov, U. K., Narayanan, K., et al. (2018) The Encyclopedia of DNA elements (ENCODE): data portal update. Nucleic Acids Research, 46(D1), D794–D801.

114. Thurman, R. E., Rynes, E., Humbert, R., Vierstra, J., Maurano, M. T., Haugen, E., Sheffield, N. C., Stergachis, A. B., Wang, H., Vernot, B., et al. (2012) The accessible chromatin landscape of the human genome. Nature, 489(7414), 75–82.

115. Kundaje, A., Meuleman, W., Ernst, J., Bilenky, M., Yen, A., Heravi-Moussavi, A., Kheradpour, P., Zhang, Z., Wang, J., Ziller, M. J., et al. (2015) Integrative analysis of 111 reference human epigenomes. Nature, 518(7539), 317–330.

116. Gardiner-Garden, M. and Frommer, M. (1987) CpG islands in vertebrate genomes. Journal of Molecular Biology, 196(2), 261–282.

117. Sheffield, N. C. and Bock, C. (2016) LOLA: enrichment analysis for genomic region sets and regulatory elements in R and Bioconductor. Bioinformatics, 32(4), 587–589.

118. Subramanian, A., Tamayo, P., Mootha, V. K., Mukherjee, S., Ebert, B. L., Gillette, M. A., Paulovich, A., Pomeroy, S. L., Golub, T. R., Lander, E. S., et al. (2005) Gene set enrichment analysis: a knowledge-based approach for interpreting genome-wide expression profiles. Proceedings of the National Academy of Sciences, 102(43), 15545–15550.

119. Ashburner, M., Ball, C. A., Blake, J. A., Botstein, D., Butler, H., Cherry, J. M., Davis, A. P., Dolinski, K., Dwight, S. S., Eppig, J. T., et al. (2000) Gene ontology: tool for the unification of biology. Nature Genetics, 25(1), 25.

120. Kanehisa, M. and Goto, S. (2000) KEGG: kyoto encyclopedia of genes and genomes. Nucleic Acids Research, 28(1), 27–30.

121. Lappalainen, T., Sammeth, M., Friedländer, M. R., Ac’t Hoen, P., Monlong, J., Rivas, M. A., Gonzalez-Porta, M., Kurbatova, N., Griebel, T., Ferreira, P. G., et al. (2013) Transcriptome and genome sequencing uncovers functional variation in humans. Nature, 501(7468), 506–511.

122. Lee, D., Cheng, A., Lawlor, N., Bolisetty, M., and Ucar, D. (2018) Detection of correlated hidden factors from single cell transcriptomes using Iteratively Adjusted-SVA (IA-SVA). Scientific Reports, 8(1), 1–13.

123. Consortium, E. P. et al. (2004) The ENCODE (ENCyclopedia of DNA Elements) project. Science, 306(5696), 636–640.

124. Benjamini, Y. and Hochberg, Y. (1995) Controlling the false discovery rate: a practical and powerful approach to multiple testing. Journal of the Royal Statistical Society: Series B (Methodological), 57(1), 289–300.

125. Günther, T. and Nettelblad, C. (2019) The presence and impact of reference bias on population genomic studies of prehistoric human populations. PLoS Genetics, 15(7), e1008302.

126. Kent, W. J. and Haussler, D. (2001) Assembly of the working draft of the human genome with GigAssembler. Genome Research, 11(9), 1541– 1548.

127. Campbell, H., Carothers, A. D., Rudan, I., Hayward, C., Biloglav, Z., Barac, L., Pericic, M., Janicijevic, B., Smolej-Narancic, N., Polasek, O., et al. (2007) Effects of genome-wide heterozygosity on a range of biomedically relevant human quantitative traits. Human Molecular Genetics, 16(2), 233–241.

128. Herráez, D. L., Bauchet, M., Tang, K., Theunert, C., Pugach, I., Li, J., Nandineni, M. R., Gross, A., Scholz, M., and Stoneking, M. (2009) Genetic variation and recent positive selection in worldwide human populations: evidence from nearly 1 million SNPs. PLoS One, 4(11), e7888.

129. Bray, S. M., Mulle, J. G., Dodd, A. F., Pulver, A. E., Wooding, S., and Warren, S. T. (2010) Signatures of founder effects, admixture, and selection in the Ashkenazi Jewish population. Proceedings of the National Academy of Sciences, 107(37), 16222–16227.

130. Edea, Z., Bhuiyan, M., Dessie, T., Rothschild, M., Dadi, H., and Kim, K. (2015) Genome-wide genetic diversity, population structure and admixture analysis in African and Asian cattle breeds. Animal: an International Journal of Animal Bioscience, 9(2), 218.

131. Raczy, C., Petrovski, R., Saunders, C. T., Chorny, I., Kruglyak, S., Margulies, E. H., Chuang, H.-Y., Källberg, M., Kumar, S. A., Liao, A., et al. (2013) Isaac: ultra-fast whole-genome secondary analysis on Illumina sequencing platforms. Bioinformatics, 29(16), 2041–2043.

132. Lever, J., Zhao, E. Y., Grewal, J., Jones, M. R., and Jones, S. J. (2019) CancerMine: a literature-mined resource for drivers, oncogenes and tumor suppressors in cancer. Nature Methods, 16(6), 505–507.

133. Chambon, M., Orsetti, B., Berthe, M.-L., Bascoul-Mollevi, C., Rodriguez, C., Duong, V., Gleizes, M., Thénot, S., Bibeau, F., Theillet, C., et al. (2011) Prognostic significance of TRIM24/TIF-1α gene expression in breast cancer. The American Journal of Pathology, 178(4), 1461–1469.

134. Lu, G. M., Wang, S., Zhao, L., Zhou, X. Q., Yan, H., Wang, Q. C., Huang, J. F., et al. (2017) Expression of CD44, TRIM24, TAGLN-2, ER and PR in breast invasive ductal carcinoma and their clinicopathologic significance. Chinese Journal of Clinical and Experimental Pathology, 33(7), 724–727.

135. Pathiraja, T. N., Thakkar, K. N., Jiang, S., Stratton, S., Liu, Z., Gagea, M., Shi, X., Shah, P. K., Phan, L., Lee, M.-H., et al. (2015) TRIM24 links glucose metabolism with transformation of human mammary epithelial cells. Oncogene, 34(22), 2836–2845.

136. Balko, J. M., Cook, R. S., Vaught, D. B., Kuba, M. G., Miller, T. W., Bhola, N. E., Sanders, M. E., Granja-Ingram, N. M., Smith, J. J., Meszoely, I. M., et al. (2012) Profiling of residual breast cancers after neoadjuvant chemotherapy identifies DUSP4 deficiency as a mechanism of drug resistance. Nature Medicine, 18(7), 1052–1059.

137. Menyhart, O., Budczies, J., Munkácsy, G., Esteva, F. J., Szabó, A., Miquel, T. P., and Győrffy, B. (2017) DUSP4 is associated with increased resistance against anti-HER2 therapy in breast cancer. Oncotarget, 8(44), 77207.

138. Deng, T., Liu, J. C., Chung, P. E., Uehling, D., Aman, A., Joseph, B., Ketela, T., Jiang, Z., Schachter, N. F., Rottapel, R., et al. (2014) shRNA kinome screen identifies TBK1 as a therapeutic target for HER2+ breast cancer. Cancer Research, 74(7), 2119–2130.

139. Long, X., Shi, Y., Ye, P., Guo, J., Zhou, Q., and Tang, Y. (2020) MicroRNA-99a suppresses breast cancer progression by targeting FGFR3. Frontiers in Oncology, 9, 1473.

140. Cichon, S., Craddock, N., Daly, M., Faraone, S., Gejman, P., Kelsoe, J., Lehner, T., Levinson, D., Moran, A., Sklar, P., et al. (2009) Psychiatric GWAS Consortium Coordinating Committee Genomewide association studies: History, rationale, and prospects for psychiatric disorders. American Journal of Psychiatry, 166(5), 540–556.

141. Nagraj, V., Magee, N. E., and Sheffield, N. C. (2018) LOLAweb: a containerized web server for interactive genomic locus overlap enrichment analysis. Nucleic Acids Research, 46(W1), W194–W199.

142. Sheffield, N. C., Thurman, R. E., Song, L., Safi, A., Stamatoyannopoulos, J. A., Lenhard, B., Crawford, G. E., and Furey, T. S. (2013) Patterns of regulatory activity across diverse human cell types predict tissue identity, transcription factor binding, and long-range interactions. Genome Research, 23(5), 777–788.

143. Reiff, R. E., Ali, B. R., Baron, B., Yu, T. W., Ben-Salem, S., Coulter, M. E., Schubert, C. R., Hill, R. S., Akawi, N. A., Al-Younes, B., et al. (2014) METTL23, a transcriptional partner of GABPA, is essential for human cognition. Human Molecular Genetics, 23(13), 3456–3466.

144. Bernkopf, M., Webersinke, G., Tongsook, C., Koyani, C. N., Rafiq, M. A., Ayaz, M., Müller, D., Enzinger, C., Aslam, M., Naeem, F., et al. (2014) Disruption of the methyltransferase-like 23 gene METTL23 causes mild autosomal recessive intellectual disability. Human Molecular Genetics, 23(15), 4015–4023.

145. Yanagihara, T., Tomino, T., Uruno, T., and Fukui, Y. (2017) Thymic epithelial cell–specific deletion of JMJD6 reduces Aire protein expression and exacerbates disease development in a mouse model of autoimmune diabetes. Biochemical and Biophysical Research Communications, 489(1), 8–13.

146. Poulard, C., Rambaud, J., Lavergne, E., Jacquemetton, J., Renoir, J.-M., Trédan, O., Chabaud, S., Treilleux, I., Corbo, L., and Le Romancer, M. (2015) Role of JMJD6 in breast tumourigenesis. PLoS One, 10(5), e0126181.

147. Wong, M., Sun, Y., Xi, Z., Milazzo, G., Poulos, R. C., Bartenhagen, C., Bell, J. L., Mayoh, C., Ho, N., Tee, A. E., et al. (2019) JMJD6 is a tumorigenic factor and therapeutic target in neuroblastoma. Nature Communications, 10(1), 1–15.

148. Lalioti, M., Antonarakis, S., and Scott, H. S. (2003) The epilepsy, the protease inhibitor and the dodecamer: progressive myoclonus epilepsy, cystatin b and a 12-mer repeat expansion. Cytogenetic and Genome Research, 100(1-4), 213–223.

149. Sudmant, P. H., Rausch, T., Gardner, E. J., Handsaker, R. E., Abyzov, A., Huddleston, J., Zhang, Y., Ye, K., Jun, G., Fritz, M. H.-Y., et al. (2015) An integrated map of structural variation in 2,504 human genomes. Nature, 526(7571), 75–81.

150. Liberzon, A., Subramanian, A., Pinchback, R., Thorvaldsdóttir, H., Tamayo, P., and Mesirov, J. P. (2011) Molecular signatures database (MSigDB) 3.0. Bioinformatics, 27(12), 1739–1740.

151. Marinho, F. V. C., Pinto, G. R., Oliveira, T., Gomes, A., Lima, V., Ferreira-Fernandes, H., Rocha, K., Magalhães, F., Velasques, B., Ribeiro, P., et al. (2019) The SLC6A3 3-UTR VNTR and intron 8 VNTR polymorphisms association in the time estimation. Brain Structure and Function, 224(1), 253–262.

152. Schlüter, T., Winz, O., Henkel, K., Eggermann, T., Mohammadkhani-Shali, S., Dietrich, C., Heinzel, A., Decker, M., Cumming, P., Zerres, K., et al. (2016) MAOA-VNTR polymorphism modulates context-dependent dopamine release and aggressive behavior in males. NeuroImage, 125, 378–385.

153. Zammit, S., Jones, G., Jones, S. J., Norton, N., Sanders, R. D., Milham, C., McCarthy, G. M., Jones, L. A., Cardno, A. G., Gray, M., et al. (2004) Polymorphisms in the MAOA, MAOB, and COMT genes and aggressive behavior in schizophrenia. American Journal of Medical Genetics Part B: Neuropsychiatric Genetics, 128(1), 19–20.

154. Vernaleken, I., Schlüter, T., Eggermann, T., Roesch, F., Zerres, K., and Mottaghy, F. (2015) Effect of MAOA-VNTR Polymorphism on Aggression and Dopamine Release. Journal of Nuclear Medicine, 56(supplement 3), 300–300.

155. Mill, J., Asherson, P., Browes, C., D’Souza, U., and Craig, I. (2002) Expression of the dopamine transporter gene is regulated by the 3’ UTR VNTR: Evidence from brain and lymphocytes using quantitative RT-PCR. American Journal of Medical Genetics, 114(8), 975–979.

156. Diatchenko, L., Nackley, A. G., Tchivileva, I. E., Shabalina, S. A., and Maixner, W. (2007) Genetic architecture of human pain perception. Trends in Genetics, 23(12), 605–613.

157. Kang, A. M., Palmatier, M. A., and Kidd, K. K. (1999) Global variation of a 40-bp VNTR in the 3’-untranslated region of the dopamine transporter gene (SLC6A3). Biological Psychiatry, 46(2), 151–160.

158. Liu, C.-C., Tsai, C.-W., Deak, F., Rogers, J., Penuliar, M., Sung, Y. M., Maher, J. N., Fu, Y., Li, X., Xu, H., et al. (2014) Deficiency in LRP6-mediated Wnt signaling contributes to synaptic abnormalities and amyloid pathology in Alzheimer’s disease. Neuron, 84(1), 63–77.

159. Hayase, Y., Amano, S., Hashizume, K., Tominaga, T., Miyamoto, H., Kanno, Y., Ueno-Inoue, Y., Inoue, T., Yamada, M., Ogata, S., et al. (2020) Down syndrome cell adhesion molecule like-1 (DSCAML1) links the GABA system and seizure susceptibility. Acta Neuropathologica Communications, 8(1), 1–17.

160. Carapito, R., Ivanova, E. L., Morlon, A., Meng, L., Molitor, A., Erdmann, E., Kieffer, B., Pichot, A., Naegely, L., Kolmer, A., et al. (2019) ZMIZ1 variants cause a syndromic neurodevelopmental disorder. The American Journal of Human Genetics, 104(2), 319–330.

161. Gymrek, M., Willems, T., Guilmatre, A., Zeng, H., Markus, B., Georgiev, S., Daly, M. J., Price, A. L., Pritchard, J. K., Sharp, A. J., et al. (2016) Abundant contribution of short tandem repeats to gene expression variation in humans. Nature Genetics, 48(1), 22–29.

162. Fotsing, S. F., Margoliash, J., Wang, C., Saini, S., Yanicky, R., Shleizer-Burko, S., Goren, A., and Gymrek, M. (2019) The impact of short tandem repeat variation on gene expression. Nature Genetics, 51(11), 1652–1659.

163. Hormozdiari, F., Kostem, E., Kang, E. Y., Pasaniuc, B., and Eskin, E. (2014) Identifying causal variants at loci with multiple signals of association. Genetics, 198(2), 497–508.

