## Supplementary.pdf for "Genome-wide characterization of human minisatellite VNTRs: population-specific alleles and gene expression differences"

#### S1 Supplementary Files

A list of the data files included in Supplementary Materials is listed in the table below.

| **Supplementary filename** | **Description** | **No. of lines (w/o header)** |
|---|---|---|
| `Reference_TRs.txt` | TR reference loci with positions and whether they overlap with different classes of annotations | 191,286 |
| `Summary_of_results.txt` | Summary of VNTRseek runs per sample, sample information, total number of genotyped TRs and VNTRs, sensitivity, and number of heterozygous calls. | 2,800 |
| `Common VNTRs.txt` | List of common VNTRs and how many individuals they were observed in | 5,676 |
| `Private_VNTRs.txt` | List of private VNTRs and how many individuals they were observed in | 22,538 |
| `Major_genotypes.txt` | Major genotype for common VNTRs and number of individuals that observed the reference allele | 5,676 |
| `Superpopulation_VNTRs.txt` | Fisher exact test results to find population-specific VNTR alleles | 23,025 |
| `Population_specific_GO_BP.xlsx` | GO term enrichment of population specific VNTR loci by GSEA | 100 |
| `RNA_VNTRs.txt` | Association of mRNA expression in blood with nearby VNTR alleles for 445 individuals | 1,071 |
| `Experiment_primers.txt` | The primers designed for experimental validation | 13 |

### S2 Filtering TRs

#### S2.1 Identifying indistinguishable TRs in the reference set

An *indistinguishable* TR belongs to a family of genomically dispersed TRs which share highly similar patterns and flanking sequence and can therefore produce misleading genotype calls. Indistinguishable TRs were identified using the procedure described in Gelfand et. al., 2014 (1), *i.e.*, each TR array from the initially filtered reference set of 228,486 TR loci was treated as a single read and all such reads were mapped to the original unfiltered TR set using VNTRseek (1). Any TR which mapped to a locus other than its own was labeled indistinguishable, resulting in 37,200 TRs labeled in this way ($\sim$ 16.3%). Indistinguishable TRs were not removed from the reference set, but genotype calls in the output of VNTRseek were flagged if the locus was indistinguishable. In the output VCF files the filtering field marks these TRs as *SC*, meaning they did not pass the *Singleton Criterion* filter. Alleles from indistinguishable TR loci detected in a sample were filtered from that sample before further processing.

#### S2.2 Filtering reference singletons to reduce false positive VNTRs

Singleton TRs, in the reference set of 228,486 loci, were removed if they caused false positive VNTR calls using simulated data. The following procedure was followed. For each reference TR, a sliding window equal in size to a specified read length was used to generate reads. The leftmost window ended one base upstream ($-1$) of the TR array start position and the rightmost window began one base downstream ($+1$) of the TR array end position, with the window moving in increments of one base. The combined simulated reads for all TRs were mapped back to the reference set with VNTRseek. A Singleton TR locus was then removed if:

- it was the source of at least one read resulting in a VNTR call, either at its own locus or another locus; or
- at least one read drawn from a different locus resulted in a VNTR call for the TR.

The procedure was repeated for three separate read lengths, 100 bp, 150 bp, and 250 bp, to produce three separate reference sets (Table S1, available at `https://doi.org/10.5281/zenodo.4065850`).

| Read Length(bp) | Reference Set Size | Reference Singletons | Singletons Removed | Final Ref. Set | Expected Genotyped (%) |
|---|---|---|---|---|---|
| 100–101 | 228,486 | 191,286 | 1,704 | 226,782 | 153,293 (80.14%) |
| 148–150 | 228,486 | 191,286 | 1,976 | 226,510 | 168,742 (88.21%) |
| 250 | 228,486 | 191,286 | 4,812 | 223,674 | 177,864 (92.98%) |

**Table S1: Filtering out reference singletons to reduce false positive VNTRs.** The original reference set contained 228,486 TR loci, labeled as singleton or indistinguishable. Using simulated reads generated from the reference set, singleton TRs that were called as false positive VNTRs or those which generated reads leading to such a result were removed (see S2.2). The "Expected Genotyped" column is the number of singleton TR loci for which the sum of array length and minimum flank lengths did not exceed the read length (for the 100/101 bp set, 100 bp was used as read length, for the 148/150 bp set, 150 bp was used). Percent is the Expected Genotyped out of all the original Reference Singletons.

#### S2.3 Filtering *multi* VNTR loci from sample results

VNTR loci which reported more alleles in a diploid sample than the expected number of chromosomes were termed *multis* in that sample. They correspond to loci with:

- three or more alleles on an autosomal chromosome,
- three or more alleles on chromosome X of a female individual,
- any allele on chromosome Y of a female individual, or
- two or more alleles on a sex chromosome of a male individual

For the two haploid samples, any locus that reported more than one allele, or any Y chromosome locus that reported any allele was termed multi.

#### S3 Heterozygous VNTR ratios by population

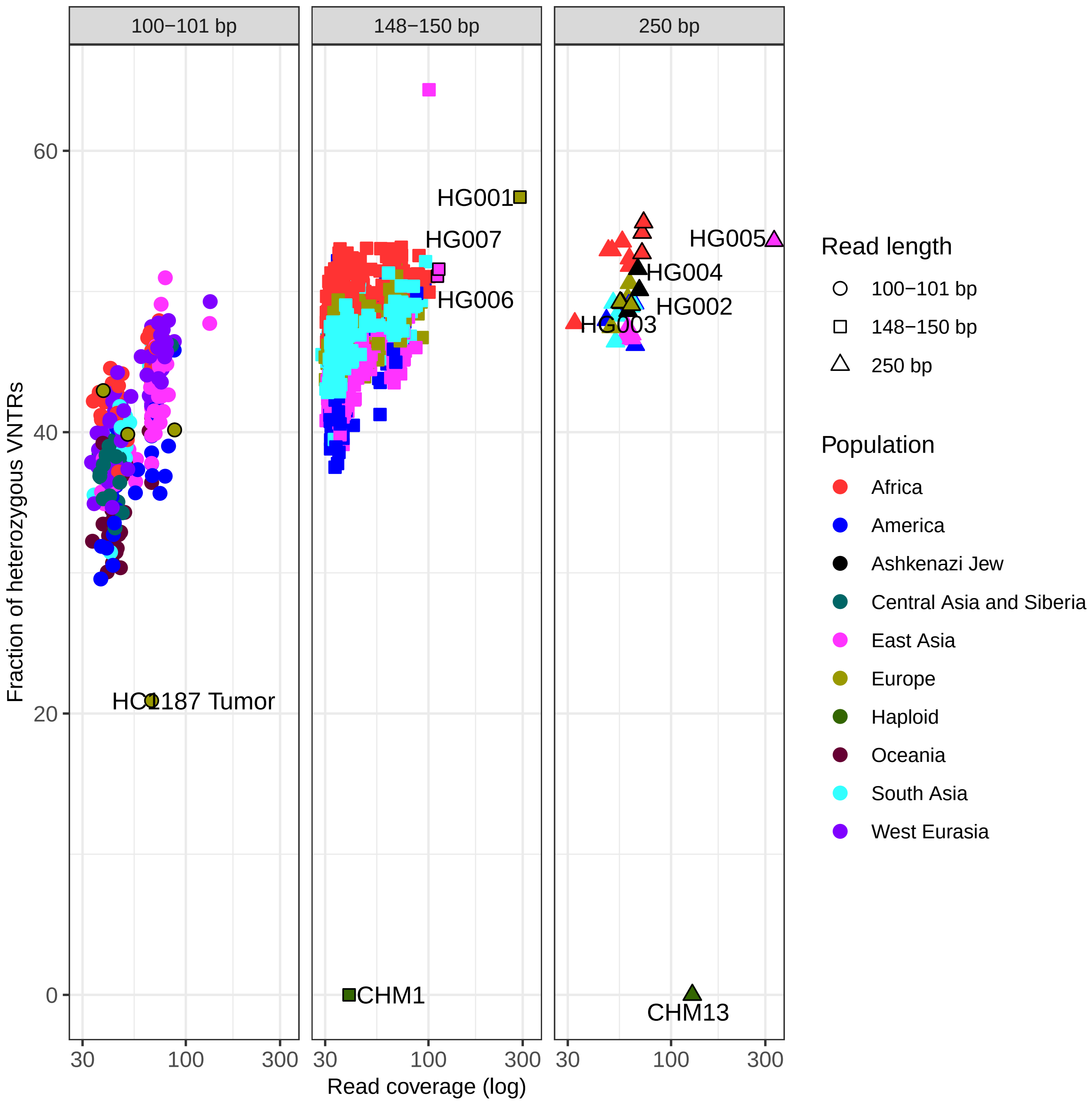


**Figure S1: Fraction per sample of VNTR loci called as heterozygous.** Samples are separated by read length. Higher read length and coverage provide more statistical power to detect heterozygous calls. Stratification by population is evident and is further displayed in Supplementary Figures S2, S3, and S4.

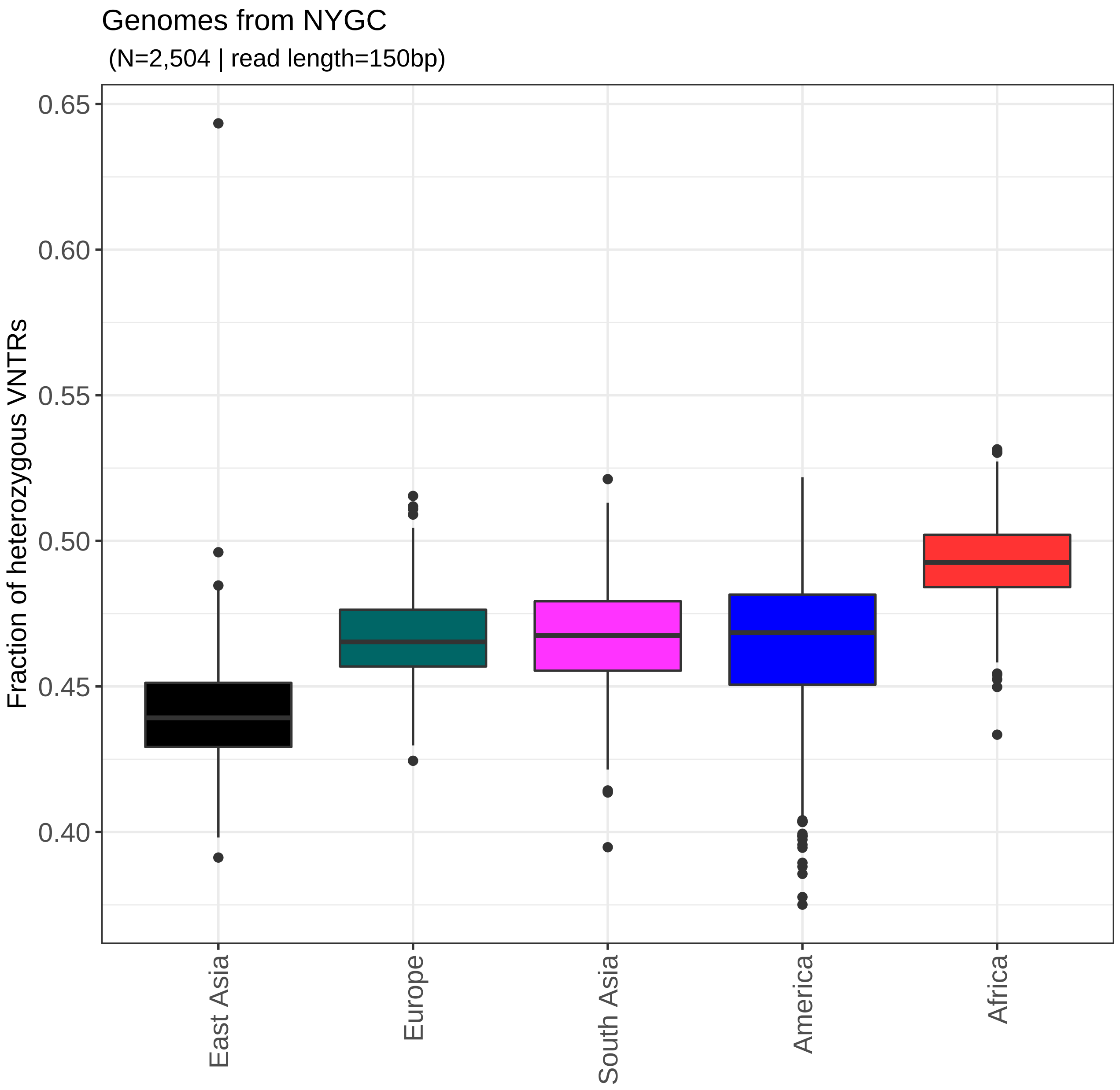


**Figure S2: Heterozygous VNTR calls on the 2,504 NYGC 150 bp samples per superpopulation.** Data are presented as box plots for each superpopulation showing the interquartile range and median (middle line). Africans had the highest percentages of heterozygous calls and East Asians had the lowest compared to the other super-populations.

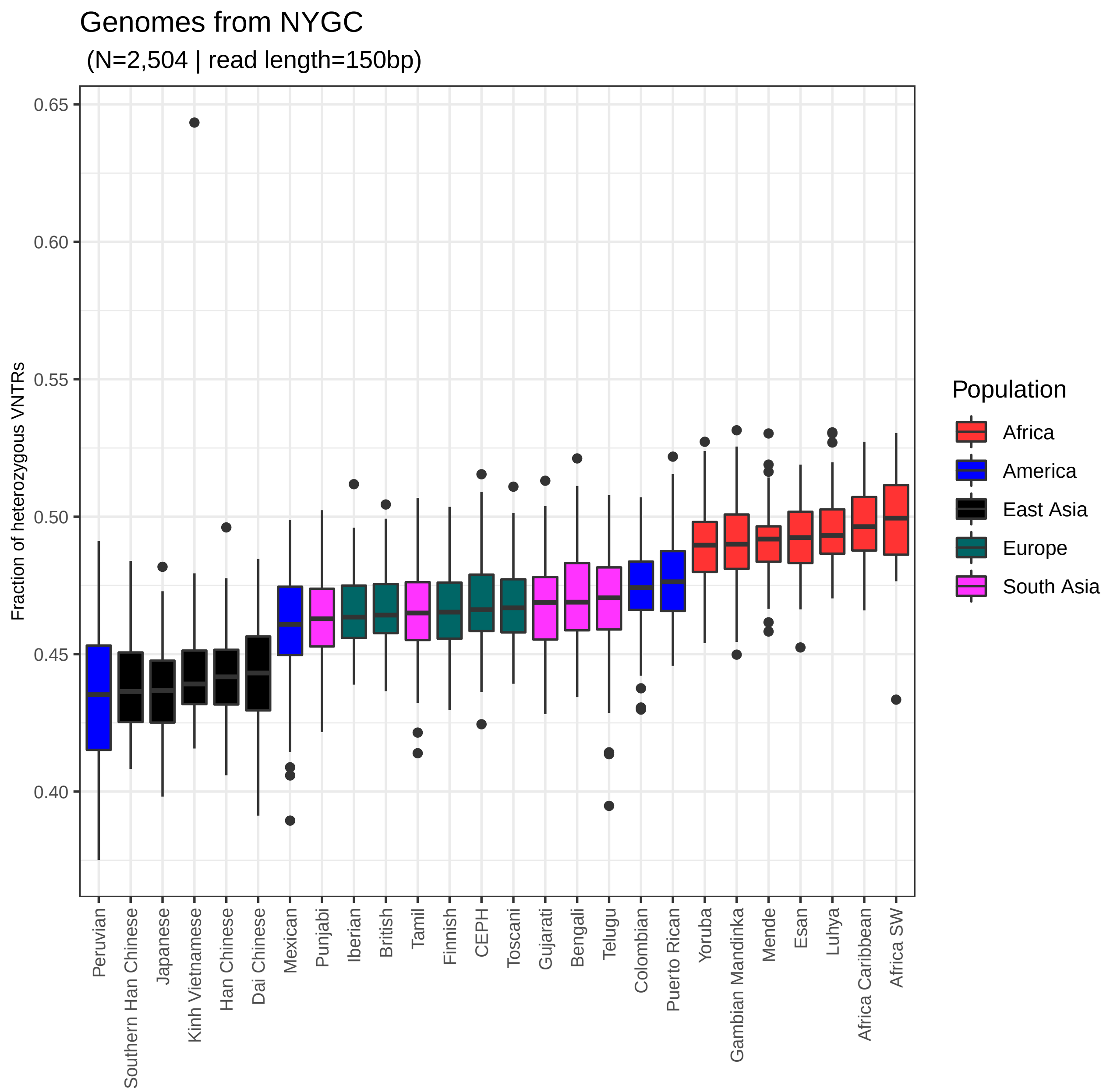


**Figure S3: Heterozygous VNTR calls on the 2,504 NYGC 150 bp samples per subpopulation.** Data are presented as box plots for each subpopulation showing the interquartile range and median (middle line). Among the admixed Americas populations, Peruvians had the lowest median percentage of heterozygous calls, similar to that of East Asians. The Peruvian population seems to have had the least mixing with African and European genomes in the NYGC dataset, based on this measure. South Asian and European sub-populations had similar frequencies of heterozygous calls.

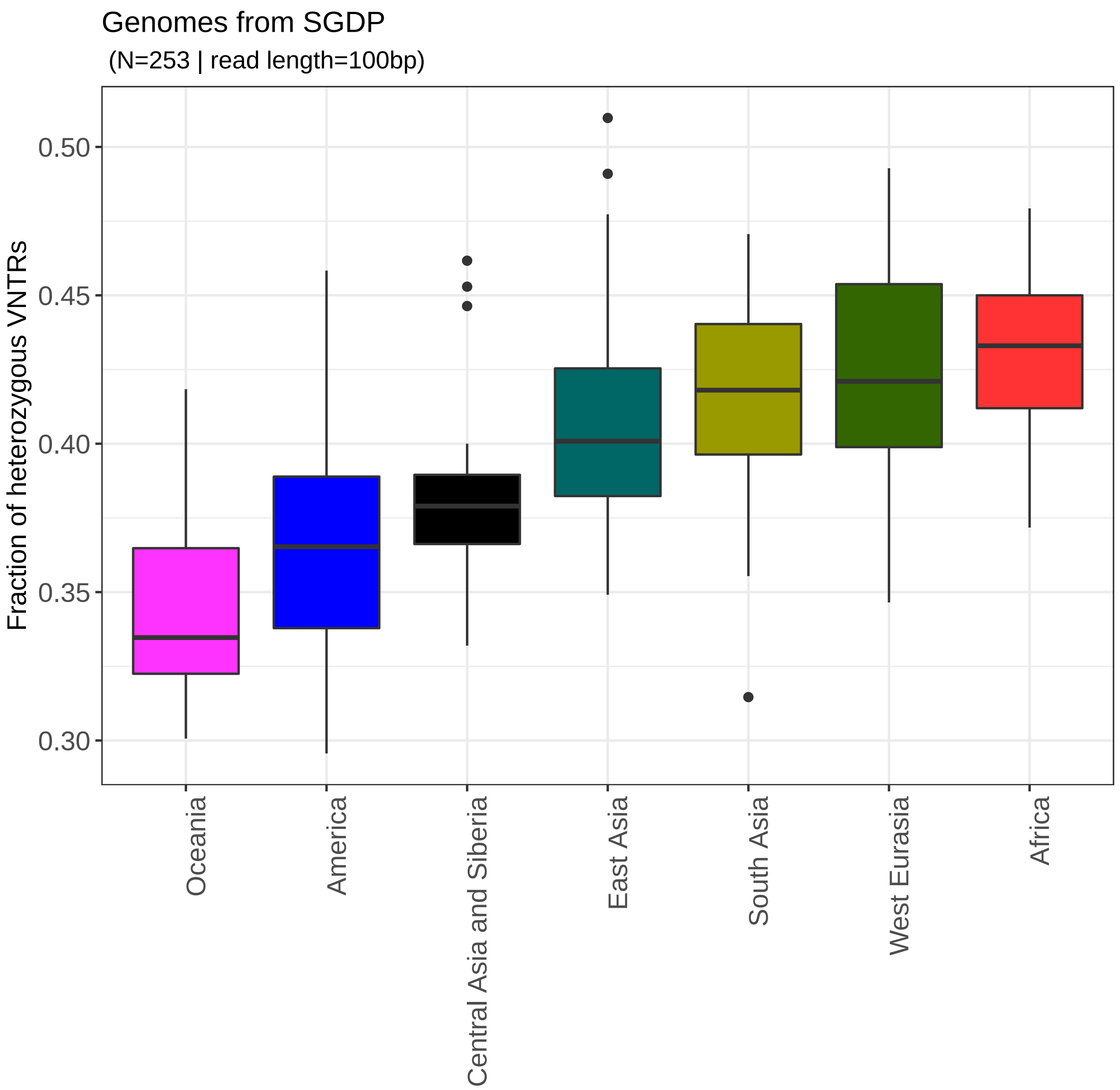


**Figure S4: Heterozygous VNTR calls on the 253 SGDP 100 bp samples per superpopulation.** Data are presented as box plots for each superpopulation showing the interquartile range and median (middle line). The SGDP aimed to sequence underrepresented populations. Although the overall percentage of heterozygous calls was lower than for the NYGC samples due to lower sensitivity with shorter reads, Africans still had the highest percentage of heterozygous calls. Interestingly genomes with Oceania ancestry had a very low percentage of heterozygous calls.

#### S3.1 Loss of heterozygosity in tumor samples

| Genotype | | | | |
|---|---|---|---|---|
| **Normal** | **Tumor** | **HC1187** | **HC2218** | **Description** |
| AA | AA | 391 | 349 | No change |
| | — | 75 | 32 | Loss of both alleles in tumor |
| | AB | 9 | 31 | Allele mutation in tumor, or failure to detect in normal |
| | BB | 2 | 1 | Allele mutation in tumor |
| AB | AB | 109 | 232 | No change |
| | — | 116 | 39 | Loss of both alleles in tumor |
| | AA | 90 | 40 | Loss of one allele (LOH) in tumor |
| | AC | 1 | 0 | Allele mutation in tumor |
| — | AA | 103 | 127 | Failure to detect in normal |
| — | AB | 36 | 84 | Failure to detect in normal |

**Table S2: Comparison of VNTR alleles in paired normal and tumor samples.** Notation: AA $\rightarrow$ homozygous: AB, AC $\rightarrow$ heterozygous; — $\rightarrow$ not detected. Due to significantly higher coverage in the tumor samples, we assumed the tumor genotyping was likely correct, whereas genotyping in the normal tissue may have failed to detect one or two alleles. The majority of VNTR loci detected in both tissues for each patient exhibited no change. The most common genotype change was loss in the tumor of one allele (LOH) or both alleles. Allele mutation was apparently uncommon.

| Sample | Coverage (×) | Multis | Total TRs | VNTRs | Heterozygous VNTRs | Ratio (%) |
|---|---|---|---|---|---|---|
| HC1187 Normal | 51 | 43 | 146,794 | 793 | 302 | 0.38 |
| HC1187 Tumor | 67 | 60 | 146,969 | 741 | 144 | 0.19 |
| HC2218 Normal | 38 | 40 | 144,710 | 724 | 293 | 0.40 |
| HC2218 Tumor | 88 | 57 | 147,812 | 864 | 328 | 0.37 |

**Table S3: Ratio of heterozygous VNTRs in paired normal and tumor samples**. The ratio of heterozygous calls in the tumor samples was lower than the paired normal tissue, suggesting potential loss of heterozygosity. In HC1187 the percentage was half that of the normal tissue. With same read length and higher coverage in the tumor samples, this finding cannot likely be attributed to artifacts.

### S4 VNTR alleles: Gain vs. Loss

**(a)**

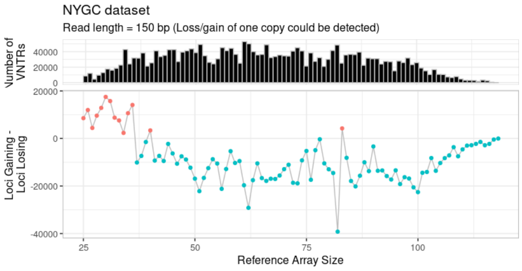


**(b)**

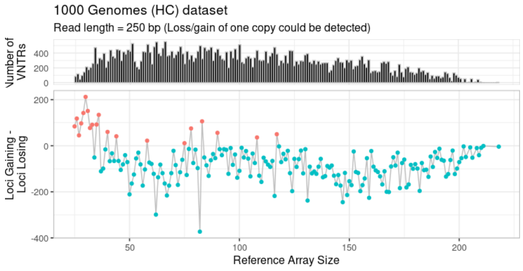


**(c)**

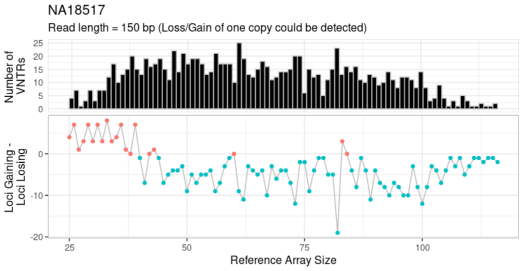


**(d)**

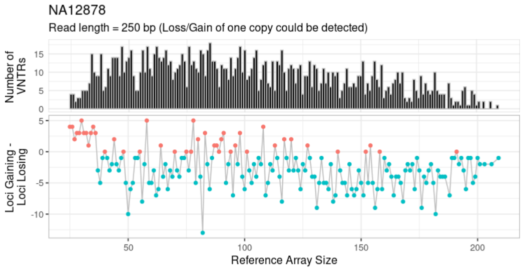


**(e)**

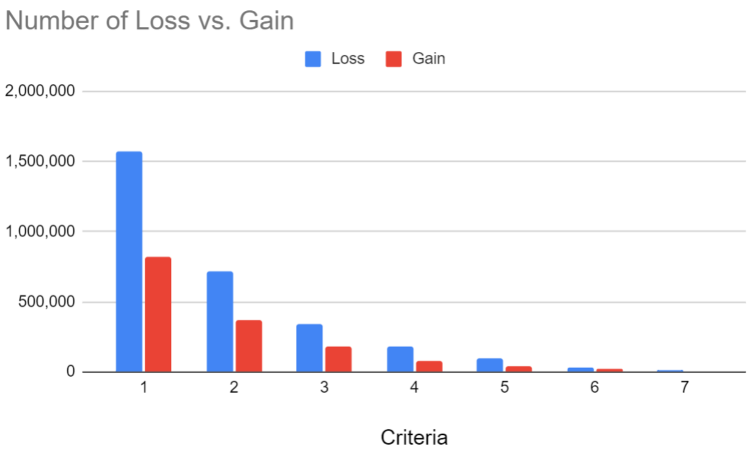


**Figure S5: Copy gain versus loss relative to the reference.** In the upper quartet of graphs, the upper subgraph is a histogram showing the number of non-reference alleles detected for those loci where both a gain and loss of one copy could be observed. Bin sizes are 1 bp. The lower subgraph counts, for each reference array length, the number of gain alleles (increase in copy number) minus the number of loss alleles (decrease in copy number). Negative numbers (aqua) indicate more loss than gain. (a) the entire NYGC dataset, (b) the entire 1000 Genomes dataset, (c) NA18517 (150 bp), (d) NA12878 (250 bp), which is labeled HG001 in GIAB. Graph (e) shows, the aggregate excess of loss over gain for the entire NYGC dataset, when considering loci for which both gain and loss of 1, 2, 3, etc. copies could be observed.

#### S5 Annotation of common and private VNTRs

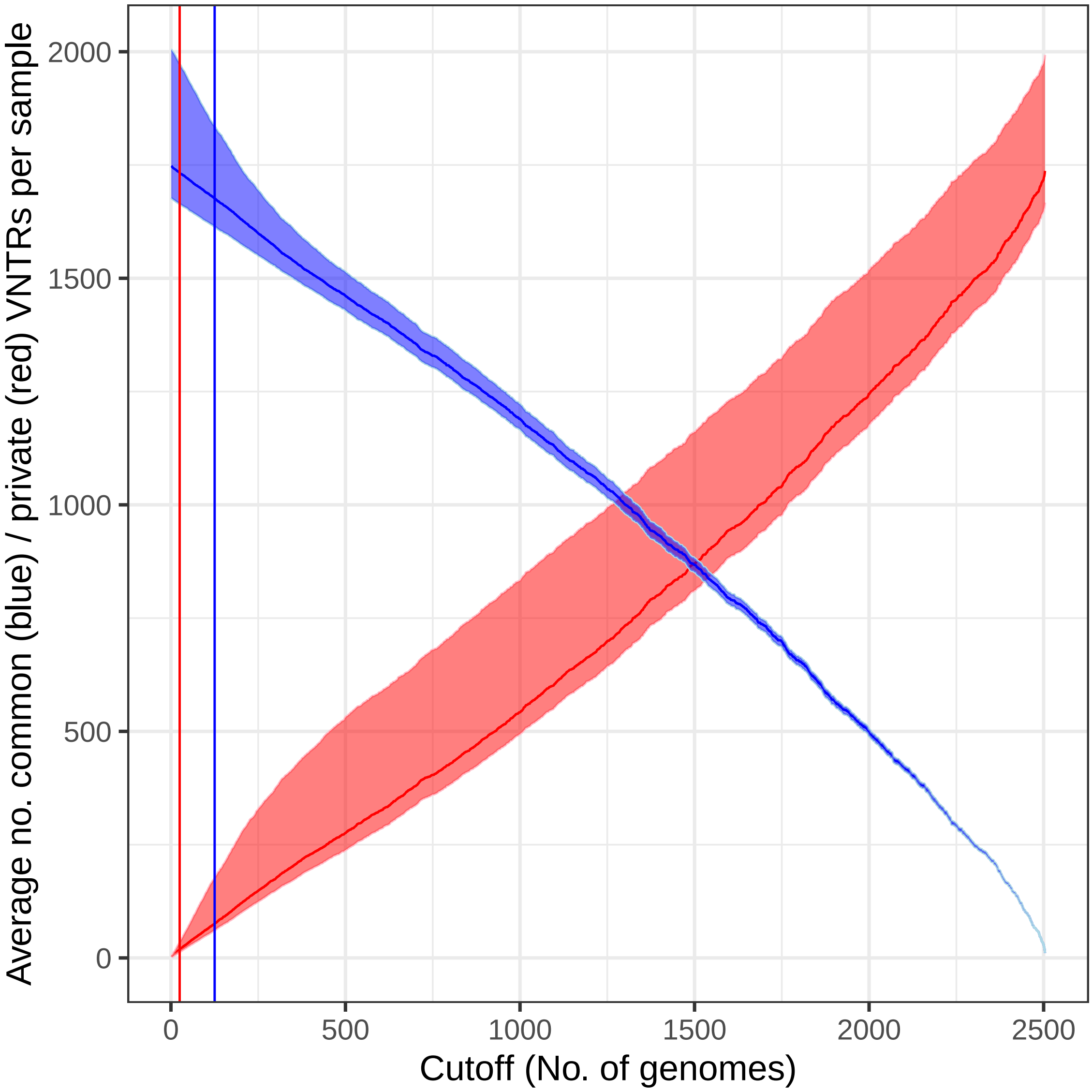


**Figure S6: Number of common/private VNTRs by cutoff in the 2,504 NYGC samples.** Blue area specifies the interquartile range (line is the median) of common VNTR loci counts per sample as the number of samples required to be called common increases from zero to 2,504. For example, at the 5% cutoff (126 - blue vertical line), there were 1,783 common VNTRs on average per genome (median 1,677). The pink area specifies the interquartile range of private VNTRs. At the 1% cutoff (25 - red vertical line), each genome had 46 private VNTRs on average (median 17). The graph shows that common VNTR loci were indeed very common since the numbers do not drop dramatically even if the cutoff were raised to 500 samples.

##### S5.1 Enrichment of common VNTRs by genomic annotation

| Feature | Reference TRs | All VNTRs | Common (>5%) | Private (<1%) |
|---|---|---|---|---|
| Total | 191,286 | 33,403 | 5,676 | 22,538 |
| Upstream (1Kb) | 12,415 | 3,424 | 671 | 2,181 |
| 5' UTR | 4,994 | 1,294 | 236 | 847 |
| Intron | 116,002 | 20,266 | 3,451 | 13,526 |
| Coding exon | 2,990 | 699 | 91 | 500 |
| 3' UTR | 6,628 | 1,295 | 238 | 817 |
| Downstream (1Kb) | 10,844 | 2,200 | 401 | 1,385 |
| Gene | 121,205 | 21,410 | 3,624 | 14,351 |
| TFBS cluster | 71,779 | 16,220 | 2,913 | 10,459 |
| DNAse cluster | 40,517 | 9,296 | 1,613 | 6,003 |
| CpG Island | 6,718 | 2,989 | 638 | 1,757 |

**Table S4: Annotation and enrichment of VNTRs.** Column *Reference TRs* shows the genomic feature annotations of the reference VNTRs. Numbers do not add to the total due to multiple classifications. We performed a Fisher's Exact Test to find enrichment of all VNTRs against all TRs and common/private VNTRs against all VNTRs. Significant p-values at the 5% threshold are presented in colored font, with blue and red indicating odds ratios less than one and greater than one, respectively. Compared to reference TRs, VNTRs were enriched in genes, gene upstream and downstream regions, TF binding sites, CpG islands, and open DNAse sites. Common VNTRs were more likely to occur in gene upstream regions, TF binding sites, and CpG islands (suggesting possible gene regulation effects); while they were less likely to occur inside exons, possibly due to disruption of protein product function. Private VNTRs, on the other hand, were less likely to occur at gene upstream and downstream regions, TF binding sites, open DNAse sites, and CpG islands (possibly due to the potential to randomly disrupt gene expression).

| Feature | Reference TRs | All VNTRs | Common (>5%) | Private (<1%) |
|---|---|---|---|---|
| Total | 191,286 | 33,403 | 5,676 | 22,538 |
| Upstream (1Kb) | 6.49 | 10.25 | 11.82 | 9.68 |
| 5' UTR | 2.61 | 3.87 | 4.16 | 3.76 |
| Intron | 60.64 | 60.67 | 60.80 | 60.01 |
| Coding exon | 1.56 | 2.09 | 1.60 | 2.22 |
| 3' UTR | 3.46 | 3.88 | 4.19 | 3.62 |
| Downstream (1Kb) | 5.67 | 6.59 | 7.06 | 6.15 |
| Gene | 63.36 | 64.10 | 63.85 | 63.67 |
| TFBS cluster | 37.52 | 48.56 | 51.32 | 46.41 |
| DNAse cluster | 21.18 | 27.83 | 28.42 | 26.64 |
| CpG Island | 3.51 | 8.95 | 11.24 | 7.80 |

**Table S5: Annotation and enrichment of VNTRs as percentages.** This table presents the enrichment data in Table S4 as percentages. We performed a Fisher's Exact Test to find enrichment of all VNTRs against all TRs and common/private VNTRs against all VNTRs. Significant p-values at the 5% threshold are presented in colored font, with blue and red indicating odds ratios less than one and greater than one, respectively.

#### S5.2 Genomic enrichment of common VNTRs by LOLAweb (2)

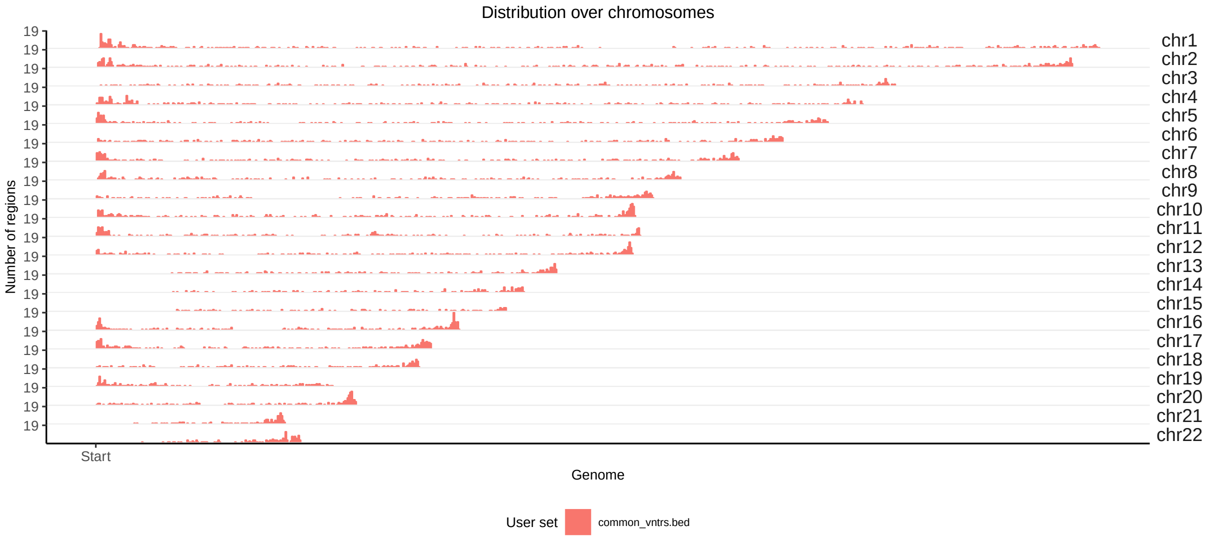


**(a)** Genomic distribution.

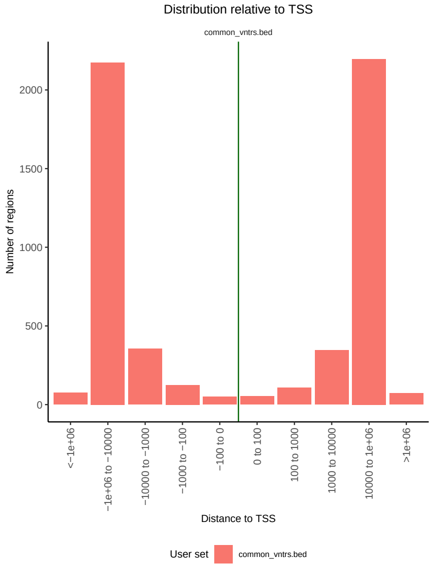


**(b)** TSS distribution.

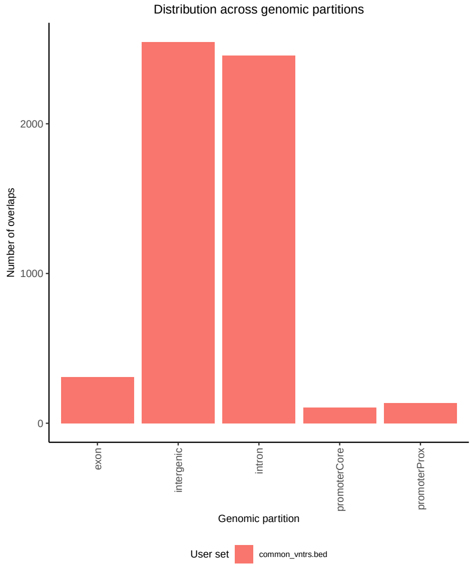


**(c)** Genomic partitions.

**Figure S7:** Genomic distributions of common VNTRs by LOLAweb.

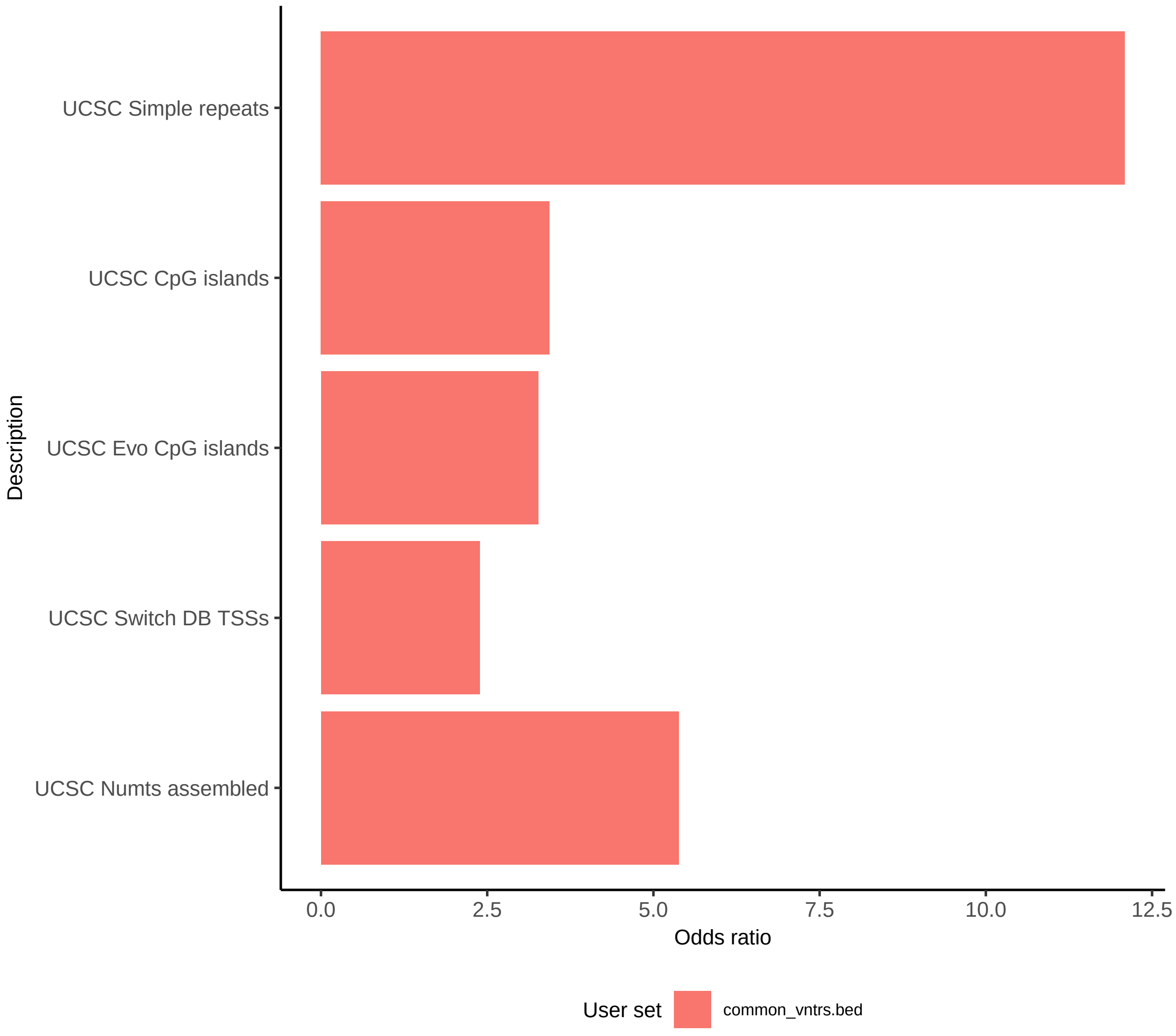


**Figure S8:** UCSC features enrichment of common VNTRs by LOLAweb.

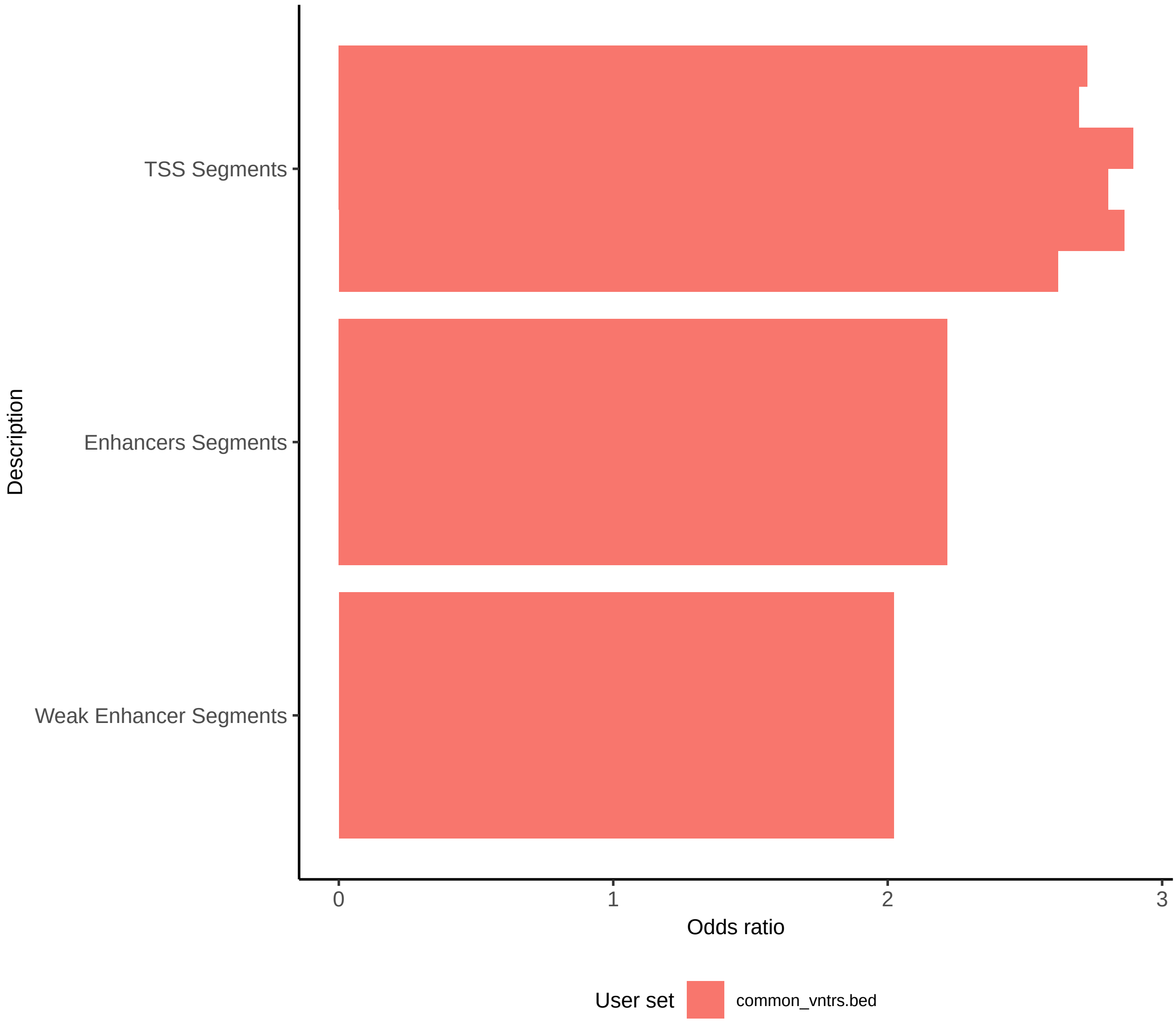


**Figure S9:** Genomic enrichment of common VNTRs in Encode segmentation by LOLAweb

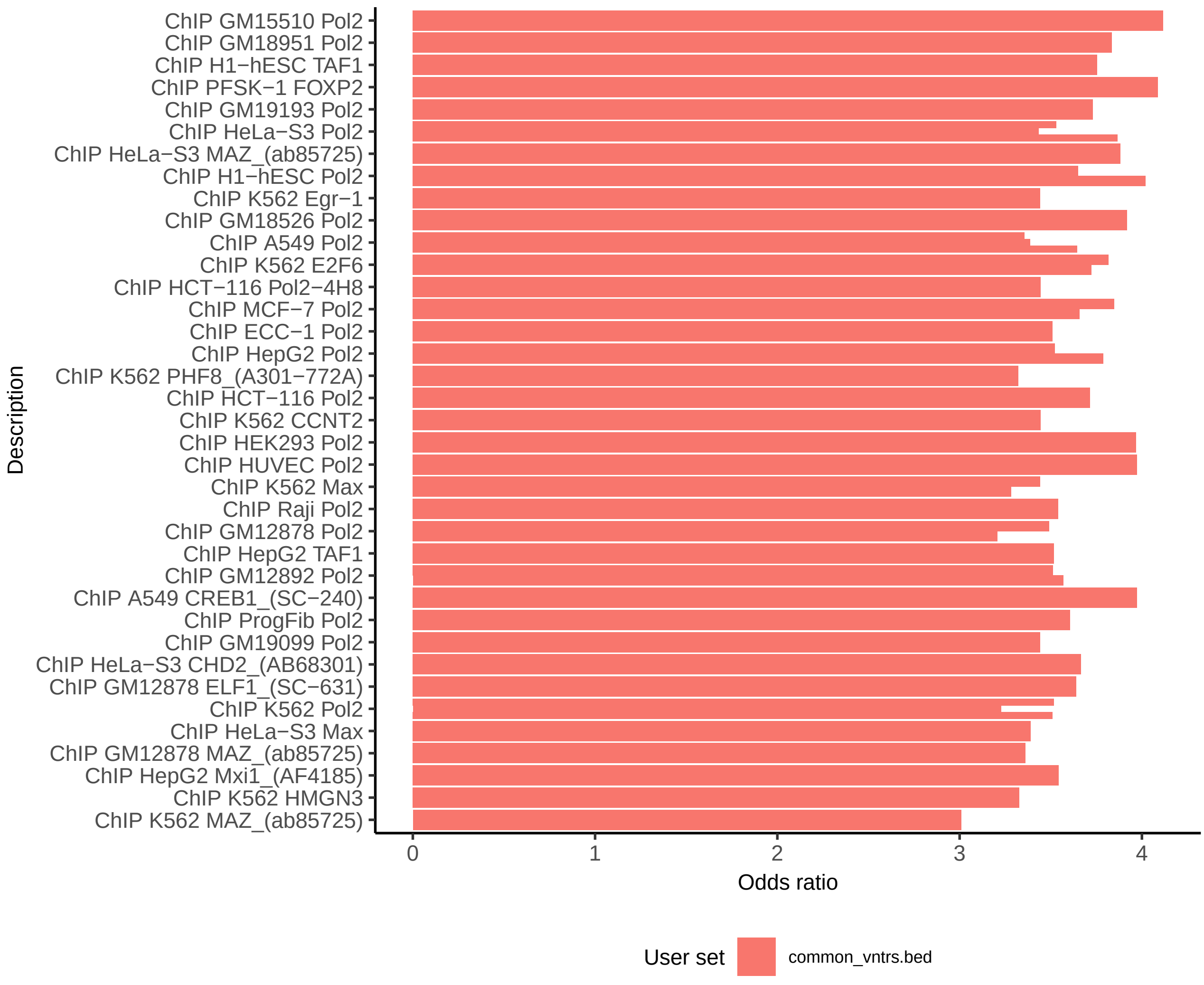


**Figure S10:** Genomic enrichment of common VNTRs in Encode TFBS by LOLAweb

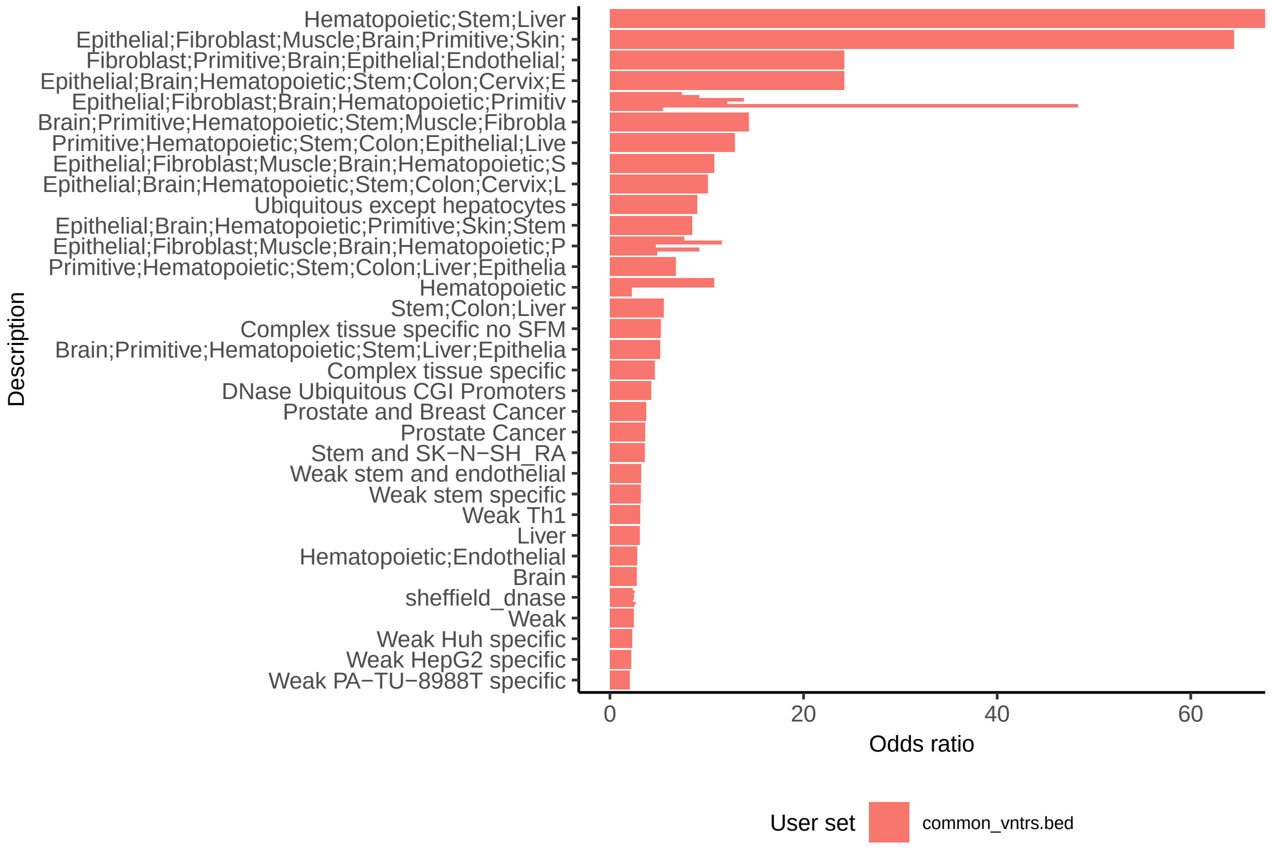


**Figure S11:** Genomic enrichment of common VNTRs in DNAse regions by tissue type using LOLAweb (3).

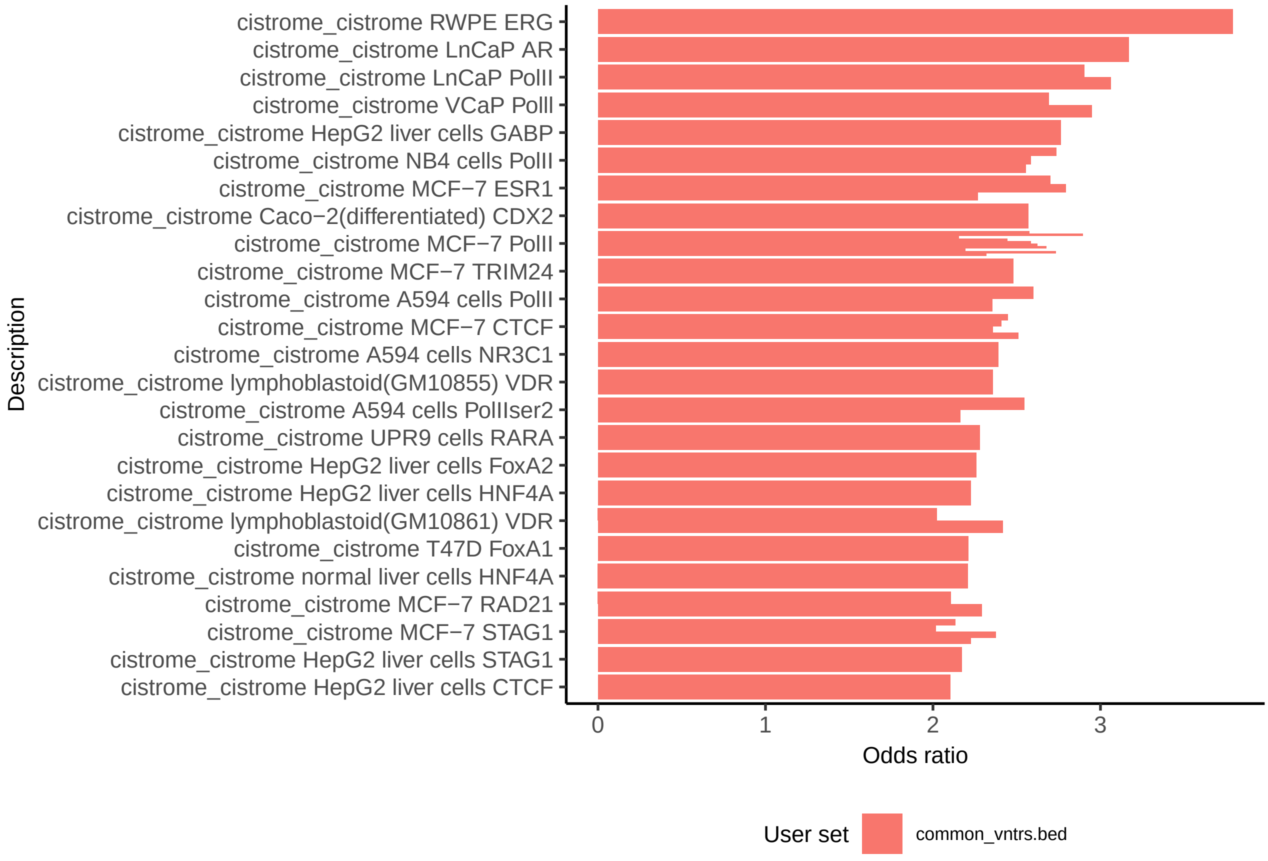


**Figure S12:** Genomic enrichment of common VNTRs in cistrome-cistrome by LOLAweb

### S6 Accuracy of VNTR predictions

#### S6.1 Experimental Validation

| # | TR id | Position (GRCh38) | Pattern size | Ref. copy no. | Array size | Description |
|---|---|---|---|---|---|---|
| 1* | 182316181 | chr10:132,254,409-132,254,809 | 105 | 3.8 | 401 | intron 1 of STK32C |
| 2 | 182316985 | chr10:133,004,939-133,005,092 | 27 | 5.8 | 154 | regulatory region targeting LINC01168 |
| 3 | 182453735 | chr13:95,693,631-95,693,714 | 30 | 2.8 | 84 | intron 1 of DNAJC3 |
| 4 | 182461997 | chr13:114,098,935-114,099,206 | 38 | 7.2 | 272 | intron 1 of RASA3 |
| 5 | 182493720 | chr14:100,557,003-100,557,217 | 70 | 3.1 | 215 | intron 1 of BEGAIN |
| 6 | 182515357 | chr15:66,039,739-66,040,018 | 34 | 8.2 | 280 | intron 5 of MEGF11 |
| 7 | 182608886 | chr17:80,918,399-80,918,567 | 27 | 6.3 | 169 | intron 21 of RPTOR |
| 8 | 182620950 | chr18:32,130,973-32,131,116 | 48 | 3.0 | 144 | exon 8 of RNF138 |
| 9 | 182982510 | chr5:1,085,607-1,085,746 | 34 | 4.1 | 140 | intron 6 of SLC12A7 |
| 10♯ | 183046759 | chr5:176,371,499-176,371,655 | 38 | 4.1 | 157 | intron 3 of ARL10 |
| 11† | 183081195 | chr6:81,752,006-81,752,072 | 15 | 4.5 | 67 | exon 2 of TENT5A |
| 12 | 183117043 | chr7:2,493,950-2,494,106 | 17 | 9.2 | 157 | regulatory region targeting MRM2, LFNG |
| 13 | 183169331 | chr7:128,947,291-128,947,357 | 15 | 4.5 | 67 | exon 6 of IRF5 |

| # | TR id | VNTRseek predictions | | | Expected bands | | | Validation bands | | |
|---|---|---|---|---|---|---|---|---|---|---|
| | | Child | Father | Mother | Child | Father | Mother | Child | Father | Mother |
| 1* | 182316181 | -2 | -2 | -2 | 237, 342 | 237 | 237 | -2,**-1** | -2,**-1** | -2, **+1** |
| 2 | 182316985 | -3, 0 | -1, 0 | -3, -2 | 118, 199 | 172, 199 | 118, 145 | yes | yes | yes |
| 3 | 182453735 | 0, +1 | 0, +1 | 0, +1 | 180, 210 | 180, 210 | 180, 210 | yes | yes | yes |
| 4 | 182461997 | -5, -4 | -5, -4 | -5, -2 | 193, 231 | 193, 231 | 193, 307 | yes | yes | yes |
| 5 | 182493720 | 0 | 0 | 0, -1 | 266 | 266 | 266, 196 | yes | yes | yes |
| 6 | 182515357 | -5 | -5, -6 | -5 | 195 | 195, 161 | 195 | yes | yes | yes |
| 7 | 182608886 | 0, -2 | 0, -3 | -2, +1 | 208, 181 | 208, 154 | 235, 181 | yes | yes | yes |
| 8 | 182620950 | 0, -1 | 0, -1 | 0, -1 | 243, 195 | 243, 195 | 243, 195 | yes | yes | yes |
| 9 | 182982510 | 0, -1 | 0 | 0, -1 | 301, 267 | 301 | 301, 267 | yes | yes | yes |
| 10♯ | 183046759 | 0 | 0, -1 | -1 | 222, 260 | 222, 184 | 184 | 0, **+1** | yes | -1, **+1** |
| 11† | 183081195 | +2, -1 | +2, +1 | +2, -1 | 182, 122 | 182, 167 | 182, 122 | yes | +2 | yes |
| 12 | 183117043 | -5, -3 | -5, -3 | 0, -3 | 180, 214 | 180, 214 | 265, 214 | yes | yes | yes |
| 13 | 183169331 | -2 | -2, 0 | -2 | 209 | 209, 239 | 209 | yes | yes | yes |

**Table S6: Experimental validation results:** Thirteen VNTR loci were selected for experimental validation in the AJ trio. All but one of the 66 bands predicted by VNTRseek were validated. †For the remaining band, the results were questionable because the two predicted alleles for the father were only 15 nucleotides different in length, which was too close to distinguish in the image. *For all three individuals, the gel contained bands (**bold**) not predicted (or detectable) by VNTRseek. The extra band for the son corresponded to the -1 allele as found in the PacBio reads. The father's extra band appeared to match with the -1 allele. The mother's extra band appeared to be a +1 allele (552 nucleotides). ♯An extra band for the mother and son (**bold**) was not predicted by VNTRseek, although it seemed to match the +1 allele that was detectable.

| Locus | Pattern Size | TR | VNTRseek | | | Expected bands | | | Validated | | |
|---|---|---|---|---|---|---|---|---|---|---|---|
| | | | son | father | mother | son | father | mother | son | father | mother |
| 1 | 105 | 182316181 | -2 | -2 | -2 | 237 | 237 | 237 | -2, **-1** | -2, **-1** | -2, **+1** |
| 2 | 27 | 182316985 | -3, 0 | -1, 0 | -3, -2 | 118, 199 | 172, 199 | 118, 145 | yes | yes | yes |
| 3 | 30 | 182453735 | 0, +1 | 0, +1 | 0, +1 | 180, 210 | 180, 210 | 180, 210 | yes | yes | yes |

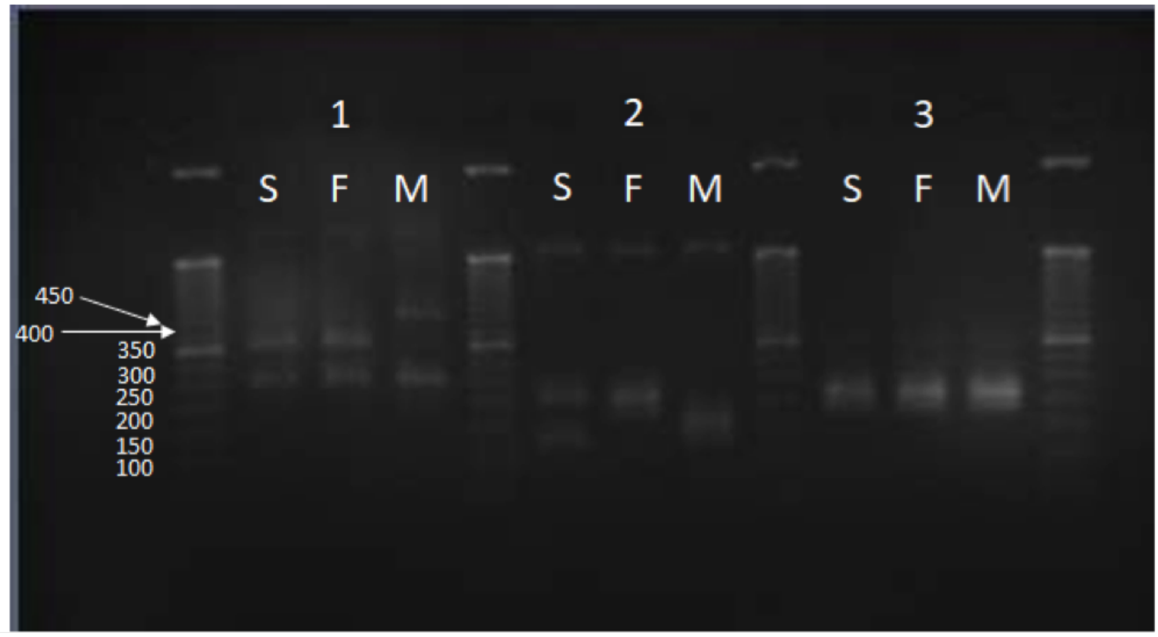


**Figure S13:** In locus 1, all three individuals share a band at 237bp. The father and son share a band at approximately 342bp and the mother has an additional band above 450bp. The bands at and above 342bp correspond to alleles that are not detectable by VNTRseek and were not predicted. The 342 bp band corresponds to a -1 allele found in the PacBio reads for the son. The mother's extra band appears to be a +1 allele (552 bp). In locus 2, the son and father share a band at 199 bp and the son and mother share a band at 118 bp. The father has an additional band at 172 bp and the mother at 145 bp. In locus 3, all individuals (son, father, mother) share the same bands, 180 and 210 bp). All three loci confirm the VNTRseek predictions.

| Locus | Pattern Size | TR | VNTRseek | | | Expected bands | | | Validated | | |
|---|---|---|---|---|---|---|---|---|---|---|---|
| | | | son | father | mother | son | father | mother | son | father | mother |
| 4 | 38 | 182461997 | -5,-4 | -5,-4 | -5,-2 | 193, 231 | 193, 231 | 193, 307 | yes | yes | yes |
| 5 | 70 | 182493720 | 0 | 0 | 0, -1 | 266 | 266 | 266, 196 | yes | yes | yes |
| 6 | 34 | 182515357 | -5 | -5, -6 | -5 | 195 | 195, 161 | 195 | yes | yes | yes |

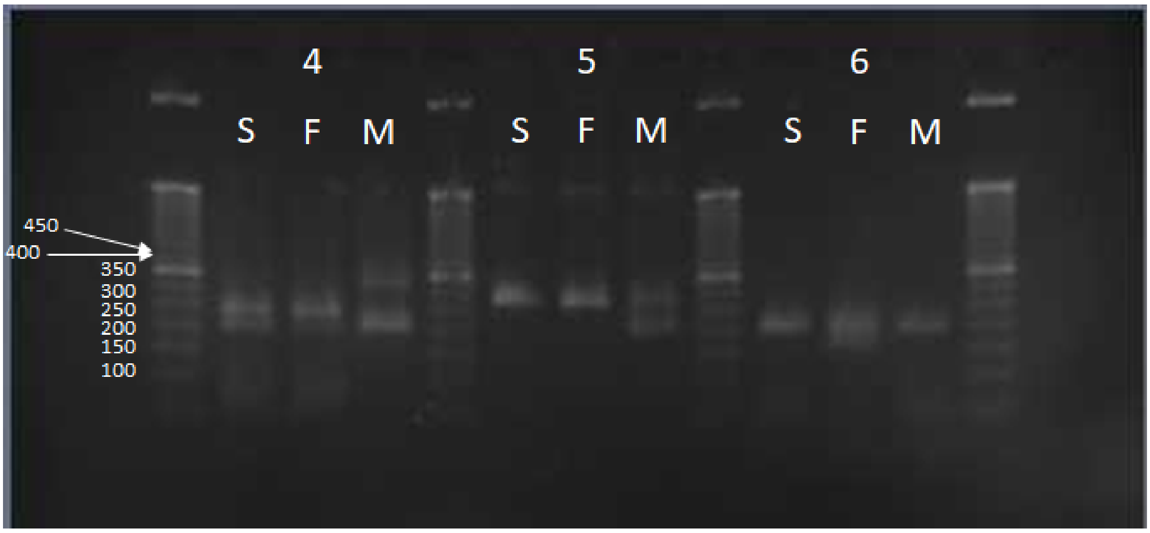


**Figure S14:** In locus 4, the son shares a larger band with the father (231bp) and a smaller band with both the mother and the father (193bp). The mother seems has a larger band at 307bp. In locus 5, all three individuals share a band at 266bp. The mother has an extra band which is smaller at 196bp. In locus 6, all individuals share a band at 195bp. The father has a smaller band at 161. All three loci confirm the VNTRSeek predictions.

| Locus | Pattern Size | TR | VNTRseek | | | Expected bands | | | Validated | | |
|---|---|---|---|---|---|---|---|---|---|---|---|
| | | | son | father | mother | son | father | mother | son | father | mother |
| 7 | 27 | 182608886 | 0, -2 | 0, -3 | -2, +1 | 208, 181 | 208, 154 | 235, 181 | yes | yes | yes |
| 8 | 48 | 182620950 | 0, -1 | 0, -1 | 0, -1 | 243, 195 | 243, 195 | 243, 195 | yes | yes | yes |
| 9 | 34 | 182982510 | 0, -1 | 0 | 0, -1 | 301, 267 | 301 | 301, 267 | yes | yes | yes |

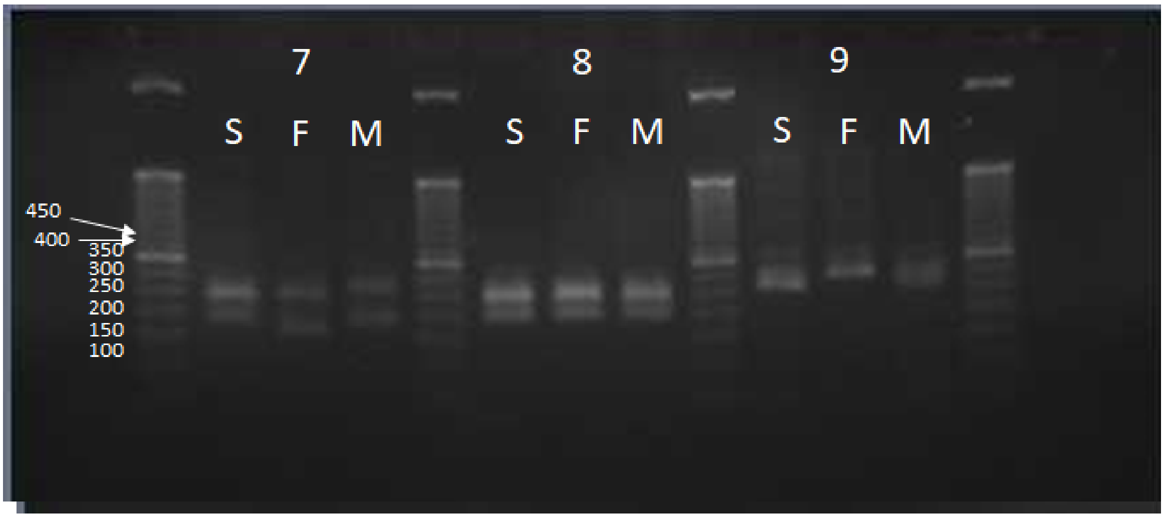


**Figure S15:** In locus 7, the son and father share a band at 208 bp and the son and mother share a band at 181 bp. The father has another smaller band at 154bp and the mother has a band slightly higher at 235 bp. In locus 8, all individuals share two bands at 195 bp and 243 bp. In locus 9 all individuals share a band at 301 bp. The son and mother share another smaller band at 267 bp. All three loci confirm the VNTRSeek predictions.

| Locus | Pattern Size | TR | VNTRseek | | | Expected bands | | | Validated | | |
|---|---|---|---|---|---|---|---|---|---|---|---|
| | | | son | father | mother | son | father | mother | son | father | mother |
| 10* | 38 | 183046759 | 0 | 0, -1 | -1 | 222 | 222, 184 | 184 | 0, **+1** | 0, -1 | -1, **+1** |
| 11 | 15 | 183081195 | +2, -1 | +2, +1 | +2, -1 | 182, 122 | 182, 167 | 182, 122 | yes | yes* | yes |

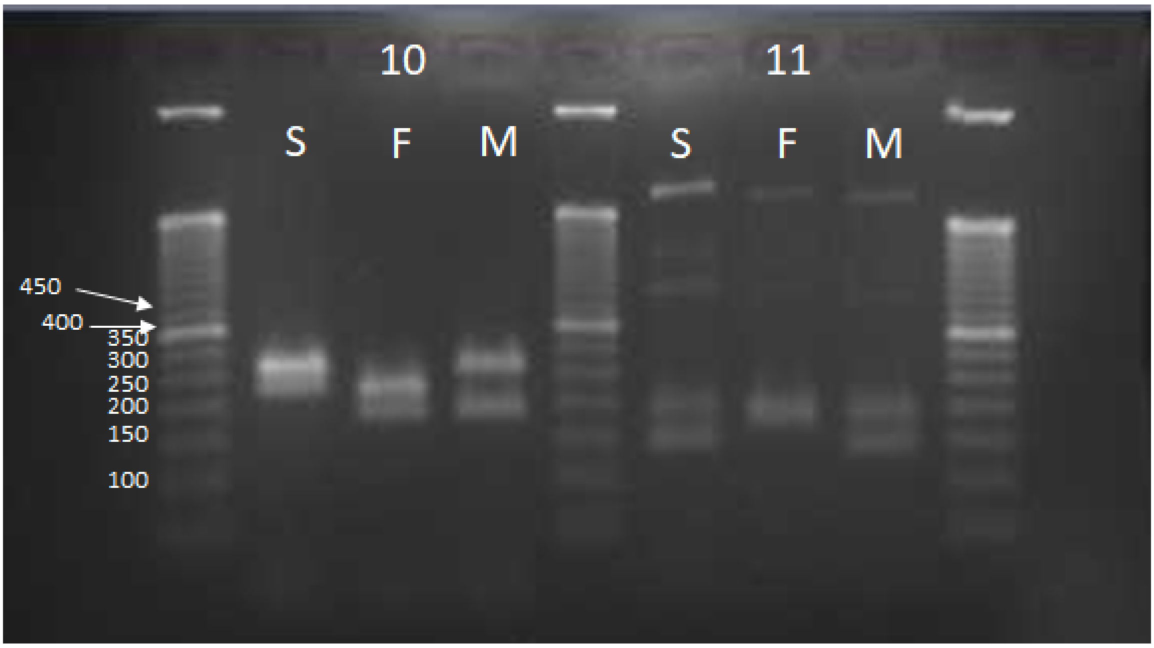


**Figure S16:** In locus 10 the son and father share a band at 222bp. The father and mother share a band at 184bp. The son and mother share a larger band which appears to be the +1 allele (260 bp), which was detectable, but not predicted in either sample by VNTRseek. The alleles predicted by VNTRSeek were validated and an extra band that could have been predicted was missed in two samples. In locus 11, all individuals share the larger band at 182bp. The son and mother have a smaller band at 122bp. If the father has two alleles as predicted by VNTRseek, then they are very close together (15 nucleotides different in length). There is a suggestion in the gel that there are two close bands. In any event, all but one of the alleles was validated.

| Locus | Pattern Size | TR | VNTRseek | | | Expected bands | | | Validated | | |
|---|---|---|---|---|---|---|---|---|---|---|---|
| | | | son | father | mother | son | father | mother | son | father | mother |
| 12 | 17 | 183117043 | -5, -3 | -5, -3 | 0, -3 | 180, 214 | 180, 214 | 265, 214 | yes | yes | yes |
| 13 | 15 | 183169331 | -2 | -2, 0 | -2 | 209 | 209, 239 | 209 | yes | yes | yes |

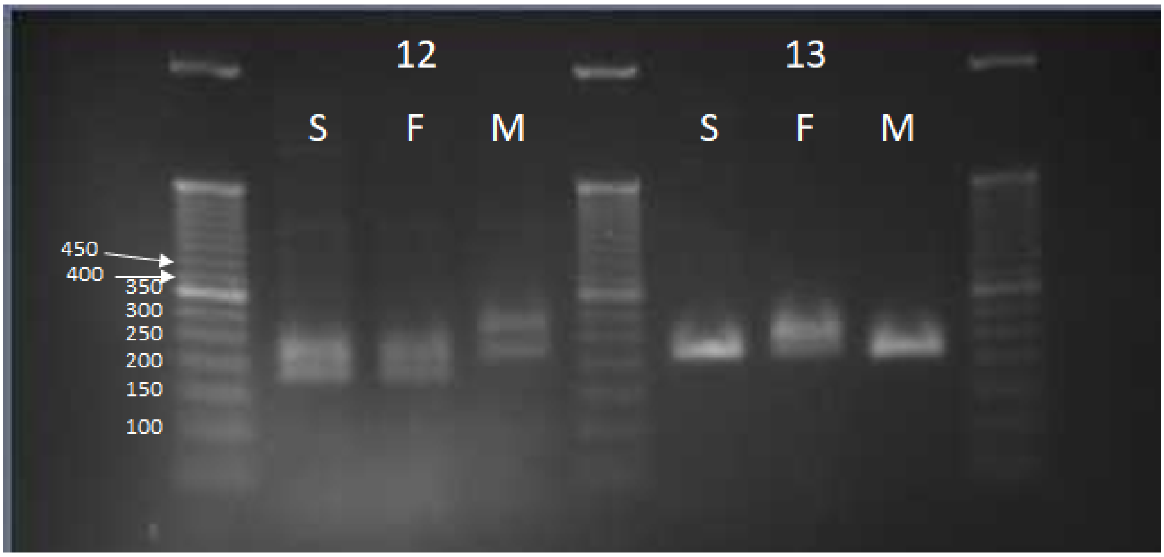


**Figure S17:** In locus 12, all individuals share a band at 214 bp. The son and father share a smaller band at 180bp. The mother has a larger band at 265 bp. In locus 13, all individuals share a band at 209 bp. The father has a larger band at 239bp. Both loci confirm the VNTRSeek predictions.

#### S6.2 Comparing to adVNTR

| Gene | TRid | Pat. | CN | Array | Δbp | Genotype | VNTRseek Detected |
|---|---|---|---|---|---|---|---|
| LCE4A | 182223827 | 24 | 2.21 | 53 | +24/+24 | 1/1 | 0/0 |
| DRD4 | 182318145 | 48 | 4.29 | 206 | 0/-48 | 0/-1 | |
| IL1RN | 182721519 | 86 | 4.00 | 344 | 0/-172 | 0/-2 | |
| C14orf180 | 182495840 | 9 | 3.11 | 28 | 0/0 | 0/0 | 0/0 |
| CSTB | 182814480 | 12 | 3.08 | 37 | 0/-12 | 0/-1 | 0/-1 |
| PAOX | 182317476 | 35 | 4.00 | 140 | -35/-35 | -1/-1 | -1/-1 |
| SSTR1 | 182469843 | 24 | 2.42 | 58 | 0/+24 | 0/1 | 0/1 |
| EIF3G | 182649420 | 21 | 2.29 | 48 | 0/-21 | 0/-1 | 0/0 |
| JAKMIP3 | 182316063 | 39 | 3.13 | 122 | 0/-78 | 0/-2 | 0/0 |
| SRSF8 | 182354360 | 21 | 2.10 | 44 | 0/-21 | 0/-1 | 0/0 |
| MAOA | 183311386 | 30 | 3.50 | 105 | +30/+30 | 1/1 | 1/1 |
| GP1BA | 182574350 | 39 | 3.92 | 153 | 0/-40 | 0/-1 | -1/-1 |
| BRWD1 | 182812082 | 22 | 2.14 | 45 | 0/-22 | 0/-1 | |
| CLCA4 | 184859017 | 6 | 5.67 | 34 | -6/-6 | -1/-1 | |
| SLC6A4 | 182584594 | 63 | 2.27 | 358 | -66/-66 | | |
| STK39 | 182741803 | 6 | 4.67 | 28 | | | |
| UBXN11 | | | | | -18/-18 | | |

| Gene | TRid | Comment |
|---|---|---|
| LCE4A | 182223827 | Error in HG001 (0/0). Not found in NA12878 150 bp and 250 bp. |
| DRD4 | 182318145 | Not found in NA12878 250 bp. Not detectable in HG001 and NA12878 150 bp. (reads too short) |
| IL1RN | 182721519 | Detectable allele (-2) Not found in NA12878 250 bp but found in the parents. Not detectable in HG001 and NA12878 150 bp. |
| C14orf180 | 182495840 | Matched in all. |
| CSTB | 182814480 | Matched in all. |
| PAOX | 182317476 | Matched in all. |
| SSTR1 | 182469843 | Matched in all. |
| EIF3G | 182649420 | Detectable allele (0) matched in all. Other allele too short for TRF detection. |
| JAKMIP3 | 182316063 | Detectable allele (0) matched in HG001 and NA12878 250 bp. Not found in NA12878 150 bp. Other allele too short for TRF detection. |
| SRSF8 | 182354360 | Detectable allele (0) matched in all. Other allele too short for TRF detection. |
| MAOA | 183311386 | Matched in NA12878 250 bp. Not detectable in HG001 and NA12878 150 bp. |
| GP1BA | 182574350 | Detectable allele (-1) matched in NA12878 250 bp, not found in NA12878 150 bp or HG001. Detectable allele (0) not found n NA12878 250 bp. The (0) allele was not detectable in HG001 and NA12878 150 bp. |
| BRWD1 | 182812082 | Not in VNTRseek reference set. |
| CLCA4 | 184859017 | Not in VNTRseek reference set. |
| SLC6A4 | 182584594 | Not in VNTRseek reference set. |
| STK39 | 182741803 | Not in VNTRseek reference set, not validated in adVNTR paper. |
| UBXN11 | | Matching TR in HG38 could not be determined. |

**Table S7: Comparison of VNTRseek allele calls to validations from the adVNTR paper.** VNTRseek validation results for three datasets from the same genome, HG001 (148 bp) from GIAB, NA12878 (150 bp) from NYGC, and NA12878 (250 bp) from 1000 Genomes Phase 3 HC for VNTR loci as reported in the adVNTR paper (4). Results were originally reported for hg19. Coordinates were converted to hg38 and TRDB TR ids (5). In one case, a matching TR could not be determined. In total, 11 out of 16 detectable alleles were correctly predicted, four were not found in the NA12878 250 bp sample, and one was incorrectly predicted in the HG001 sample. Note that for the IL1RN gene, the missed allele in the NA12878 250 bp sample was correctly predicted in the parent samples NA12891 and NA12892.

##### S6.3 Validations using long precise reads from PacBio

| Locus Category | All | All VNTR | All Hom | Hom Ref. | Hom VNTR | All Het | Het 0/1 | Het 1/2 |
|---|---|---|---|---|---|---|---|---|
| VNTRseek genotyped | 170,481 | 2597 | 169,175 | 1167,884 | 1,291 | 1,306 | 1,139 | 167 |
| With 0 alleles validated | 4,121 | 226 | 4,074 | 3,895 | 179 | 47 | 38 | 9 |
| With 1 allele validated | 165,303 | 1,314 | 164,996 | 165,101 | 1,112 | 202 | 173 | 29 |
| With 2 alleles validated | 1,057 | 1,057 | – | – | – | 1,057 | 928 | 129 |
| Alleles PPV | 97.46% | 87.83% | 97.59% | 97.68% | 86.13% | 88.67% | 89.07% | 85.93% |

**Table S8: VNTRseek validation in PacBio long reads.** PacBio Circular Consensus Sequencing reads from the HG002 genome (6), with an average length of 13.5 Kbp and an estimated 99.8% sequence accuracy, were computationally tested to determine if they confirmed VNTRseek predicted alleles from the GIAB Illumina reads. If a VNTRseek predicted allele copy number from the GIAB data was matched in at least one PacBio read previously mapped to the VNTR locus, the allele was considered confirmed. Matching copy numbers were defined as differing by at most 0.25 copy. Counts are shown for all loci and separately for homozygous and heterozygous loci. Het 0/1 are heterozygous VNTR loci called with one reference allele and one variant allele. Het 1/2 are heterozygous VNTR loci called with two variant alleles. Alleles PPV (positive predictive value) is the percentage of VNTRseek predicted *alleles* that were validated. Overall, more than 97% of predicted alleles were validated, and at the predicted VNTR loci, more than 87% of alleles were validated.

##### S6.4 Consistency of VNTR genotypes with Mendelian inheritance

| Trio: | AJ | HAN | CEPH | YRI |
|---|---|---|---|---|
| Heterozygous in both parents | 358 | 224 | 302 | 469 |
| Inconsistent | 2 | 2 | 2 | 7 |
| Heterozygous in all three | 262 | 178 | 215 | 349 |
| Inconsistent | 0 | 2 | 1 | 3 |
| Heterozygous in all three and different | 59 | 32 | 53 | 105 |
| Inconsistent | 0 | 2 | 1 | 3 |
| All on chrY of son | 911 | 884 | - | - |
| Inconsistent | 0 | 0 | - | - |
| All on chrX of son | 6,701 | 6,493 | - | - |
| Inconsistent | 2 | 0 | - | - |

**Table S9: Consistency with Mendelian inheritance of VNTR genotypes in trios from GIAB and the 1000 Genomes Phase 3 HC datasets.** Only loci detected in all members of a trio were considered, but with increasingly stringent criteria. Loci were overwhelmingly consistent.

#### S6.5 Consistency of VNTRseek allele predictions across platforms

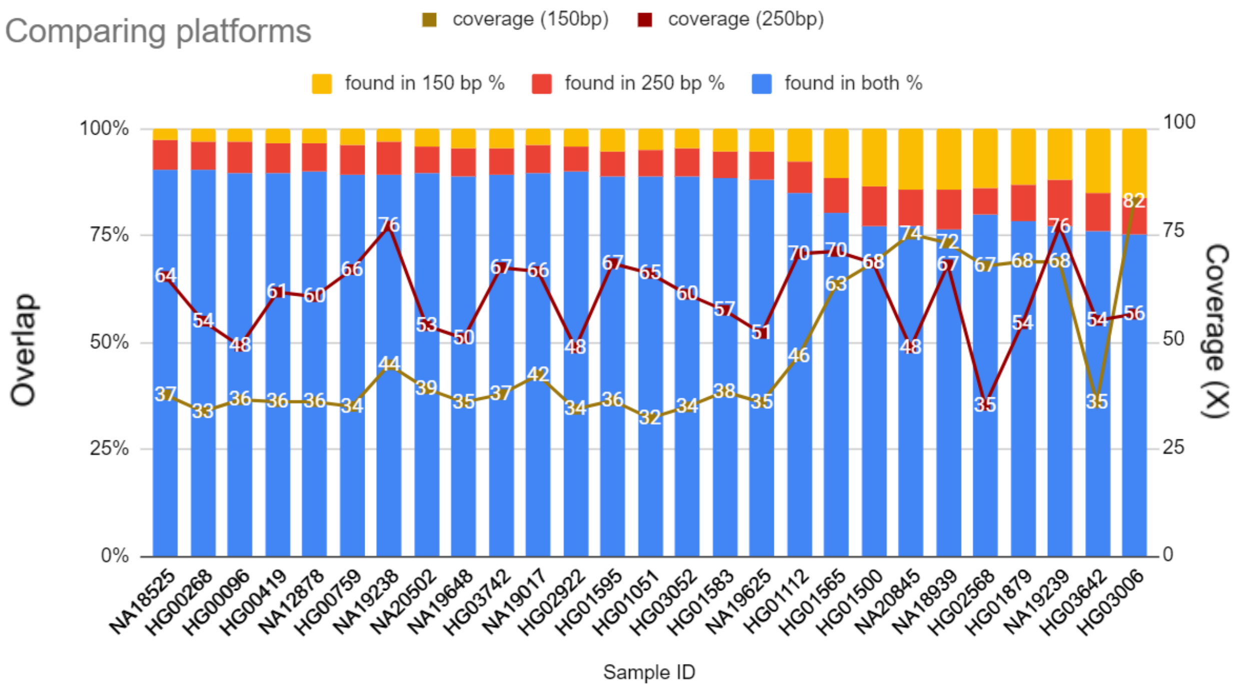


**Figure S18: Comparison of VNTRseek allele prediction across platforms.** Comparison was done on the 27 genomes in common in the 1000 Genomes Phase 3 HC and NYGC datasets. Only non-reference alleles detected in both genomes were included. Agreement of allele calls in both platforms ranged from 76% to 91%. Each bar represents one genome. Colored bar heights indicate percentage of agreement between predicted alleles (blue) or alleles found in one sample only (read and yellow). Lines indicate the read coverage in the two platforms in each sample. Generally, as the read coverage of the 150 bp sample increased above that of the 250 bp sample, more alleles were found in the former that were not detected in the latter, which was expected due to higher statistical power. Conversely, when the coverage was higher in the 250 bp sample, that platform found more alleles not detected in the 150 bp sample.

### S7 Analysis of VNTRs across populations

#### S7.1 Prediction of Ancestry from common VNTRs

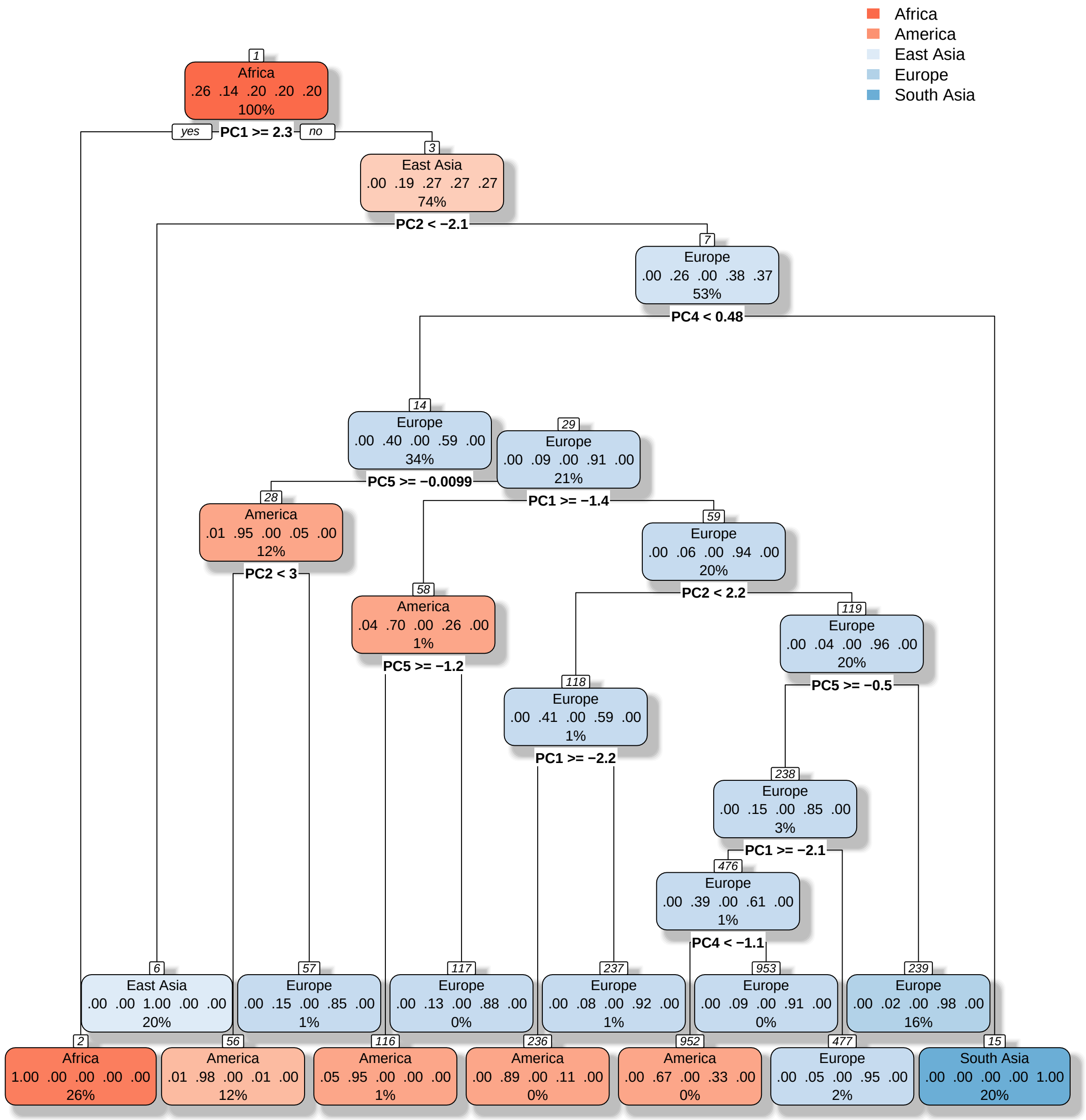


**Figure S19: Decision tree for prediction of superpopulation ancestry from common VNTRs.** Each box is a node in the decision tree. Color and superpopulation name indicate majority label in the training data entering the node. Five decimal values are the fractions of population labels in training data entering the node with values in order as Africa, America, East Asia, Europe, and South Asia. Percentage shown is percent of training data entering the node. Equation with PC number indicates Principal Component test value to exit the node down "yes" (left) branch.

| N=751 | African | American | East Asian | European | South Asian | Precision |
|---|---|---|---|---|---|---|
| African | 205 | 0 | 0 | 0 | 0 | 100% |
| American | 4 | 91 | 0 | 3 | 0 | 93% |
| East Asian | 0 | 1 | 146 | 0 | 0 | 99% |
| European | 0 | 6 | 0 | 141 | 0 | 96% |
| South Asian | 0 | 0 | 0 | 0 | 154 | 100% |
| Recall | 98% | 93% | 100% | 98% | 100% | accuracy=98% |

**Table S10: Confusion matrix of decision tree results on the test data to predict ancestry.** Common VNTR predictions on 2,504 unrelated genomes from NYGC were used to train a model to predict ancestry. Principal Component Analysis was performed to reduce dimensionality. The first 10 Principal Components were used to train 70% of the data (train). The model was tested on the remaining test data (30%). The confusion matrix on the test data is presented here. The total number of genomes in the test was 751. Columns indicate the *true* label, rows the *predicted* label. The last column shows the *precision*, and the last row shows the *recall*. Populations of African, East Asian, and South Asian ancestry were the easiest to predict. People with American ancestry, on the other hand, had more admixed genomes and fewer samples (as described by the data source) making them more difficult to predict. Overall accuracy was 98%.

##### S7.2 Population-specific VNTR alleles

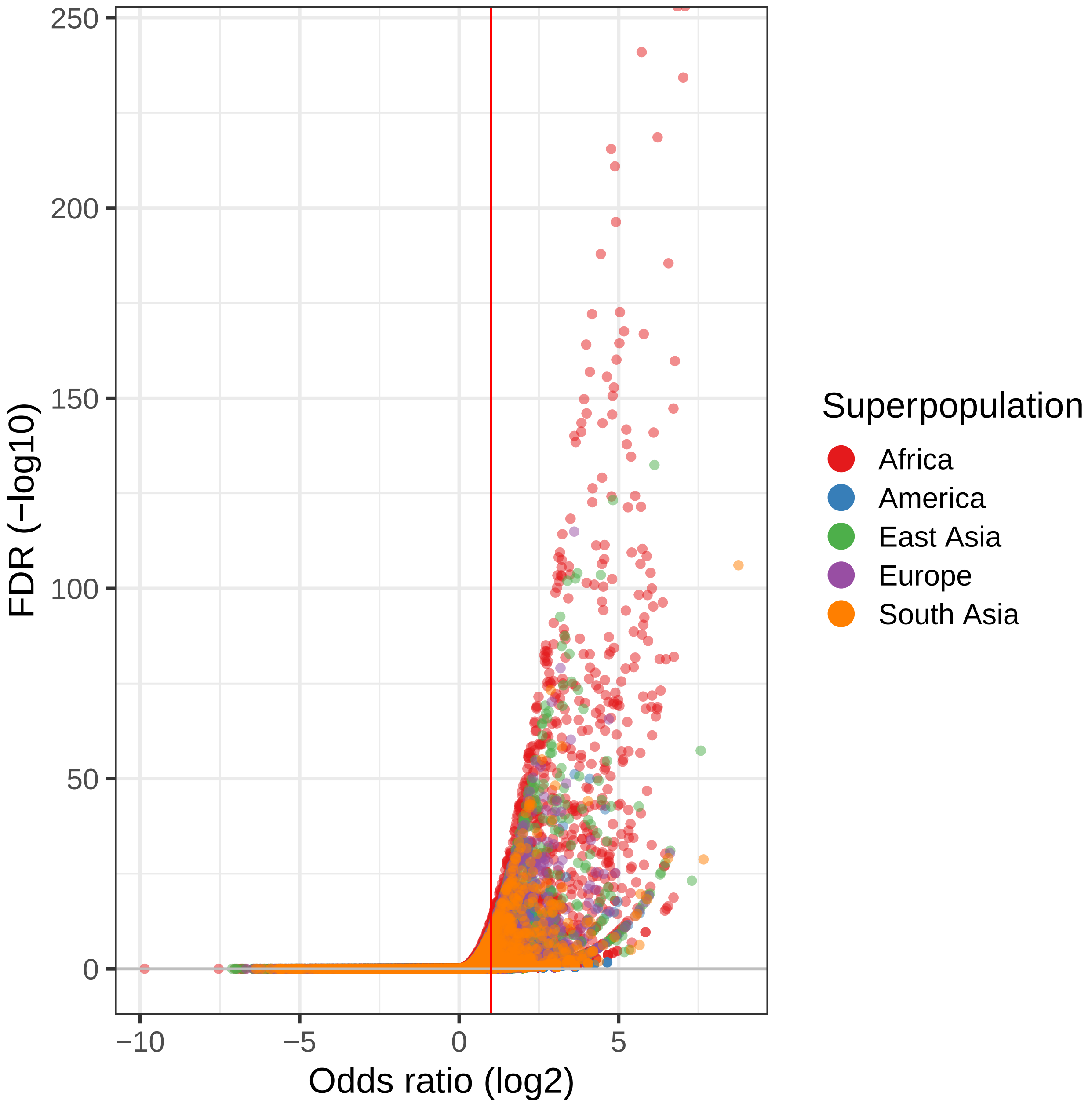


**Figure S20: Volcano plot for alleles at population-specific VNTR loci.** One-way Fisher's Exact Test was used to find common VNTR *alleles* over-represented in each population versus the others. Each dot in the volcano plot represents one allele tested for overrepresentation in one population (depicted by color). The p-values were adjusted using FDR. Alleles with FDR<5% (above the horizontal gray line nearly coinciding with zero) and with odds ratio > 2 (right of the red line) were selected.

| TRid | Allele | Specific to | Odds Ratio (log2) | FDR | AFR N=661 | AMR N=347 | EAS N=504 | EUR N=503 | SAS N=489 |
|---|---|---|---|---|---|---|---|---|---|
| 182229555 | (+1) | African | 6.73 | 1E-82 | 148 | 2 | 1 | 1 | 0 |
| 182232436 | (0) | African | 7.03 | 5E-235 | 381 | 15 | 0 | 2 | 1 |
| 182247194 | (+2) | East Asian | 7.29 | 7E-24 | 0 | 0 | 36 | 0 | 0 |
| 182272465 | (-2) | African | 4.85 | 4E-34 | 73 | 7 | 0 | 0 | 0 |
| 182311248 | (+1) | South Asian | 8.76 | 8E-107 | 0 | 0 | 0 | 1 | 147 |
| 182423923 | (-1) | East Asian | 6.12 | 4E-133 | 4 | 3 | 208 | 5 | 7 |
| 182454990 | (+1) | South Asian | 7.63 | 1E-28 | 0 | 0 | 0 | 0 | 43 |

**Table S11: Example of population specific alleles.** Seven significant alleles were chosen to draw a virtual gel (main text). The details of the test on those seven VNTR alleles and the raw counts in each population for the specified allele are given here. $N$ is the total number of genomes in that population. Allele column indicates copy number change relative to the reference allele.

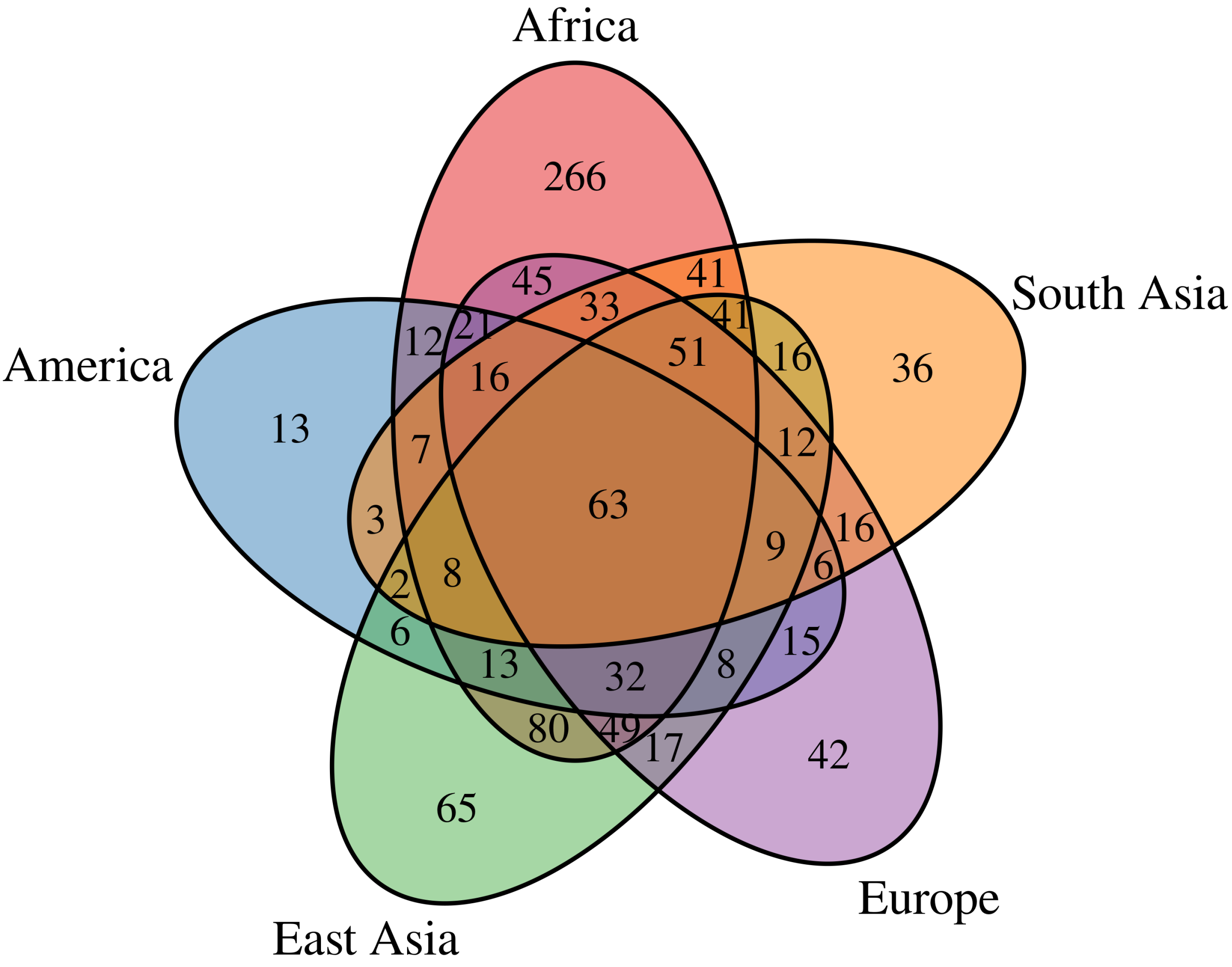


**Figure S21: Venn diagram of population-specific VNTR loci.** Africans have the highest number of loci with population-specific alleles, followed by East Asians. Americans had the least which could be because of the lower number of samples (statistical power) and/or more admixed genomes (7). Interestingly 63 loci had an allele that is over-represented in each population.

#### S7.3 Gene set enrichment of population-specific VNTRs

| Gene Set Name | KEGG id | Gene set size (K) | No. of genes in overlap ($k$) | $k$/K | p-value | FDR q-value |
|---|---|---|---|---|---|---|
| Endocytosis | hsa04144 | 181 | 11 | 6% | 2.51 e-5 | 4.67 e-3 |
| Fatty acid metabolism | hsa01212 | 42 | 5 | 12% | 1.98 e-4 | 1.84 e-2 |
| Arrhythmogenic right ventricular cardiomyopathy (ARVC) | hsa05412 | 74 | 6 | 8% | 3.87 e-4 | 2.4 e-2 |
| Calcium signaling pathway | hsa04020 | 178 | 9 | 5% | 5.29 e-4 | 2.46 e-2 |
| Vascular smooth muscle contraction | hsa04270 | 115 | 7 | 6% | 7.39 e-4 | 2.75 e-2 |

**Table S12: KEGG pathways enriched for population-specific VNTR genes.** GSEA (8) was used to find enrichment of the 560 protein coding genes overlapping with 1,096 population-specific VNTR loci.

#### S7.4 Genes enriched in three GO terms.

Below are the genes with VNTRs that were enriched for biological processes behavior, neuron development and differentiation:

**Behavior:** ADORA1, CAPN2, CDH23, CPT1A, DAB1, DACH1, DEAF1, DRD4, GRID1, ITGA8, JPH4, KCNA2, LRRK2, NPHP4, NRXN2, NTRK1, OPRD1, OTOG, P2RX2, PARK7, POU4F1, PRKCZ, RETN, SGIP1, SHANK2, TACR2, TAFA2.

**Neuron development:** ABI1, ACAP3, AGBL4, AGRN, ANOS1, BRSK2, CAMK1D, CAMSAP2, CDH23, CNTN1, DAB1, DOCK10, DPYSL4, DSCAML1, EHD1, ENAH, EPHA10, FARP1, HERC1, IFT88, JAM3, KNDC1, LRRC4C, LRRK2, MOB2, MYO16, NANOS1, NEUROD4, NPHP4, NRL, NTM, NTNG1, NTRK1, OPCML, PARD3, POU4F1, PPFIA2, PRKCQ, PRKCZ, PRKG1, RET, RETREG3, SHANK2, SRRM4, SZT2, TENM3, TENM4, TWF1, UBE4B, USH1C, ZMIZ1.

**Neuron differentiation:** ABI1, ACAP3, AGBL4, AGRN, ANOS1, BRINP2, BRSK2, CAMK1D, CAMSAP2, CASZ1, CBLN1, CDH23, CHD5, CNTN1, DAB1, DISP3, DOCK10, DPYSL4, DSCAML1, EHD1, ENAH, EPHA10, FARP1, HERC1, IFT88, JAM3, KNDC1, LRP6, LRRC4C, LRRK2, MOB2, MYO16, NANOS1, NEUROD4, NPHP4, NRL, NTM, NTNG1, NTRK1, OPCML, PARD3, PAX7, POU4F1, PPFIA2, PRKCQ, PRKCZ, PRKG1, RET, RETREG3, SHANK2, SRRM4, SZT2, TENM3, TENM4, TWF1, UBE4B, USH1C, WNT4, ZMIZ1.

#### S8 Correlation of VNTR genotypes with gene expression

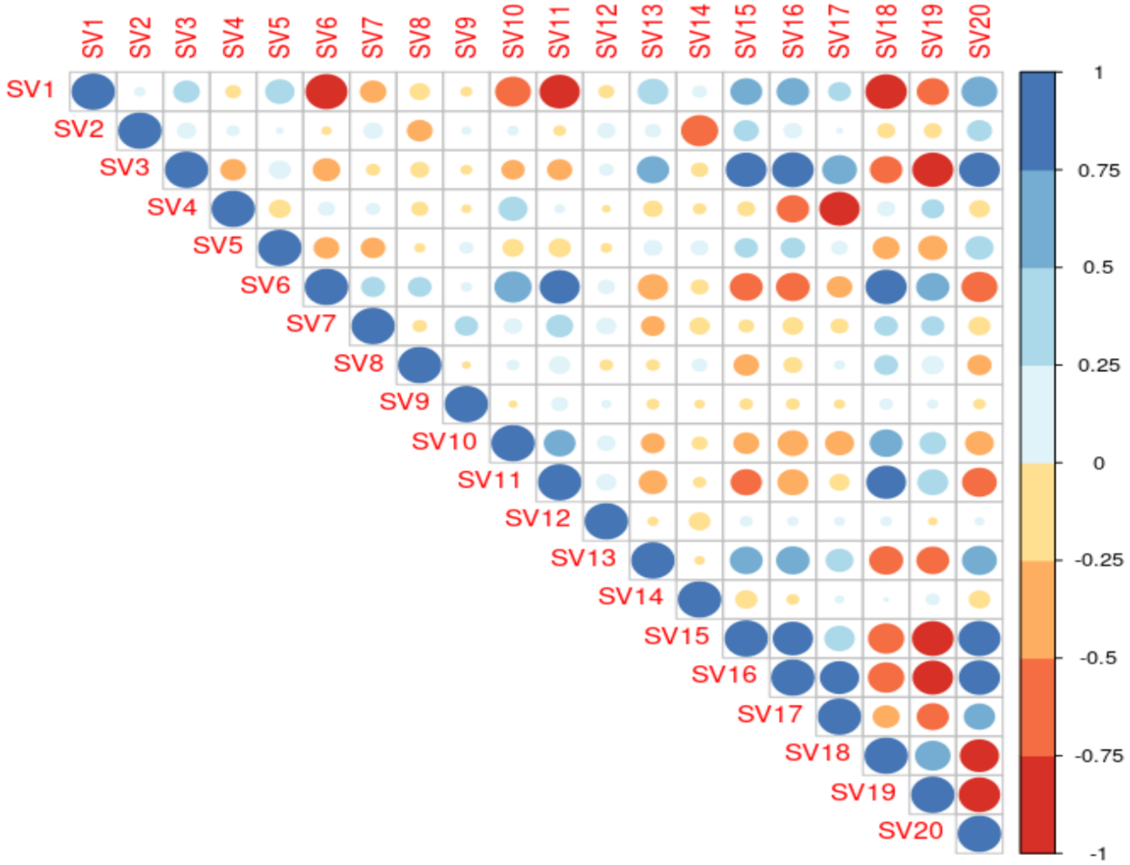


**Figure S22: Detecting hidden covariates with SVA** To detect unknown confounders, we applied Iteratively Adjusted Surrogate Variable Analysis (iasva) (9) on the log2 normalized TPM values of mRNA expression from the Geuvadis consortium (Accession: E-GEUV-1). We observed the first covariates were independent and the covariates 6-10 were >85% correlated to other covariates. Thus, we chose five hidden factors to include in our model.

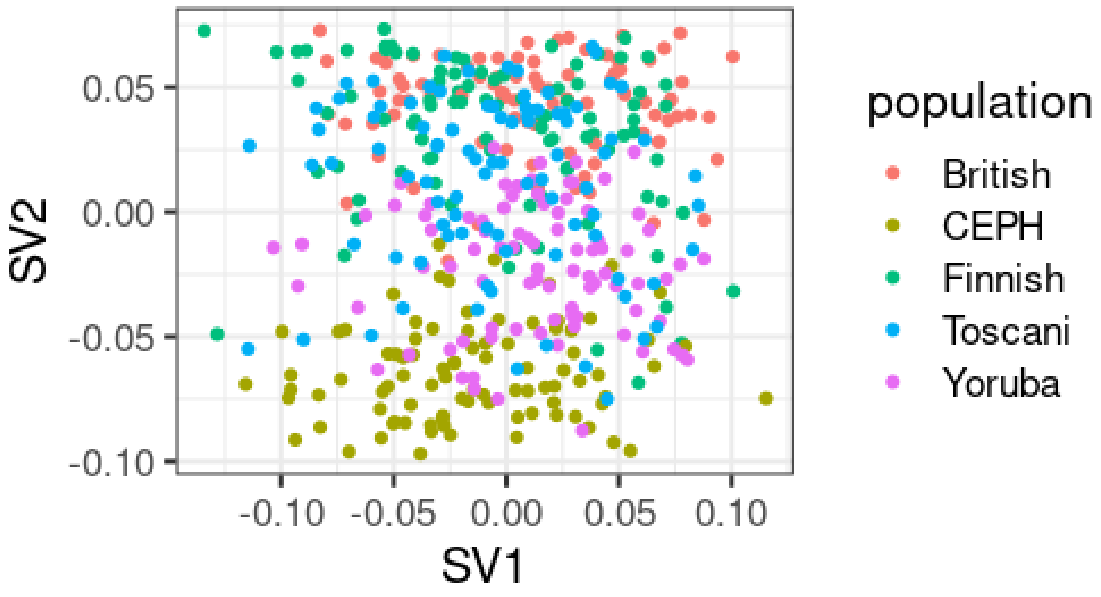


**Figure S23:** First and second hidden factors.

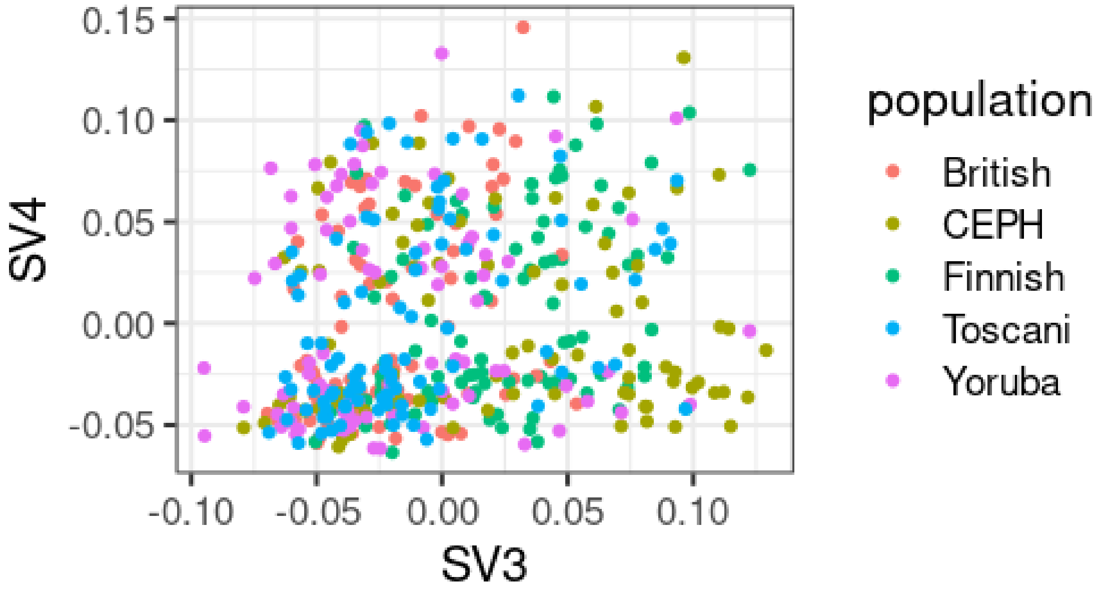


**Figure S24:** Third and fourth hidden factors.

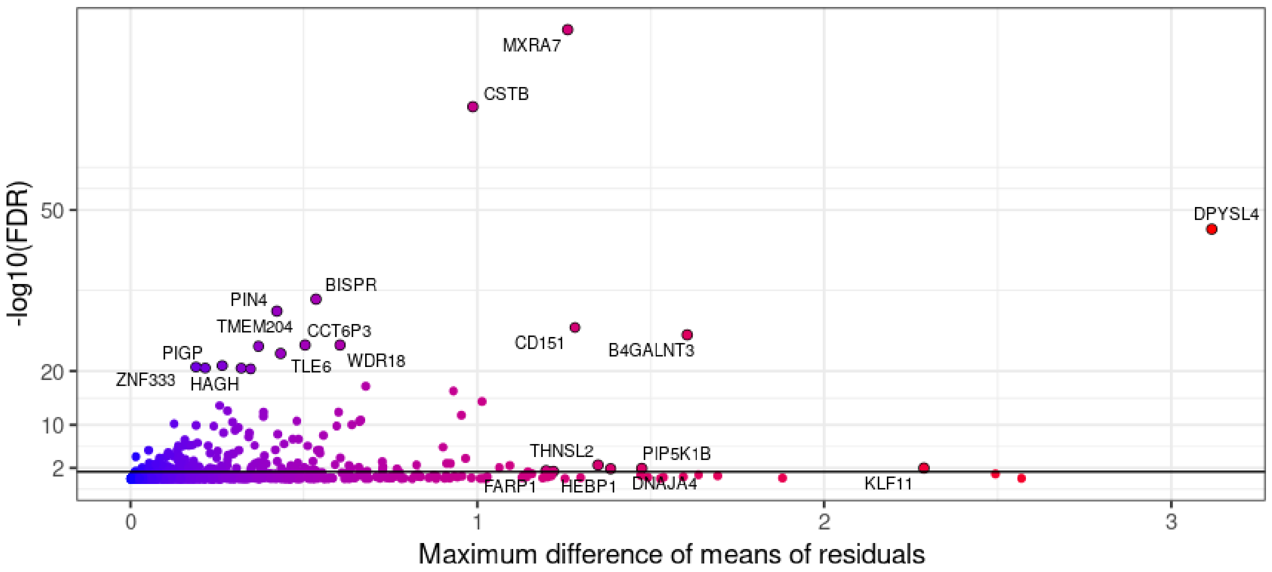


**Figure S25: Correlation of gene expression with VNTR genotype.** In 193 gene-VNTR pairs (dots above the black line), a significant difference in gene expression was correlated with VNTR genotype. These pairs consist of 187 genes and 188 VNTR loci. The y-axis is $-\log_{10}$ of the FDR value and the dashed black line denotes FDR=0.05. The X-axis is the maximum difference between the mean of residuals for the different genotypes. Top genes by significance and/or mean difference are labeled.

#### S8.1 Examples of eQTL VNTRs

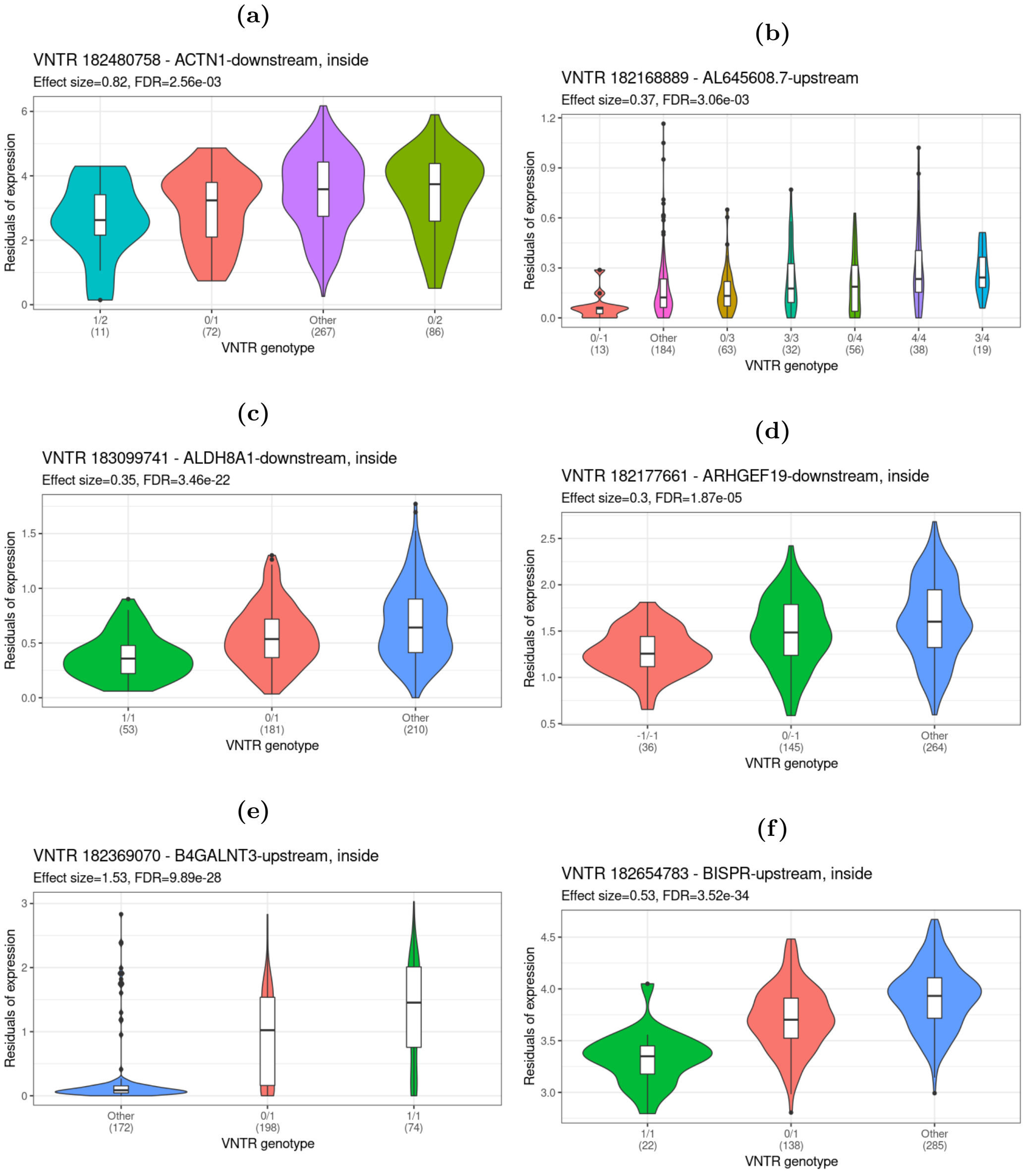


**Figure S26: Gene expression differences and VNTR genotype.** Shown are violin plots of residuals for six genes which displayed significant differential expression when samples were partitioned by VNTR allele genotype. Genotype is indicated in labels on the x-axis and numbers refer to copies gained or lost relative to the reference allele. “Other” indicates a partition with undetected alleles presumed outside the range of VNTRseek detection (see main text). Number of samples in each partition is shown in parenthesis. Associated VNTR loci either overlapped the genes or were within 10 Kbp upstream or downstream. In these examples, the effect size for at least one genotype class was significant.

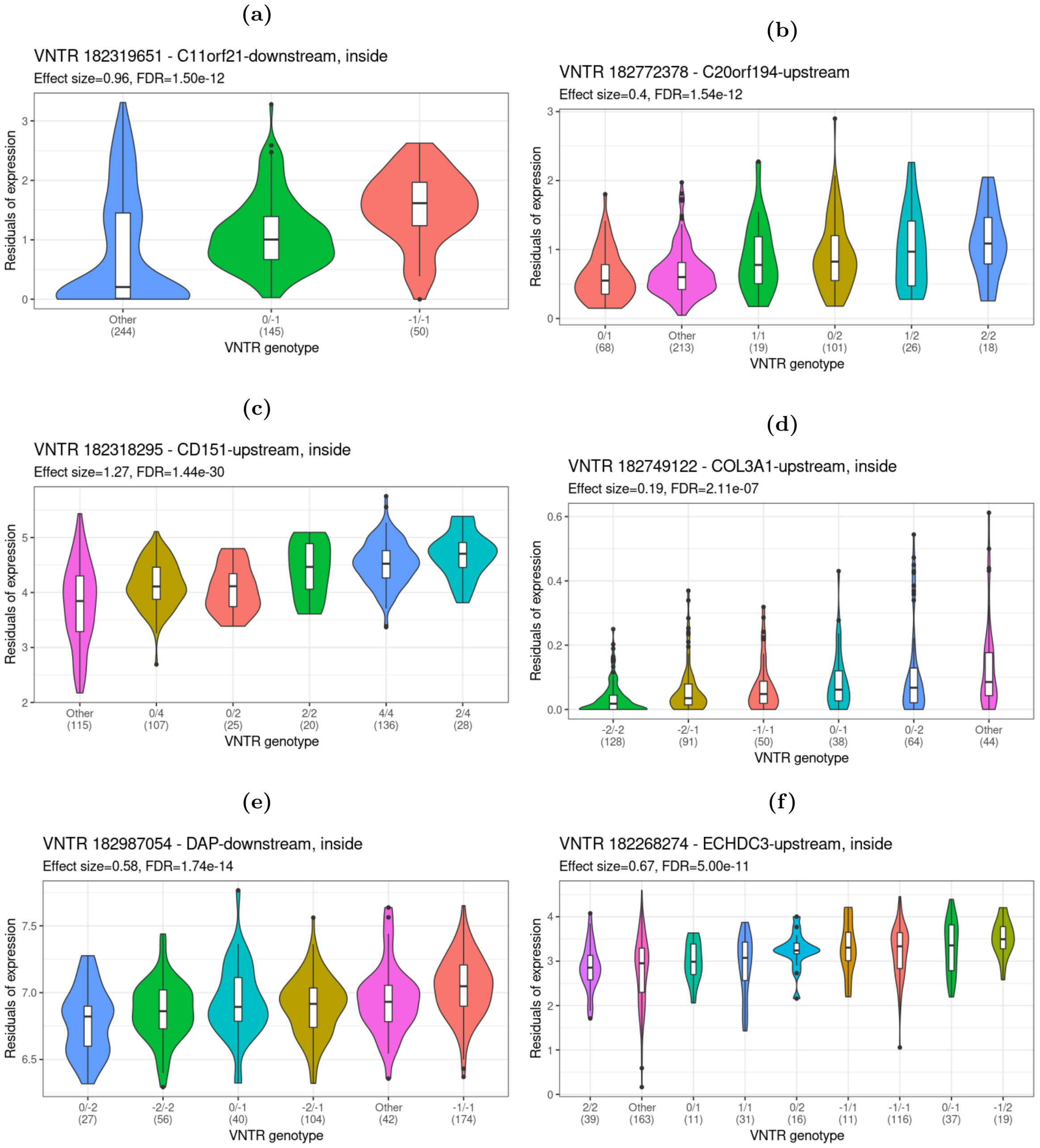


**Figure S27: Continued... Gene expression differences and VNTR genotype.** See caption for Supplementary Figure S26. Six examples are shown here.

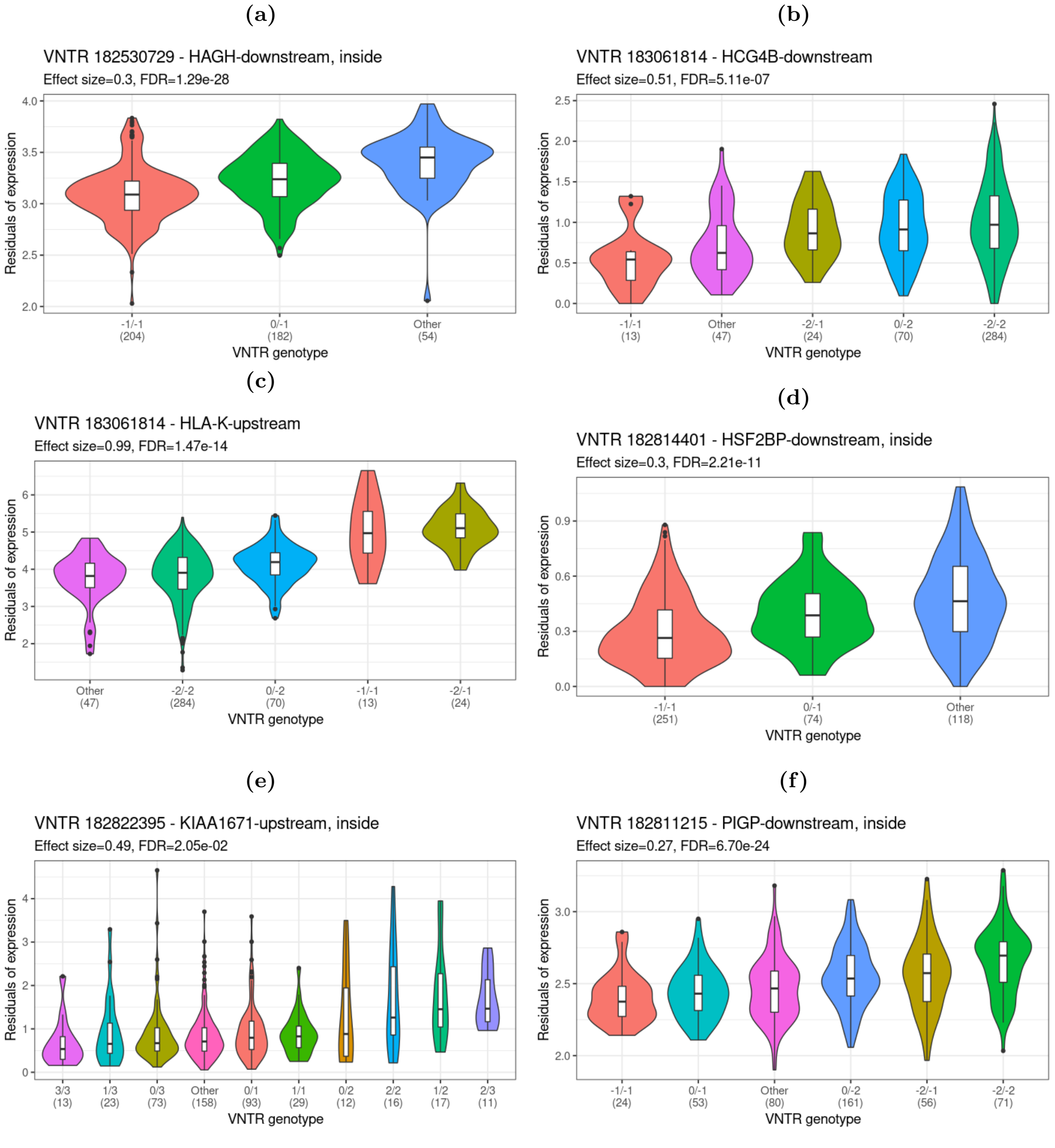


**Figure S28: Continued... Gene expression differences and VNTR genotype.** See caption for Supplementary Figure S26. Six examples are shown here.

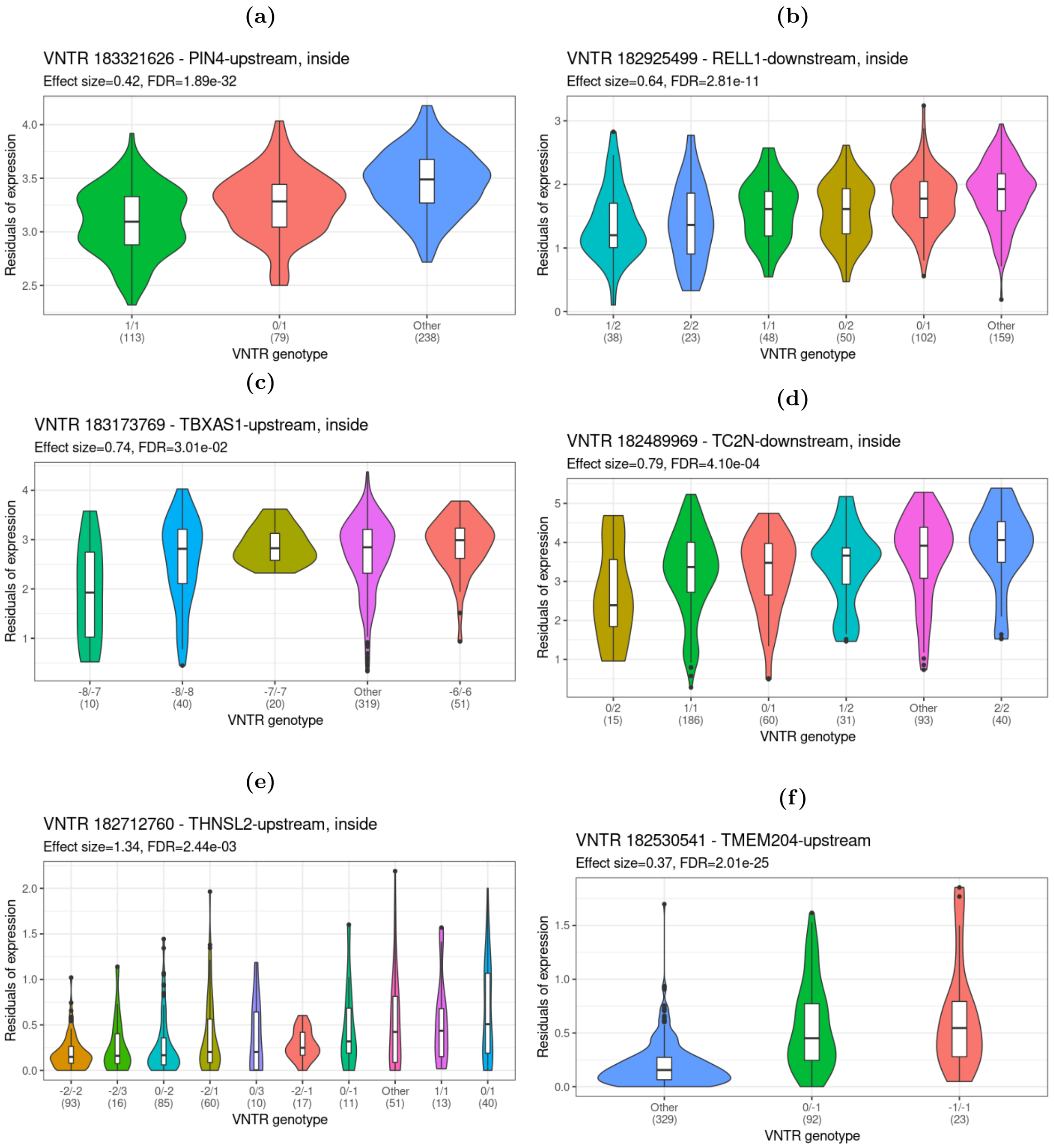


**Figure S29: Continued... Gene expression differences and VNTR genotype.** See caption for Supplementary Figure S26. Six examples are shown here.

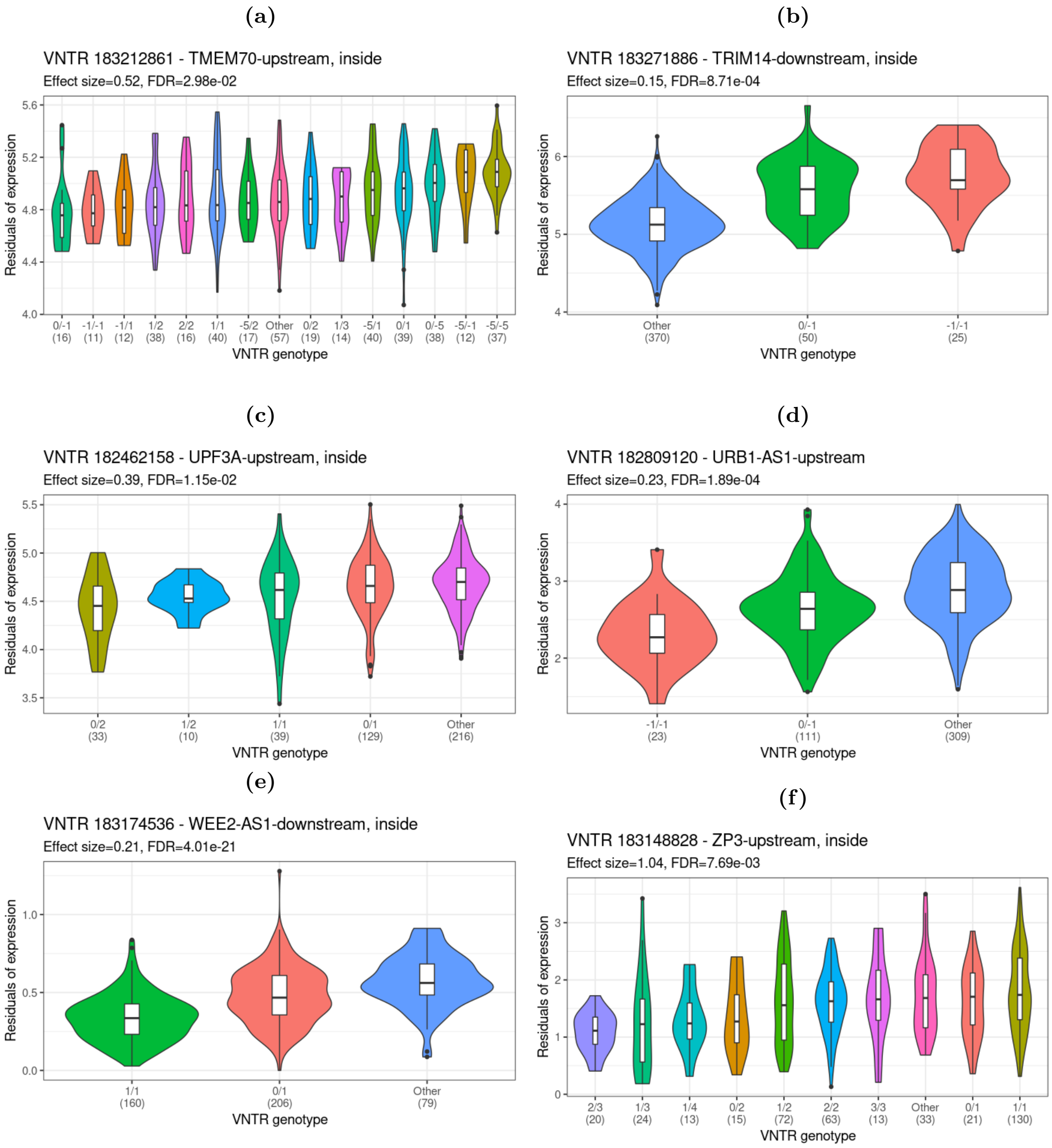


**Figure S30: Continued... Gene expression differences and VNTR genotype.** See caption for Supplementary Figure S26. Six examples are shown here.

##### S8.2 Overlap of eQTL VNTRs with histone marks and DNAse hypersensitive sites

| Mark | Source ID | Number of overlaps |
|---|---|---|
| H3K27me3 | ENCFF695TUH | 0 |
| H3K27ac | ENCFF835NLI | 33 |
| H3K36me3 | ENCFF695TUH, ENCFF829MGL | 8 |
| H3K4me1 | ENCFF710GQV | 11 |
| H3K4me2 | ENCFF772RNW | 43 |
| H3K4me3 | ENCFF566VFL, ENCFF920KUR | 48 |
| H3K79me2 | ENCFF522JVO | 20 |
| H3K9ac | ENCFF797IEH | 36 |
| H3K9me3 | ENCFF874UEV | 0 |
| H4K20me1 | ENCFF774QTB | 1 |
| DNAse | ENCFF588OCA | 33 |
| Total eQTL VNTRs that overlap marks | | 81 |

**Table S13: Number of overlaps of histone marks and DNAse hypersensitive site peaks with the eQTL VNTRs.** Data from ENCODE experiments on the GM12878 cell line were selected. Narrow peak calls in bed format on GRCh38 were downloaded. Any intersection of a peak with a VNTR locus was counted as an overlap. The ID of the file from ENCODE is given in the "Source ID" column. A total of 81 eQTL VNTRs overlapped with one or more marks.

| | Overlaps mark | No overlap | Marginal row totals |
|---|---|---|---|
| eQTL | 81 | 107 | 188 |
| Not eQTL | 195 | 584 | 779 |
| Marginal column totals | 276 | 691 | 967 |

**Table S14: Enrichment of histone marks and DNAse hypersensitive sites in eQTL VNTRs.** eQTL VNTRs were more likely to overlap with histone marks and DNAse hypersensitive sites than non-eQTL VNTRs at p-value $< 0.00001$.

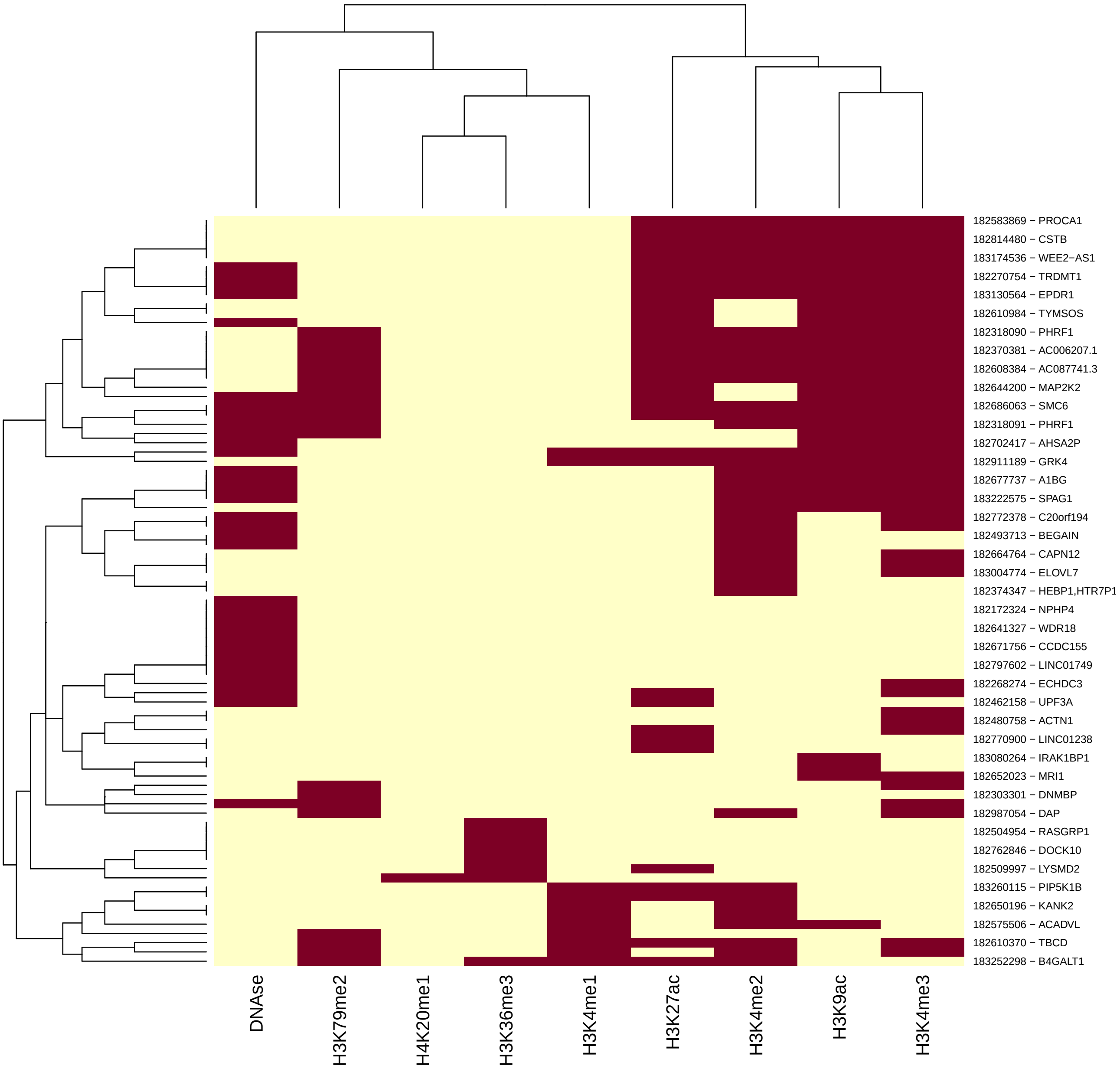


**Figure S31: Clustering of eQTL VNTRs with peaks for histone marks and DNAse hypersensitivity sites.** For source of data see Table S13. A total of 81 out of 188 eQTL VNTRs overlapped with histone marks and DNAse hypersensitivity sites. Hierarchical clustering was performed to illustrate the overlaps. Marks are in columns, VNTR-gene pairs are in rows.

#### S8.3 eQTL loci with population-biased alleles

**Table S15: Intersection of eQTL VNTRs with population-specific VNTRs.** Shown are 49 eQTL VNTR – gene pairs (48 genes and 47 VNTRs) where the VNTRs were also found to be population specific. Maximum mean difference refers to the maximum mean difference of the residuals for all pairs of genotypes.

| Gene | TR id | Maximum mean difference | FDR |
|---|---|---:|---|
| AC004865.2 | 182187335 | 0.66 | 8E-03 |
| AC006207.1 | 182370381 | 0.36 | 7E-07 |
| ADGRD1 | 182422326 | 0.18 | 2E-02 |
| AL645608.6 | 182168797 | 0.10 | 5E-02 |
| AL645608.7 | 182168889 | 0.37 | 3E-03 |
| ANO9 | 182317968 | 0.68 | 6E-18 |
| AP003071.5 | 182344338 | 0.16 | 4E-02 |
| AP2A2 | 182318359 | 0.13 | 6E-11 |
| ARHGEF19 | 182177661 | 0.29 | 2E-05 |
| B4GALNT3 | 182369070 | 1.60 | 2E-27 |
| C11orf21 | 182319651 | 0.95 | 2E-12 |
| CD151 | 182318295 | 1.28 | 7E-29 |
| CFAP54 | 182405813 | 0.09 | 5E-02 |
| CLIP1 | 182417629 | 0.09 | 3E-02 |
| DHRS3 | 182176034 | 0.23 | 5E-02 |
| DIP2B | 182387680 | 0.10 | 4E-02 |
| DNMBP | 182303301 | 0.31 | 3E-10 |
| DPYSL4 | 182316137 | 3.12 | 4E-47 |
| ECHDC3 | 182268274 | 0.66 | 2E-11 |
| EPS8L2 | 182318207 | 0.53 | 2E-02 |
| FARP1 | 182454752 | 1.20 | 3E-02 |
| FUT11 | 182293260 | 0.16 | 1E-02 |
| HEBP1 | 182374347 | 1.22 | 4E-02 |
| HTR7P1 | 182374347 | 0.06 | 3E-03 |
| LHPP | 182312799 | 0.20 | 6E-03 |
| MEAF6 | 182188225 | 0.12 | 7E-05 |
| MGMT | 182314781 | 0.20 | 5E-02 |
| MRPL51 | 182371420 | 0.07 | 3E-02 |
| NEBL | 182272465 | 0.35 | 6E-03 |
| NPHP4 | 182172261 | 0.28 | 3E-04 |
| NPHP4 | 182172324 | 0.31 | 6E-06 |
| NTRK1 | 182225943 | 0.11 | 3E-02 |
| NVL | 182251099 | 0.18 | 2E-02 |
| PARD3 | 182278429 | 0.49 | 4E-08 |
| PDHX | 182331520 | 0.14 | 1E-04 |
| PGGHG | 182317816 | 0.34 | 3E-09 |
| PHRF1 | 182318090 | 0.42 | 6E-04 |
| PHRF1 | 182318091 | 0.06 | 1E-02 |
| PTPRVP | 182242989 | 0.27 | 1E-02 |
| RET | 182281227 | 0.02 | 4E-02 |
| RGS5 | 182228303 | 0.09 | 2E-02 |
| S100A10 | 182223480 | 0.47 | 4E-08 |
| SELENON | 182182434 | 0.73 | 2E-02 |
| TCP11L1 | 182330802 | 0.26 | 3E-14 |
| TMEM52 | 182169710 | 0.19 | 7E-03 |
| TNS2 | 182388680 | 0.03 | 9E-03 |

*Continued on next page*

Table S15 – *Continued from previous page*

| Gene | TR id | Maximum mean difference | FDR |
|---|---|---|---|
| TRDMT1 | 182270754 | 0.19 | 1E-10 |
| UPF3A | 182462158 | 0.38 | 2E-02 |
| ZNF641 | 182386450 | 0.18 | 6E-07 |

### S9 Performance of VNTRseek on simulated data

Performance of VNTRseek was tested on a simulated genome. Using all the reference set TRs, random gain or loss of one pattern copy was inserted such that:

- one-sixth of the TR loci had *heterozygous gain* of one copy (0/+1),
- one-sixth of the TR loci had *homozygous gain* of one copy (+1/+1),
- one-sixth of the TR loci had *heterozygous loss* of one copy (0/-1),
- one-sixth of the TR loci had *homozygous loss* of one copy (-1/-1), and
- two-sixths of the TR loci had no change (0/0).

Two genotypes, paternal and maternal, were simulated using simuG (10) with 3,000,000 random SNPs and the TR changes in copy as vcf files. Paired-end Illumina reads were simulated using the ART read simulator (11) to represent the same profile as the NYGC data: HiSeq2500 error profile, read length of 150 bp, fragment length mean of 550 bp, and fragment length standard deviation of 150 bp. The fragment coverage was calculated to give read coverage of 30X, 50X, 70X, and 100X. In each case, half the reads were from the paternal genome and half from the maternal genome.

VNTRseek was run on each dataset at the different coverages, and the precision and recall were determined for alleles overall and within the detectable range of VNTRseek, *i.e.*, allele copy number of at least 1.9 (TRF limitation) and array size of 130 bp or less (allowing for flanking sequence of 10 bp on each side of the array). Results are presented in Figure S32 and Table S16 to Table S19. The runtimes are given in Table S20.

The precision, or positive predictive value, reduced slightly as the coverage increased, but in all cases was greater than 93%. The recall for the detectable alleles ranged from 41% to 44% for the -1 alleles, from 85% to 90% for the +1 alleles, and above 98% for the reference alleles. The very low level of detection for -1 alleles in the "all alleles" graph is due to the TRF limitation. Because a very high percentage of the reference TRs had copy number under 2.9 copies. loss of one copy put them outside the detectable range.

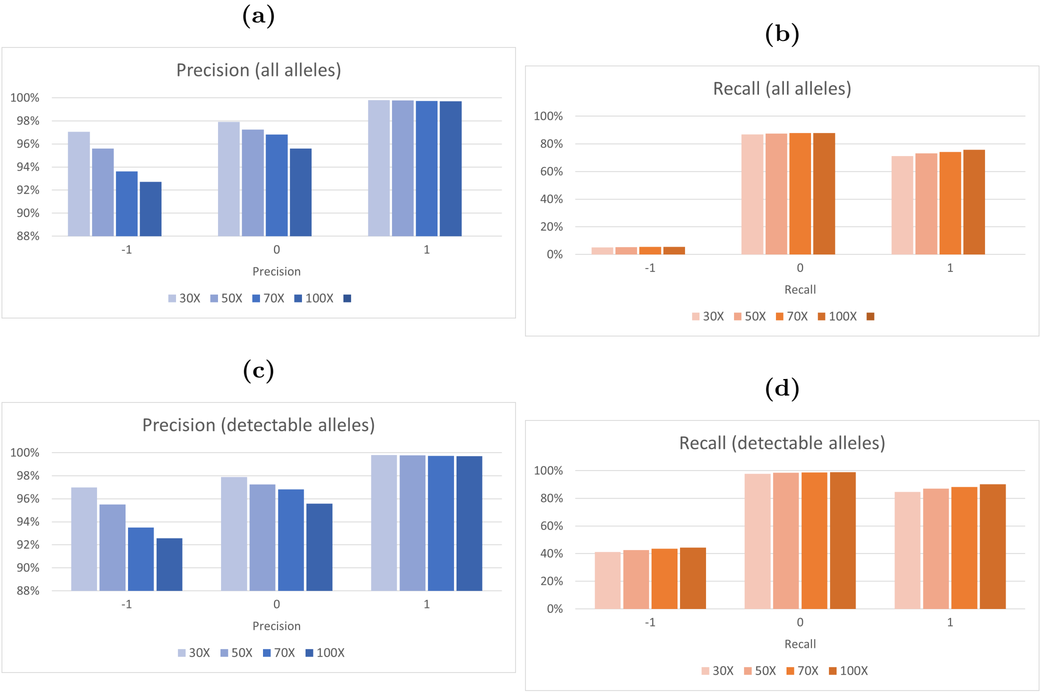


**Figure S32: VNTRseek performance by coverage.** Precision and recall for the entire collection of simulated TR alleles (sub-figures a and b) and the alleles in the detectable range of VNTRseek (sub-figures c and d) are shown. Precision, or positive predictive value, is the fraction of the predicted alleles that are correct (TP/(TP+FP)). Recall is the fraction correctly detected out of the total of that type (TP/(TP + FN)).

| **30X** | True -1 | True 0 | True 1 | Other |
|---|---|---|---|---|
| TP | 3,056 | 106,065 | 49,005 | 0 |
| FP | 95 | 2,275 | 96 | 106 |
| FN | 4,380 | 2,463 | 8,942 | 0 |
| Precision | 97% | 98% | 100% | NA |
| Recall | 41% | 98% | 85% | NA |

**Table S16: Detectable alleles confusion matrix of simulation at 30X coverage.**

| **50X** | True -1 | True 0 | True 1 | Other |
|---|---|---|---|---|
| TP | 3,161 | 106,833 | 50,405 | 0 |
| FP | 149 | 3,037 | 118 | 154 |
| FN | 4,275 | 1,695 | 7,542 | 0 |
| Precision | 95% | 97% | 100% | NA |
| Recall | 43% | 98% | 87% | NA |

**Table S17: Detectable alleles confusion matrix of simulation at 50X coverage.**

| **70X** | True -1 | True 0 | True 1 | Other |
|---|---|---|---|---|
| TP | 3,246 | 107,084 | 51,076 | 0 |
| FP | 226 | 3,524 | 143 | 184 |
| FN | 4,190 | 1,444 | 6,871 | 0 |
| Precision | 93% | 97% | 100% | NA |
| Recall | 44% | 99% | 88% | NA |

**Table S18: Detectable alleles confusion matrix of simulation at 70X coverage.**

| **100X** | True -1 | True 0 | True 1 | Other |
|---|---|---|---|---|
| TP | 3,297 | 107,294 | 52,225 | 0 |
| FP | 265 | 4,954 | 161 | 220 |
| FN | 4,139 | 1,234 | 5,722 | 0 |
| Precision | 93% | 96% | 100% | NA |
| Recall | 44% | 99% | 90% | NA |

**Table S19: Detectable alleles confusion matrix of simulation at 100X coverage.**

| Coverage | User time | System time | Wall clock time | CPU time | Max memory |
|---|---|---|---|---|---|
| 30X | 6:16:26:45 | 5:49:42 | 14:44:06 | 6:22:16:28 | 37G |
| 50X | 11:01:02:20 | 8:24:16 | 1:00:32:53 | 11:09:26:37 | 38G |
| 70X | 15:03:35:49 | 9:10:04 | 1:08:58:30 | 15:12:45:53 | 39G |
| 100X | 22:02:53:22 | 10:39:42 | 2:07:48:25 | 22:13:33:04 | 40G |

**Table S20: Run time of VNTRseek on simulated datasets.**

### References


1. Gelfand, Y., Hernandez, Y., Loving, J., and Benson, G. (2014) VNTRseek-a computational tool to detect tandem repeat variants in high-throughput sequencing data. *Nucleic Acids Research,* **42**(14), 8884–8894.

2. Nagraj, V., Magee, N. E., and Sheffield, N. C. (2018) LOLAweb: a containerized web server for interactive genomic locus overlap enrichment analysis. *Nucleic Acids Research,* **46**(W1), W194–W199.

3. Sheffield, N. C., Thurman, R. E., Song, L., Safi, A., Stamatoyannopoulos, J. A., Lenhard, B., Crawford, G. E., and Furey, T. S. (2013) Patterns of regulatory activity across diverse human cell types predict tissue identity, transcription factor binding, and long-range interactions. *Genome Research,* **23**(5), 777–788.

4. Bakhtiari, M., Shleizer-Burko, S., Gymrek, M., Bansal, V., and Bafna, V. (2018) Targeted genotyping of variable number tandem repeats with adVNTR. *Genome Research,* **28**(11), 1709–1719.

5. Gelfand, Y., Rodriguez, A., and Benson, G. (2007) TRDB—the tandem repeats database. *Nucleic Acids Research,* **35**(suppl_1), D80–D87.

6. Wenger, A. M., Peluso, P., Rowell, W. J., Chang, P.-C., Hall, R. J., Concepcion, G. T., Ebler, J., Fungtammasan, A., Kolesnikov, A., Olson, N. D., et al. (2019) Accurate circular consensus long-read sequencing improves variant detection and assembly of a human genome. *Nature Biotechnology,* **37**(10), 1155–1162.

7. Sudmant, P. H., Rausch, T., Gardner, E. J., Handsaker, R. E., Abyzov, A., Huddleston, J., Zhang, Y., Ye, K., Jun, G., Fritz, M. H.-Y., et al. (2015) An integrated map of structural variation in 2,504 human genomes. *Nature,* **526**(7571), 75–81.

8. Subramanian, A., Tamayo, P., Mootha, V. K., Mukherjee, S., Ebert, B. L., Gillette, M. A., Paulovich, A., Pomeroy, S. L., Golub, T. R., Lander, E. S., et al. (2005) Gene set enrichment analysis: a knowledge-based approach for interpreting genome-wide expression profiles. *Proceedings of the National Academy of Sciences,* **102**(43), 15545–15550.

9. Lee, D., Cheng, A., Lawlor, N., Bolisetty, M., and Ucar, D. (2018) Detection of correlated hidden factors from single cell transcriptomes using Iteratively Adjusted-SVA (IA-SVA). *Scientific Reports,* **8**(1), 1–13.

10. Yue, J.-X. and Liti, G. (2019) simuG: a general-purpose genome simulator. *Bioinformatics,* **35**(21), 4442–4444.

11. Huang, W., Li, L., Myers, J. R., and Marth, G. T. (2012) ART: a next-generation sequencing read simulator. *Bioinformatics,* **28**(4), 593–594.
